# Circulating tumor cells shed shearosome extracellular vesicles in capillary bifurcations that activate endothelial and immune cells

**DOI:** 10.1101/2024.04.17.589880

**Authors:** Angelos Vrynas, Salime Bazban-Shotorbani, Sara Arfan, Karishma Satia, Brian Cunningham, Gauhar Sagindykova, Mymuna Ashna, Aoyu Zhang, Diana Visan, Aisher Chen, Teige Matthews-Palmer, Mathew Carter, Fiona Blackhall, Kathryn L. Simpson, Caroline Dive, Paul Huang, Sam H. Au

## Abstract

Circulating tumor cells (CTCs) and their clusters are the drivers of metastasis, but we have an incomplete understanding of how they interact with capillary beds. Using microfluidic models mimicking human capillary bifurcations, we observed cell size- and bifurcation-dependent shedding of nuclei-free fragments by patient CTCs, CTC-derived explant cells and numerous cancer cell lines. Shedding reduced cell sizes up to 61%, facilitating their transit through bifurcations. We demonstrated that shed fragments were a novel subclass of large extracellular vesicles (LEVs), “shearosomes”, that require shear stress for their biogenesis and whose proteome was associated with immune-related pathways. Shearosomes exhibited functions characteristic of previously identified EVs including cell-directed internalization by endothelial and immune cells, and intercellular communication abilities such as disruption of endothelial barrier integrity, polarization of monocytes into M2 tumor-promoting macrophages and interactions between endothelial and immune cells. Cumulatively, these findings suggest that CTCs shed shearosomes in capillary beds that drive key processes involved in the formation of pre-metastatic niches.

## Introduction

Circulating tumor cells (CTCs) can travel through the vasculature to establish metastatic tumors in distant organs ^1–4^. CTCs are often entrapped in capillaries because of discrepancy in diameters between these vessels (5-10 μm) ^1^ and CTCs, which are often much larger^4^. However, entrapment of CTCs alone is not sufficient for successful metastasis. Instead, the microenvironment of distant organs needs to be modified to make these sites more conducive to the migration, survival, and proliferation of disseminated tumor cells ^2–4^. This process, known as the formation of premetastatic niches (PMN), is characterized by the disruption of blood vessel barrier integrity that promotes tumor cell extravasation, the activation of endothelial cells, polarization and/or recruitment of stromal cells including fibroblasts, macrophages and other immune cells ^5^. It is currently accepted that primary tumors are responsible for the formation of PMN through the secretion of various signaling molecules and vesicles ^5,6^. However, we hypothesize that there exists another mechanism whereby CTCs can prime the formation of PMN after becoming entrapped in capillaries.

In this work, we characterized the shedding of large cytoplasmic fragments by patient CTCs and CTC-derived cells entrapped in microfluidic models of human capillary bifurcations. This shedding phenomenon was previously observed in vivo, was attributed to shear stress, and led to the internalization of fragments by monocytes, macrophages, neutrophils and dendritic cells ^7^. What we do not understand are the identity of these fragments, the mechanism and frequency of fragment biogenesis, and how these fragments drive key processes in metastasis such as PMN formation. Our analysis indicates that shedding is a biomechanical process that generates a novel class of large extracellular vesicles (LEVs) from CTCs. LEVs were readily internalized by endothelial cells, monocytes and M1 macrophages and can drive key processes in the formation of the PMN such as disruption of endothelial barrier integrity, endothelial cell activation, polarization of monocytes and M1 macrophages into M2 tumor-promoting macrophages, and secondary co-interactions between endothelial cells and immune cells. Overall, we propose that this mechanism of shedding allows CTCs to influence their local microenvironment in ways that improve their probabilities of metastatic success.

### Capillary bifurcations promote cell shedding of large extracellular vesicles

To mimic the diversity of geometries, present in human microvasculature (i.e. capillaries and small arterioles), we engineered 13 distinct variants of microfluidic constricted bifurcation devices alongside constricted non-bifurcated (linear) (7 μm) and non-constricted (20 μm) control devices (**Fig. 1A and Supplementary Fig. 1 and Supplementary Table 1**) alongside computational fluid dynamic simulations of each variant (**Supplementary Fig. 2,3**) that show fluid shear stress profiles of constricted bifurcations within the range of values reported for microvasculature^8^. These devices allowed us to systematically explore via live imaging (**Fig. 1B**) the influence of physiological levels of pressure (10-30 cm H_2_O) on the behavior of primary CTCs isolated from two small cell lung carcinoma (SCLC) patients, SCLC patient CTC-derived xenograft (CDX) cells, six cancer cell lines (MDA-MB 231, B16F10, Me290, LNCaP, PDAC, OVSAHO), primary cancer-associated fibroblasts, THP-1 monocytes and THP-1-derived M0 macrophages at capillary-sized bifurcations. Upon introduction of cells into microfluidic bifurcation devices, we observed the formation of large (1.17-11.32 μm, mean: 4.9 ± 1.9 μm) Calcein-AM-positive, Hoechst-negative cellular fragments in real-time transit of all aforementioned cell types at capillary bifurcations within seconds (**Fig. 1C-F and Supplementary Fig. 4A-F, and Supplementary Movies S1 to S4**). We later demonstrate that these fragments matched all the MISEV2018 classification requirements of extracellular vesicles ^9^ (described below), and we henceforth call these fragments “large extracellular vesicles” (LEVs). MDA-MB 231 cells shed at various constricted bifurcation geometries with a mean frequency of 28.5 ± 19%, a significantly higher rate than in linear geometries (3.6 ± 5%) (**Fig. 1G**) or static conditions (**Supplementary Fig. 4G**). Furthermore, centrifugal forces on cells at comparable levels did not lead to enhanced cell shedding (**Supplementary Fig. 4H**). Among the tested geometrical variations in the constricted bifurcation geometries, only smaller daughter bifurcation diameters (**Fig. 1H**) and narrower angles between daughter bifurcations (**Supplementary Fig. 4I-L**) significantly increased the frequency of cell shedding (p<0.05). As expected, reduction of the pressure across capillary bifurcations led to a decrease in the frequency of cell shedding (**Fig. 1I**). Most cells remained viable after shedding LEVs (described below), including immune cells such as monocytes (**Supplementary Fig. 4E**) and the proliferation rate of MDA-MB-231 cells post transit remained statistically indistinguishable from control cells (**Supplementary Fig. 4M,N**). Additionally, clusters of tumor cells also shed LEVs (**Supplementary Fig. 5A and Supplementary Movie S5**), at higher frequencies than single cells (83.8 ± 6% vs 52.2 ± 3%, P<0.05) (**Supplementary Fig. 5B**). Clusters that contained three or more cells had higher probability of experiencing at least one shedding event in comparison to doublets (**Supplementary Fig. 5C**), and clusters released more LEVs than singlets (**Supplementary Fig. 5D,E and Supplementary Movie S5**), which is likely a reflection of the greater number of cells within clusters.

**Fig. 1.**
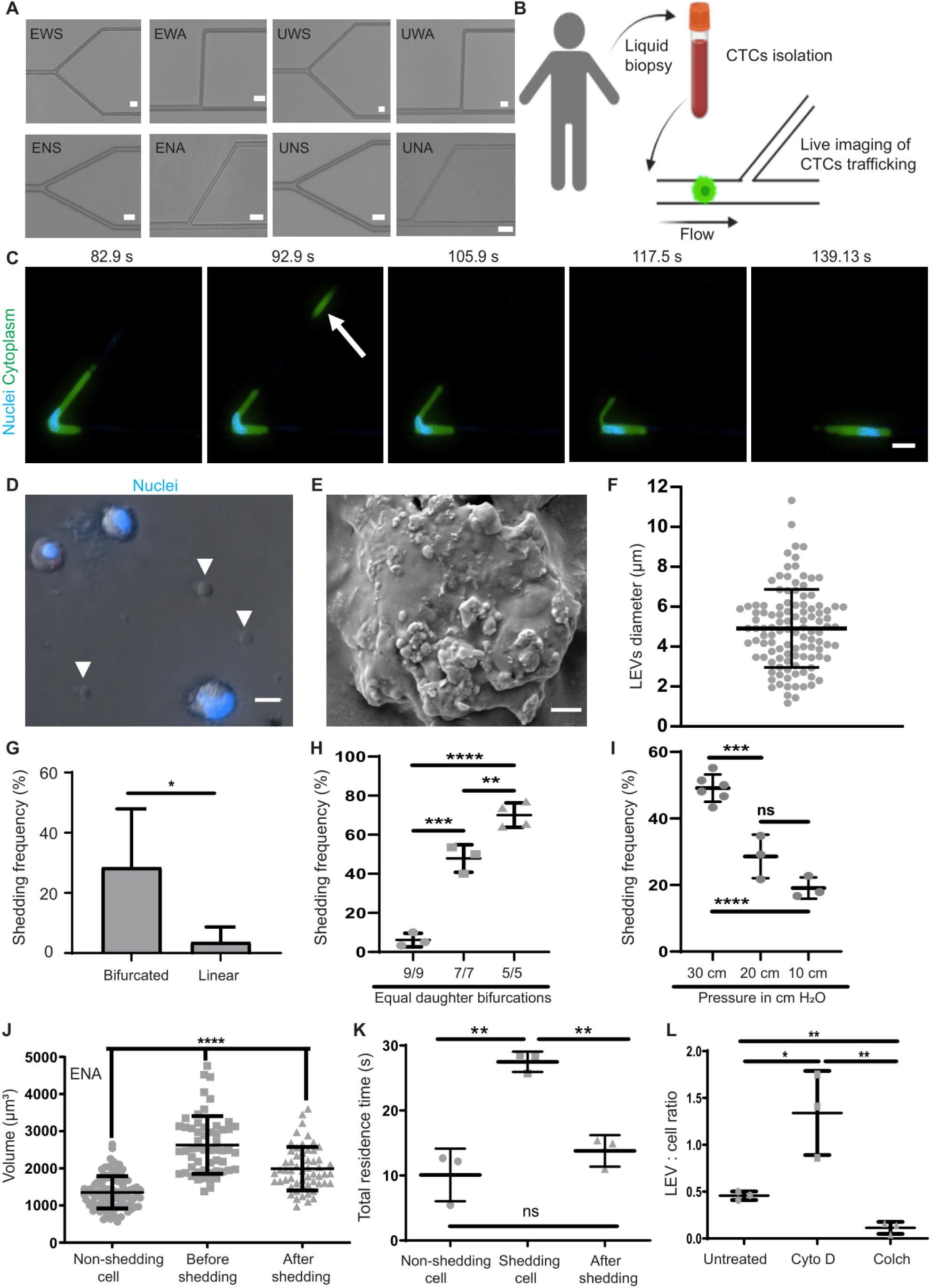
Biogenesis mechanisms of LEVs by tumor cells in capillary geometries & their fate post shedding. (**A**) Images of 8 bifurcation variants (E:Equal, U:Unequal, W:Wide, N:Narrow, S:Symmetry, A:Asymmetry). Scale bar 20 μm. (**B**) Experimental workflow of live imaging. Created with Biorender.com. (**C**) Timelapse of a MDA-MB231 breast cancer cell shedding LEV (white arrow). Cytoplasm was stained with CMFDA cell tracker (green) and nuclei with Hoechst-33342 (blue). Scale bar 20 μm. (**D**) Hoechst-positive MDA-MB231 cells and Hoechst-negative LEVs. Scale bar 10 μm. (**E**) Scanning electron microscopy image of a LEV. Scale bar 2 μm. (**F**) Diameter (μm) of LEVs (n=111 LEVs). (**G**) Shedding frequency of MDA-MB231 cells in various bifurcation variants. (**H**) Shedding frequency of MDA-MB231 cells in ENA_9/9 (n=3), ENA_7/7 (n=3) & ENA_5/5 (n=4). (**I**) Shedding frequency of MDA-MB231 cells under various pressures (n at least 3) (**J**) Volume (μm^3^) of MDA-MB231 non-shedding cells (n=95), or ^10^shedding cells (n=56). (**K**) Mean total residence time (s) of non-shedding MDA-MB231 cells, shedding cells and after shedding (n=3). (**L**) LEV:cell ratio of untreated, Cytochalasin-D (Cyto-D) treated and Colchicine (Colch)-treated MDA-MB231 cells, enumerated post transit from bifurcation UNA_5/9 (n=3).

We then investigated if cytoplasmic fragments that matched the morphological characteristics of LEVs were present in the blood of patients. We first isolated CTCs from blood samples of two SCLC patients (Ethical Committee reference 07/H1014/96) by negative affinity selection and then stained the effluent to identify cellular fragments. We identified 64 and 356 CMFDA-positive, Hoechst-negative cellular fragments that matched the morphology of LEVs in these blood samples (**Supplementary Fig. 6A,B**). Both patients presented higher counts of cellular fragments compared to CTCs (**Supplementary Fig. 6B**), and fragments ranged in size from 1.03 to 8.73 μm (mean: 2.4 ± 1.4 μm) (**Supplementary Fig. 6C**).

### LEV biogenesis is a cell size-dependent and a volumetric subtractive process that facilitates CTC transit

We first quantified the total volume of cells before encountering bifurcations in our microfluidic devices. We found that the mean volume of cells that shed LEVs was significantly larger (∼1.9 times) than their non-shedding counterparts (**Supplementary Movie S6**) during transit in bifurcation variant ENA (2628 ± 8×10^2^ μm^3^ vs 1354 ± 4×10^2^ μm^3^, P<0.0001) (**Fig. 1J and Supplementary Fig. 7A,B**) and 1.7-2.4 times larger for the seven other tested bifurcation geometries (**Supplementary Fig. 7C-F**). Overall, cell size positively correlated with shedding frequency (r^2^ = 0.61) (**Supplementary Fig. 8A**). Smaller cells experienced minimal shedding of LEVs in linear models (**Supplementary Fig. 8B**) but were much more capable when transiting through bifurcations (**Supplementary Fig. 7C-F and Supplementary Fig. 8C-E**). This size dependency is also supported by our experiments with immune cells, where blood-resident monocytes shed with significantly lower frequency than larger macrophages (**Supplementary Fig. 4D)**, which is expected given the larger sizes of macrophages^10^. Shedding cells had a mean ∼25% reduction in tumor cell volume post-shedding during transit in bifurcation variant ENA (2628 ± 8×10^23^ vs 1991 ± 6×10^2^ μm^3^, P<0.0001) (**Fig. 1J**), which varied from 19 to 22.7% for other tested bifurcation variants (**Supplementary Fig. 7D-F**). We also found that bifurcations with equal effective diameter daughter bifurcations caused significantly greater volumetric losses in cells than their unequal counterparts (**Supplementary Fig. 9**).

We then recorded how long shedding vs. non-shedding cells remained within capillary bifurcations (residence time). Non-shedding cells exhibited a mean residence time of 10 ± 4 s (**Fig. 1K**). Cells that eventually shed remained in devices ∼3 times longer (**Fig. 1K**). Interestingly, after shedding LEVs, the mean residence time of cells was dramatically reduced to levels statistically indistinguishable from the residence time of non-shedding cells (**Fig. 1K**). We postulate that the reductions in cell size imposed by shedding, reduced the hydrodynamic resistance of these cells, leading to commensurate reductions in residence times. This suggests that shedding of LEVs facilitates the transit of relatively large but not small CTCs through capillary bifurcations.

### F-actin modulates LEV biogenesis and post-shedding cell viability

After exploring the importance of biophysical parameters such as capillary bifurcation geometries and cell size on LEV biogenesis, we wondered whether the actin cytoskeleton was involved in shedding. To this end, we pre-treated cells with Cytochalasin-D (Cyto-D) or Colchicine (Colch) to inhibit or promote F-actin polymerization, respectively (**Supplementary Fig. 10A**) at concentrations that preserved cell viability (**Supplementary Fig. 10B,C**). Cyto-D significantly increased the probability of LEVs biogenesis compared to untreated conditions during single cell transit through capillary bifurcations (**Fig. 1L and Supplementary Fig. 11A**) or non-constricted geometries (**Supplementary Fig. 11B**). On the other hand, Colch had the opposite effect, reducing LEVs biogenesis in a statistically significant manner compared both to untreated and Cyto-D treated cells in all tested conditions (**Fig. 1L and Supplementary Fig. 11B**). Cyto-D treatment did not significantly increase the shedding frequency from tumor cell clusters in capillary bifurcations (**Supplementary Fig. 12A,B**) or non-constricted geometries (**Supplementary Fig. 12C**). We believe that may be because clusters already exhibited high shedding frequency even without chemical treatment (**Supplementary Fig. 5B**). Interestingly, Cyto-D treatment increased LEVs biogenesis from single cells but not clusters under shear (**Supplementary Fig. 12D**).

We then quantified cell death 3 hours after cell transit in capillary geometries. A small fraction of untreated cells (∼15%) were stained dead after transit in capillary bifurcations (**Supplementary Fig. 13A-B**). Interestingly, Cyto-D-pre-treated cells died at statistically significant higher rates post-transit through bifurcations (**Supplementary Fig. 13A-B**) and non-constricted devices (**Supplementary Fig. 13C-middle**), but remained unaltered in constricted linear geometries (**Supplementary Fig. 13C-left**), static conditions (**Supplementary Fig. 13C-right**) or under Colch treatment (**Supplementary Fig. 13B**). Cell viability was further assessed using live cell fluorescent imaging. Lysis of cells during transit in capillary bifurcations was observed (**Supplementary Fig. 13D and Supplementary Movie S7**) and Cyto-D induced higher rates of cell lysis at capillary bifurcations (**Supplementary Fig. 13E**). We then went on to confirm that the higher rates of cell lysis in Cyto-D-pre-treated cells occurred in shedding cells (**Supplementary Fig. 13F**), and thus Cyto-D-pre-treated cells had far greater probabilities of cell lysis post shedding (**Supplementary Fig. 13G**). Altogether, F-actin modulation influences both LEV biogenesis and viability when cells are subjected to shear and capillary bifurcation forces.

### Cytoplasmic fragments meet all established criteria of large extracellular vesicles

We developed purification protocols using centrifugation or filtration that successfully separated cells and their shed LEVs to purities above 99.9%, validated by flow cytometry (**Fig. 2A and Supplementary Fig. 14A-D**) and microscopy (**Fig. 2B and Supplementary Fig. 14E,F**). We then evaluated purified shear-derived LEVs using the Minimal Information for Studies of Extracellular Vesicles (MISEV 2018) criteria ^9^. We used immunocytochemistry to verify that LEVs were positive for both cytosolic (CK18, HSP90) and transmembrane (CD81) proteins, similar to their cells of origin, using flow cytometry (**Fig. 2C,D, and Supplementary Fig. 15**), microscopy (**Supplementary Fig. 16A-D**) and western blot (**Supplementary Fig. 16E**). All We then used three distinct imaging modalities, fluorescent light microscopy (**Fig. 1D**), scanning electron microscopy (**Fig. 1E**), and cryogenic transmission electron microscopy (cryoEM) (**Supplementary Fig. 16F**) to visualize LEVs. Furthermore, crytoEM confirmed that vesicles contained cytosol bound by a lipid bilayer with phosphate leaflet spacing of 4nm (**Supplementary Fig. 16F,G**). Densities typical of membrane-tethered proteins were abundant on the exterior side of the membrane (**Supplementary Fig. 16F,G**).

**Fig. 2.**
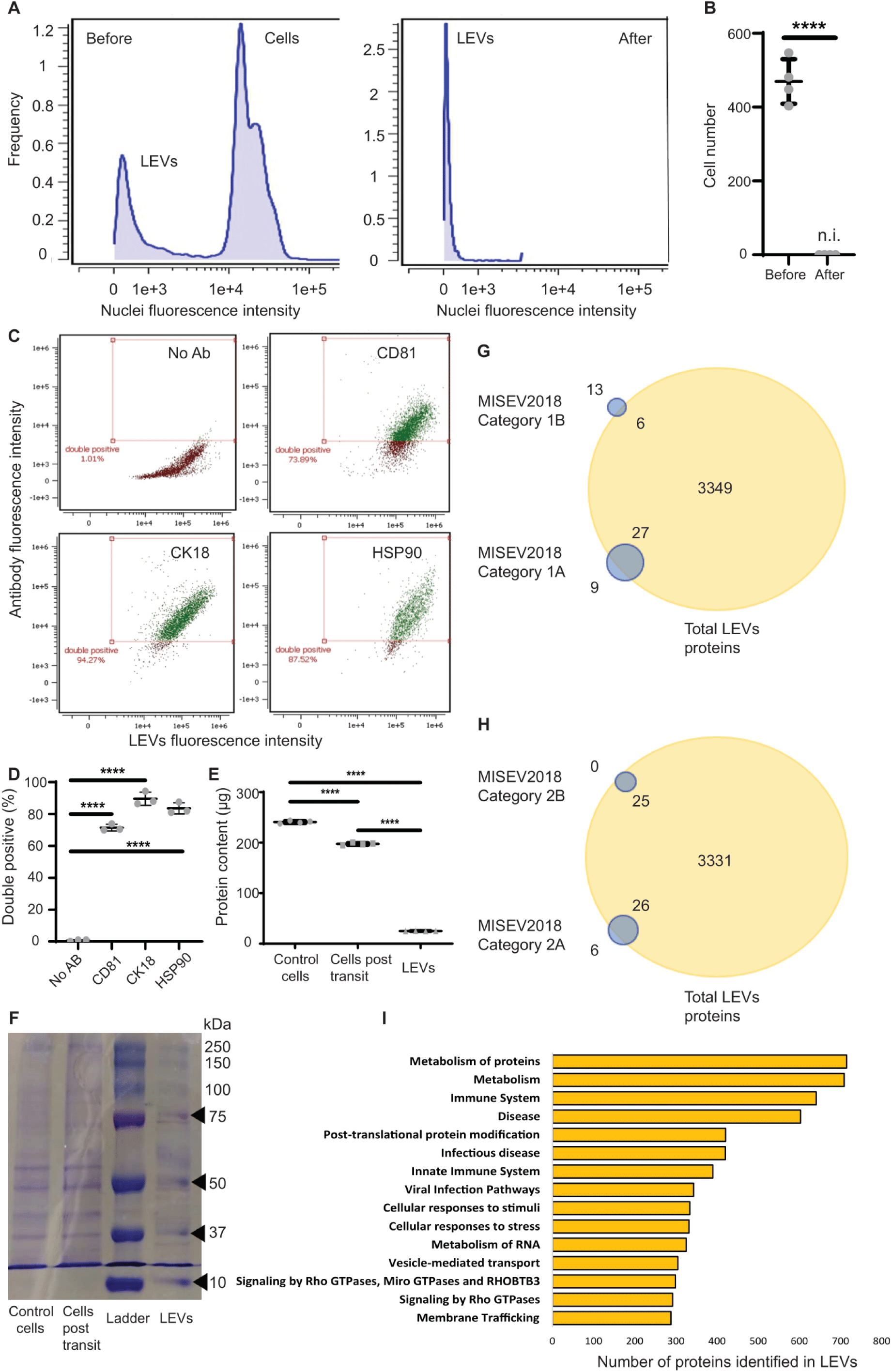
LEVs purification and interrogation of protein content & correlation to downstream functionalities. (**A**) Histogram of MDA-MB 231 cells and LEVs before and after centrifugation analyzed via flow cytometry. (**B**) Quantification of MDA-MB 231 cell number before and after centrifugation (n=4). (**C-D**) Flow cytometry analysis of LEV proteins. (**C**) Scatter plots of events double positives for CMTPX red cell tracker pre-stained LEVs and CD81 (top right-green), CK18 (bottom left-green) and HSP90 (bottom right-green). Non-antibody stained (No Ab) LEVs (top left-red) were used as control (n=3). & quantification (**D**) Quantification of data in (C). (**E**) Total protein content (μg) of control MDA-MB 231 cells, cells post transit and LEVs (n=4). (**F**) Protein blot for control MDA-MB 231 cells, cells post transit and LEVs against a ladder. (**G**) MISEV2018 category 1 transmembrane and GPI-anchored proteins identified in LEVs through data independent acquisition (DIA) mass spectrometry (MS)-based proteomics. (**H**) MISEV2018 category 2 cytosolic proteins identified in LEVs through DIA MS-based proteomics. (**I**) Top 15 Reactome gene sets represented in LEVs proteome. Gene sets are ranked based on number of overlapping LEVs proteins.

### LEVs are rich in proteins relevant to immune responses

We then sought to investigate the content of LEVs. Parental cells lost approximately 18% of their total protein content after transit through capillary bifurcations (**Fig. 2E and Supplementary Fig. 17A,B**). About 10% of the original total protein content in cells was detected in recovered LEVs (**Fig. 2E and Supplementary Fig. 17A,B**), suggesting that some proteins may be shed to the extracellular space during shedding. Further protein analyses revealed the overlap of proteins between LEVs and cells at various molecular weight bands (**Fig. 2F**). Interestingly, similar to other EVs, we found RNA present in LEVs (**Supplementary Fig. 17C**), including several mRNA transcripts (**Supplementary Fig. 17D**).

We then undertook mass spectrometry based proteomic profiling to determine protein composition of LEVs. Our analysis defined 3,382 proteins within the LEVs, representing 60% of all proteins identified in parental cells from which LEVs were derived (**see Supplementary Table 2 and ProteomeXchange – dataset identifier: PXD050444 for the full list of proteins**). When compared with the MISEV2018 panel of proteins for categories 1 and 2 ^9^, 33 of 55 transmembrane proteins from MISEV2018 categories 1A and 1B (27 non-tissue specific and 6 cell/tissue specific) ^9^ (**Fig. 2G and Supplementary Table 3**) and 51 of 57 cytosolic proteins from MISEV2018 categories 2A and 2B ^9^ (**Fig. 2H and Supplementary Table 4**) were identified in LEVs including CK18, HSP90, and CD81, previously identified by immunohistochemistry (Supplementary Table 2). Interrogation of Reactome database of the top 15 gene sets which are composed of LEV proteins, identified a range of different biological pathways including multiple immune-related pathways (immune system, infectious disease, innate immune system, viral infection pathways), regulation of vesicle transport and trafficking (membrane trafficking, signaling by Rho GTPases and vesicle-mediated transport) as well as protein metabolism (**Fig. 2I**). Interestingly, we identified proteins such as transforming growth factor beta (TGFβ1) and interleukin-6 (IL-6) in LEVs (**Supplementary Table 2**), that are mutually responsible for activation of endothelial cells and monocytes ^11–14^. Our proteomic analysis also identified 66 distinct proteins in the LEVs, annotated for protein secretion function (Supplementary Table 5), suggesting that LEVs may retain secretory capabilities from parental cells. These prompted us to conduct additional experiments exploring cell-LEV interactions.

### Endothelial and immune cells actively internalize LEVs

We first explored if LEVs could be actively internalized by cells, similar to previously identified smaller extracellular vesicles ^15,16^. We co-cultured LEVs purified from MBA-MB-231 breast cancer cells with human umbilical vein endothelial cells (HUVECs), THP1 human monocytes or THP-1 differentiated M1 macrophages. We verified the internalization of LEVs in all tested cell types via co-localization with cells’ cytoplasm in Z-stack slices (**Fig. 3A-C**) and quantified their internalization via flow cytometry (**Fig. 3D-F, and Supplementary Fig. 18**). We also observed that while some LEVs external to cells exhibited chaotic movements consistent with Brownian motion, those co-localized with cells did not (**Supplementary Movie S8**), further supporting the notion that LEVs were internalized. We also examined if monocytes and M1 macrophages internalized LEVs under flow. Monocytes internalized LEVs under flow to similar levels to static co-culture conditions (**Fig. 3E and Supplementary Fig. 19A**), whereas M1 macrophages internalized LEVs under flow to a lesser extent (**Fig. 3F and Supplementary Fig. 19B**). The mean diameter of LEVs internalized by HUVECs was 2.8 ± 1.2 μm (**Supplementary Fig. 20A**), smaller than the mean diameter of all shed LEVs (4.9 ± 1.9 μm) (**Fig. 1F**).

**Fig. 3.**
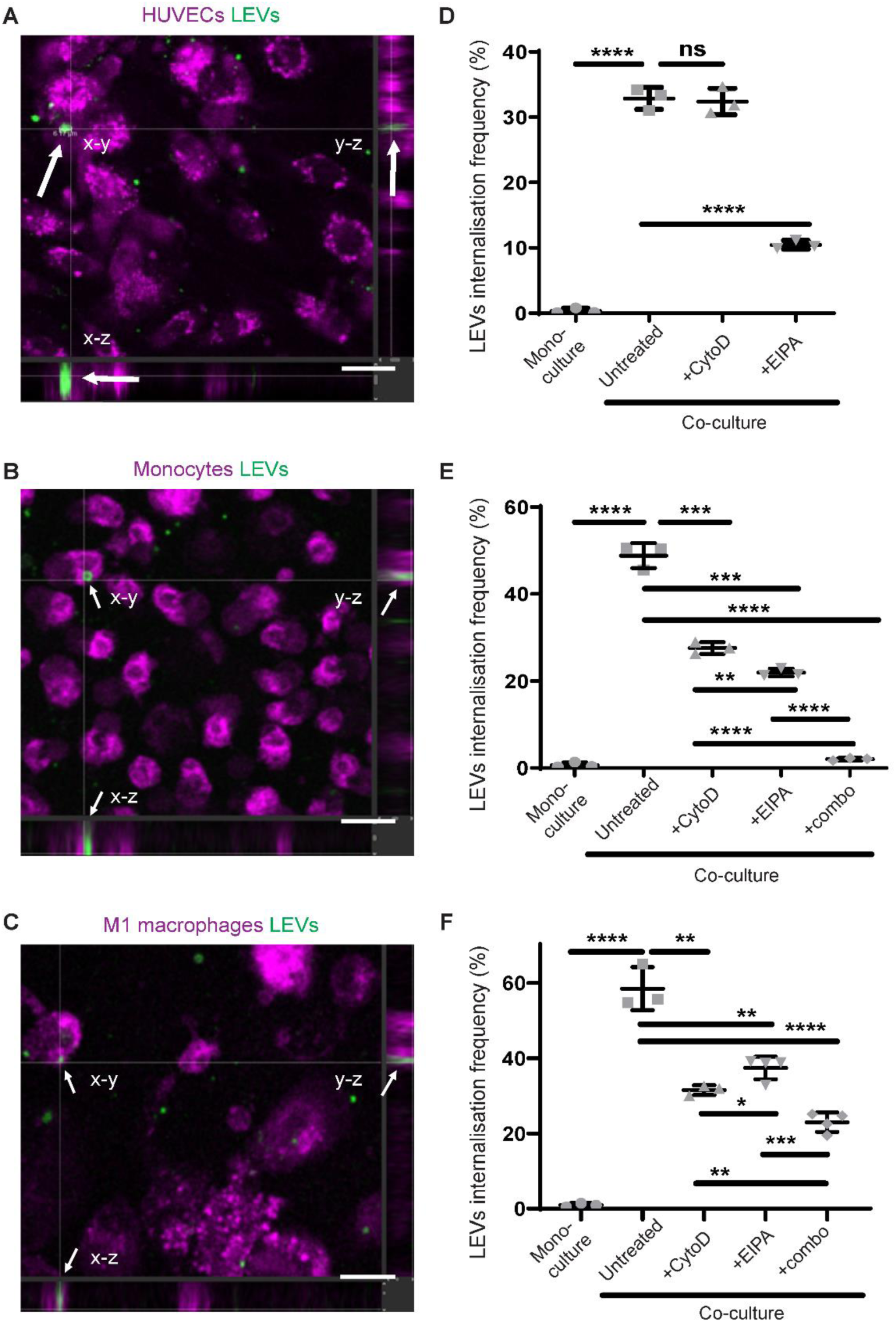
HUVECs, THP-1 monocytes and THP-1 differentiated M1 macrophages internalize LEVs. (**A-C**) Orthogonal cross sectioning of multifluorescent confocal images validating the internalization of LEVs by HUVECs (**A**), monocytes (**B**) and M1 macrophages (**C**). Cells were stained with CMTPX cell tracker (shown as magenta) and LEVs with CMFDA cell tracker (green). Scale bar 20 μm. Arrows indicate the internalized LEV in all planes. (**D-F**) Quantification of LEVs internalization frequency (double positive events) by HUVECs (**D**), monocytes (**E**) and M1 macrophages (**F**) (n=3 at least).

We then sought to identify the mechanisms responsible for LEV internalization. The two major mechanisms responsible for cellular internalization of large (>1 µm) particles are phagocytosis and micropinocytosis ^17^. Cytochalasin D, a phagocytosis inhibitor ^18^, inhibited the mean internalization frequency of LEVs by monocytes by 43% (P<0.0005) (**Fig. 3E**) and M1 macrophages by 46% (P<0.005) (**Fig. 3F**), but not HUVECs (p>0.05) (**Fig. 3D**). On the other hand, 5-(N-Ethyl-N-isopropyl)-Amiloride (EIPA), a macropinocytosis inhibitor ^18^, inhibited the mean internalization frequency of LEVS by monocytes by 55% (P<0.0005) (**Fig. 3E**), M1 macrophages by 36% (P<0.005) (**Fig. 3F**) and HUVECs by 68% (P<0.0001) (**Fig. 3D**). Interestingly, the combination of both Cyto-D and EIPA fully abrogated LEV internalization by monocytes (**Fig. 3E**), but not entirely in M1 macrophages (**Fig. 3F**). We quantified no internalization of fluorescently labelled beads by any cell type (**Supplementary Fig. 20B-D**).

### LEVs polarize monocytes and M1 macrophages towards CD206^+^ M2 macrophage lineages

The polarization of monocytes and macrophage phenotypes is an important contributor to PMN formation ^6,19^. We found that purified and washed LEVs (details in methods) co-cultured with THP-1 monocytes promoted monocyte adhesion (**Fig. 4A and Supplementary Fig. 21A**), led to the acquisition of stretched morphologies (**Supplementary Fig. 21B**) and enhanced proliferation (**Supplementary Fig. 21C**). M2 macrophage marker CD206 ^20^ had higher expression levels in LEV-treated monocytes (**Fig. 4B-E**), while M1 macrophage marker TNF-α^20^ was absent (**Fig. 4B**). To investigate if proteins secreted by LEVs were responsible for monocyte differentiation, we incubated LEVs in cell culture media for 24 hours and collected their conditioned media (CM). Monocytes treated with CM from LEVs, exhibited significantly lower mean fluorescence intensity of CD206 (**Fig. 4C,D**), and fewer CD206+ cells (**Fig. 4E,F**) than LEV-treated counterparts. We hypothesized that intravesicular TGF-β may have contributed to the polarization of monocytes towards M2 macrophages. Silencing siRNA in MDA-MB-231 prior to LEV biogenesis led to an approximately 20% decrease in the number of monocytes that exhibited polarization markers towards M2 states (**Fig. 4F**). We investigated the degree of M2 macrophage polarization by exploring gene expression levels. Indeed, LEV treatment caused significant upregulation of M2 macrophage related genes such as CD206 ^20,21^ and fibronectin ^20^ both in LEV-treated monocytes (**Fig. 4G**) and M1 macrophages (**Fig. 4H**). However, arginase-1, another M2 marker ^21^, was only statistically increased in LEV-treated M1 macrophages (**Fig. 4G,H**). Other genes such as interleukin-6 and CXCL10, whose association to M1 or M2 polarization is contentious, were elevated in some LEV-treated conditions but not to statistically significant levels (**Supplementary Fig. 22**).

**Fig. 4.**
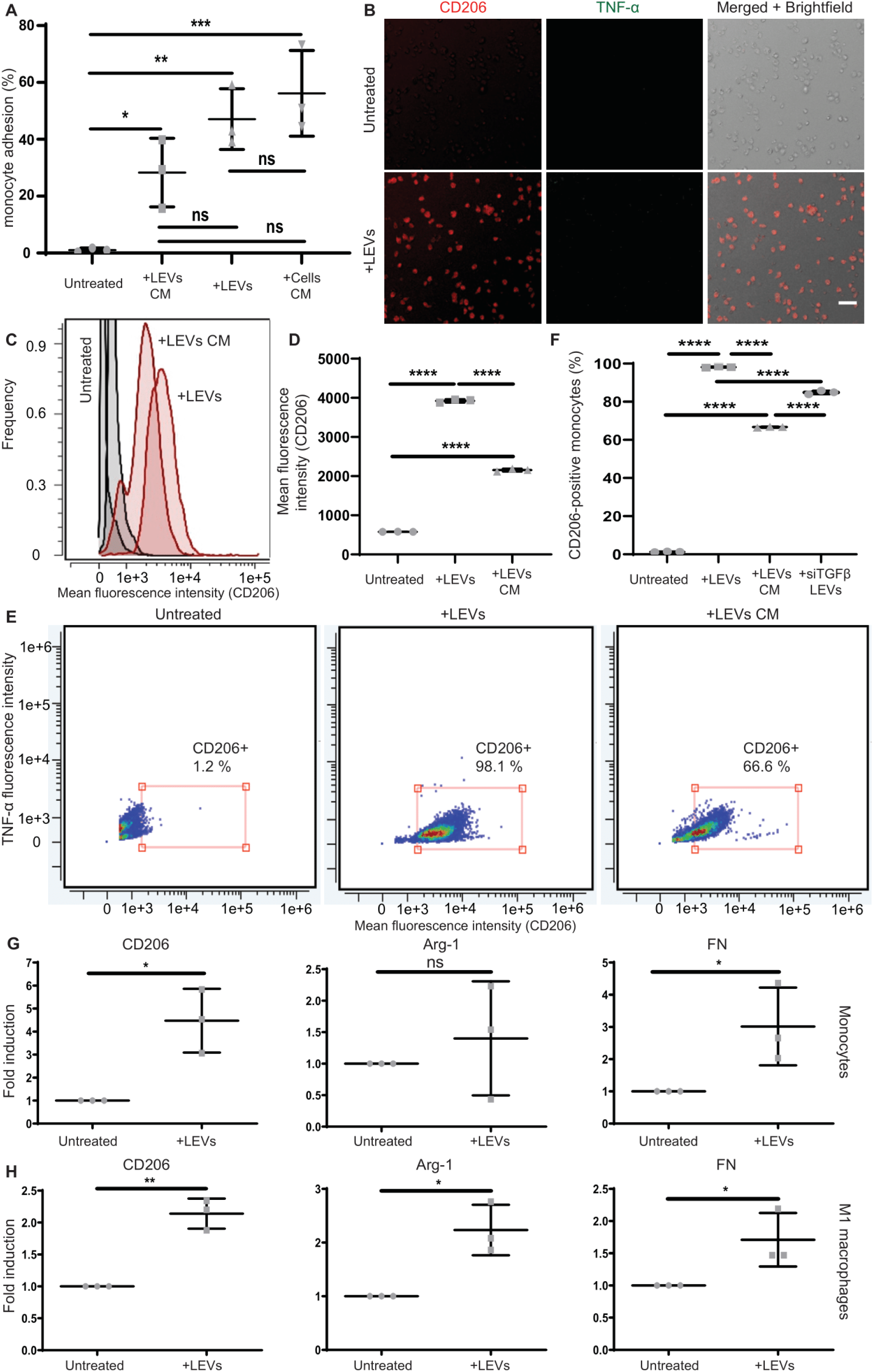
LEVs polarize monocytes and M1 macrophages to CD206^+^ M2 tumor-promoting macrophages. (**A**) Percentage of monocytes that remained adhered. Monocytes were co-cultured with LEVs or cultured with CM from LEVs or CM from MDA-MB231 cells. Untreated monocytes served as negative control. (n=3). (**B**) Multifluorescent images of untreated or LEV-treated monocytes stained for CD206 (red) and TNF-a (green). Scale bar 50 μm. (**C-D**) Histogram (**C**) & quantification (**D**) of CD206 mean fluorescence intensity for untreated monocytes, LEV-treated monocytes or monocytes treated with LEVs’ CM for 30hr, analyzed via flow cytometry (n=3). (**E-F**) Dot plots (**E**) & quantification (**F**) of CD206-positive events for untreated monocytes (left), LEV-treated monocytes (middle), monocytes treated with LEVs’ CM (right) and monocytes treated with LEVs from siTGFβ-treated MDA-MB-231 cells for 30hr, analyzed via flow cytometry (n=3). (**G-H**) Reverse transcription quantitative polymerase chain reaction of untreated or LEV-treated cells for genes CD206 (left), Arginase-1 (Arg-1) (middle) and fibronectin (FN) (right), for monocytes (**G**) and M1 macrophages (**H**) (n=3).

### LEVs disrupt the endothelial barrier integrity and activate endothelial cells

The disruption of the vascular endothelium is an important component of PMN formation ^5,6^. LEVs added to confluent monolayers of human endothelial cells caused morphological changes (**Supplementary Fig. 23A**) and reduced both mRNA (**Fig. 5A**) and protein expression levels of intercellular adhesions i.e. V-E-cadherin versus controls (**Fig. 5B-C and Supplementary Fig. 23B**). Furthermore, LEVs disrupted the continuity of endothelial monolayers, significantly increasing the cell-free area (**Fig. 5D and Supplementary Fig. 23C**). Similar to the above experiments, LEVs derived from siTGFβ MDA-MB231 cells did not lead to reductions in HUVEC cell-free area, suggesting that intravesicular TGFβ appears to damage endothelial integrity (**Supplementary Fig. 23D**). To test whether proteins secreted from LEVs affected endothelial cells, we cultured purified LEVs for 24hr and collected their CM. We found similar disruptions in VE-cad junctions when LEV CM was applied on HUVECs (**Fig. 5B-bottom left**). To investigate if these findings could be recapitulated in more physiological microenvironment, we generated 3D endothelial networks in hydrogel-laden microfluidic devices (**Fig. 5E**). Perfusion of the microfluidic 3D endothelial networks with LEVs (generated as described above), led to similar disruptions to endothelial cells (**Fig. 5F and Supplementary Fig. 23E**). Treatment with LEVs disrupted VE-cad localization within the 3D endothelial networks, evident by the 60% mean reduction of mean VE-cad fluorescence intensity (**Fig. 5G**). Moreover, analyses of the untreated and LEV-treated networks revealed that LEV treatment induced an increase in the cell-free area (**Fig. 5H**), similarly to 2D HUVEC monolayers (**Fig. 5D**).

**Fig. 5.**
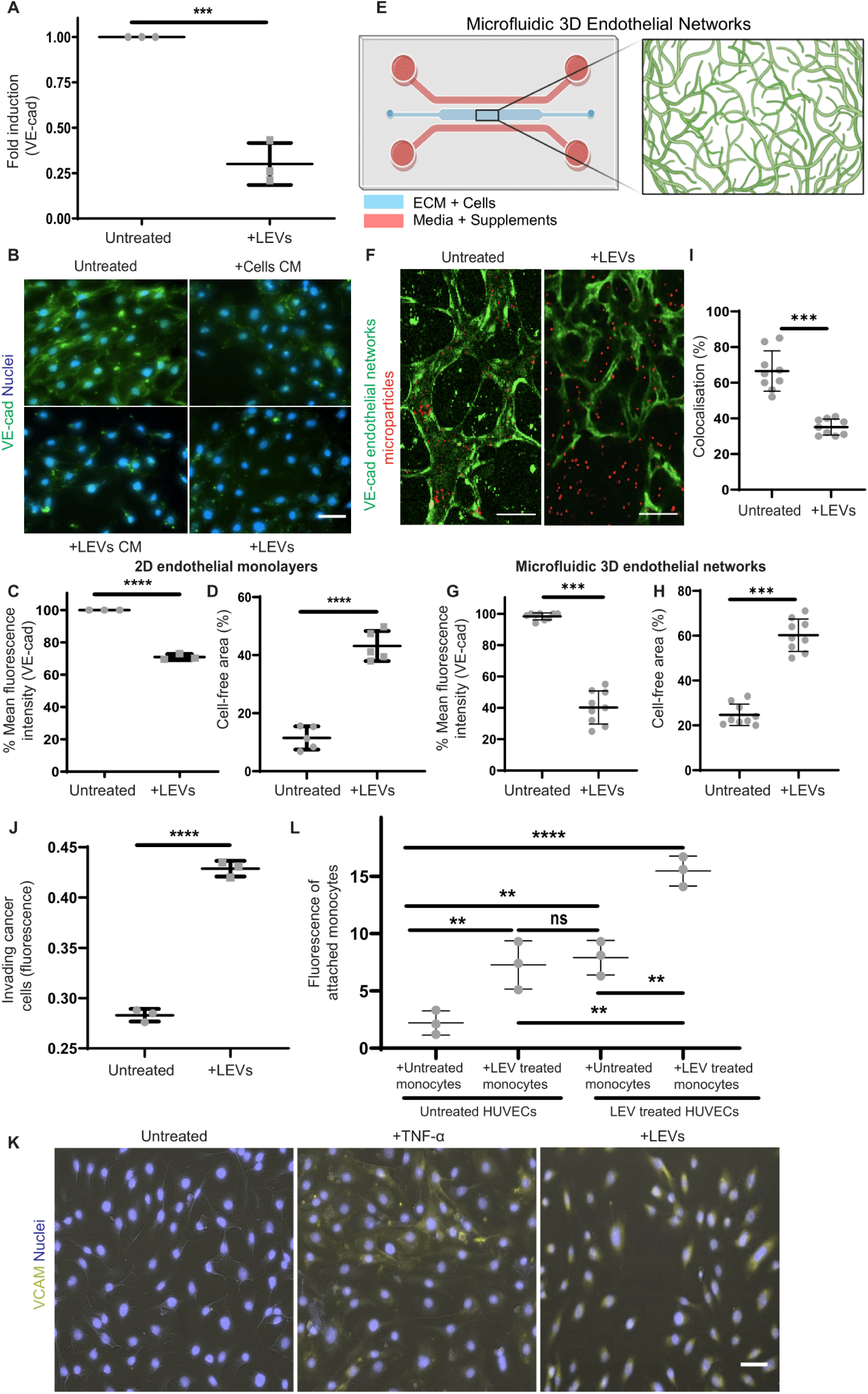
LEVs disrupt and activate the endothelium. (**A**) Reverse transcription polymerase chain reaction of untreated or LEV-treated HUVECs (30hr) for VE-cad gene (n=3). (**B**) Vascular endothelial cadherin (VE-cad) (green) staining of LEV-pre-treated HUVECs (bottom right), LEVs CM-pre-treated HUVECs (bottom left) or MDA-MB 231 cell CM-pre-treated HUVECs (top right) and untreated HUVECs (top left) in 2D. Nuclei was stained with Hoechst-33342 (blue). Scale bar 50 μm. (**C**) Quantification of VE-cad % mean fluorescence intensity of HUVECs in 2D, analyzed via flow cytometry (n=3). (**D**) Quantification of cell-free area from untreated or LEV-treated 2D HUVEC monolayers (n=5 areas). (**E**) Schematic of 3D vascular network formation in a microfluidic chip. Created with Biorender. (**F**) Fluorescent staining of microfluidic 3D endothelial networks (green) perfused by dextran 1 μm microparticles (red) in untreated or LEV-treated microfluidic 3D endothelial networks. Scale bar: 100 μm. (**G**) Quantification of VE-cad % mean fluorescence intensity of untreated and LEV-treated 3D endothelial networks (n=3). (**H**) Quantification of % cell-free area of untreated and LEV-treated 3D endothelial networks (n=3). (**I**) Quantification of % of 3D endothelial networks and microparticles colocalization (n=3). (**J**) Fluorescence of invading MDA-MB231 cells after 24hr, in untreated or LEV-pre-treated 2D HUVEC monolayers (30hr) (n=3). (**K**) Vascular cell adhesion molecule (VCAM-yellow) staining of untreated HUVECs (top), TNF-α treated HUVECs (middle) and LEV-pre-treated HUVECs (bottom) in 2D. Nuclei was stained with Hoechst-33342 (blue). Scale bar 50 μm. (**L**) Fluorescence of attached monocytes on 2D HUVEC monolayers (n=3).

To explore whether these changes in endothelial barrier integrity promoted leakiness, we examined initially the colocalization of dextran 1 μm microparticles with microfluidic 3D endothelial networks after perfusion of LEVs (**Fig. 5F**). We quantified a significant decrease of microparticles colocalization with endothelial networks after exposure to LEVs (**Fig. 5I**), suggesting an increase in leakiness. Moreover, we examined the transmigration of MDA-MB 231 tumor cells and the permeability of fluorescently-tagged dextran through endothelial cell-coated Transwell™ inserts. We noticed a 51% increase in the transmigration of tumor cells (**Fig. 5J**) and a 30% increase in fluorescent dextran permeability through LEV-treated HUVECs in comparison to controls (**Supplementary Fig. 23F,G**).

We then explored if LEVs affected other markers indicative of endothelial activation ^22^. Compared to untreated controls, LEVs co-cultured with HUVECs induced significant increases in vascular cell adhesion molecule (VCAM) expression (**Fig. 5K**) and nitric oxide (NO) production (**Supplementary Fig. 23H,I**). Since endothelium activation has been correlated to immune modulation via promoting monocyte differentiation ^23^ & attachment to endothelium ^24,25^, this prompted us to explore how LEVs may mediate the cross-talk between monocytes and endothelial cells. We therefore cultured untreated or LEV-pre-treated HUVECs for 24 hr, washed LEVs away and replenished with fresh media and collected CM from the cells 24 hr after. CM from LEV-pre-treated HUVECs induced differentiation of monocytes to CD206-positive, TNF-α-negative M2 macrophages, but this was not the case for untreated monocytes or monocytes exposed to CM from untreated HUVECs (**Supplementary Fig. 24**). To further examine if LEVs influenced the physical interactions between monocytes and HUVECs, we co-cultured: a) LEV-pre-treated monocytes with HUVECs, b) LEV-pre-treated monocytes with LEV-pre-treated HUVECs, and c) untreated monocytes with LEV-pre-treated HUVECs. We found significant increases in monocyte attachment to HUVECs monolayers when monocytes and/or HUVECs were pre-treated with LEVs, as opposed to untreated controls (**Fig. 5L**). Altogether, these findings suggest that LEVs shed from entrapped tumor cells drive key aspects involved in PMN formation by disrupting the endothelial barrier integrity, activating endothelial cells and interacting with immune cells.

## Discussion

Our analysis indicates that CTCs entrapment in capillary bifurcations can lead to shedding of a previously uncharacterized class of large extracellular vesicle with potential to drive key aspects of PMN formation. We demonstrated that the generation of these particles under physiological microvascular pressures is a shear stress-driven process that occurs more frequently when: a) cells encounter smaller diameter vessels and bifurcations, b) cells are larger, and c) cells are treated with F-actin polymerization inhibitors. When the sensitivity of cell shedding of LEVs to capillary-sized bifurcations, integrity of cytoskeletal elements, and fluid shear stress are taken within the context of the short timescales of particle shedding, we believe that this phenomenon is primarily a biomechanical response that likely effects many solid tumor types. We observed this behavior in all cells that we tested, including patients CTCs and cells derived from small cell lung cancer, triple negative breast cancer, melanoma, prostate cancer, pancreatic ductal adenocarcinoma, or high grade serous ovarian cancer.

The large size of masses shed from tumor cells (1.17-11.32 µm) (**Fig. 1F**) led to reductions of 3-61% in cell volume and facilitated cell transit through capillary bifurcations. The observation that particle shedding may facilitate the transit of CTCs through capillary beds is also supported by previous analyses of CTC hemodynamics that show a ∼5-6^th^ power law relationship between cell size and hydrodynamic resistance ^26,27^. Therefore, even small reductions in cell size may dramatically reduce hydrodynamic resistance and the probability of CTC occlusion.

Reductions in CTC size after transit through capillary bifurcations due to shedding of LEVs may also help to explain previous observations from patient liquid biopsies. The sizes of CTCs isolated from pre-capillary vessels of metastatic breast cancer (n=30) ^28^ and hepatocellular carcinoma patients (n=73) ^29^ were on average 25-50% larger than those isolated post-capillary from peripheral circulation. A likely contributor to this observation is that larger CTCs simply occlude in capillary constrictions and are therefore less abundant downstream of capillary beds ^28^. However, our data reveals that tumor cells can lose up to 61% (mean: 21 ± 9%) (**Supplementary Fig. 7**) of their volume upon shedding, and that shedding can occur in 82% of cells under some conditions. This suggests that forces applied to CTCs in the microvasculature may actively reduce the size of some CTCs via cell shedding, explaining partially the above clinical findings.

We show that fragments shed from CTCs during transit in capillary-sized bifurcations, match all the criteria of extracellular vesicle classification as outlined by MISEV2018 guidelines, which are: a) imaging of particles using two complementary techniques and b) the demonstration that particles consist of at least three proteins, including at least one transmembrane and one cytosolic protein ^9^. Given differences in both their mode of biogenesis and their molecular characteristics from previously identified EVs (i.e. exosomes, microvesicles or large oncosomes (LOs)) ^9,30^, we believe that these cytoplasmic fragments shed from CTCs in circulation are a novel class of a large extracellular vesicle. We propose the nomenclature of “shearosomes” for this subclass of large extracellular vesicles because of their unique requirement of shear stress for biogenesis.

Because the diameters of shearosomes (1-11 um) overlap significant with those of LOs (1-10μm) ^30^, it is difficult to distinguish these two classes of EVs from patient samples until unique surface markers or biophysical properties can be discovered and validated. Previous proteomic analysis of DU145 prostate cancer cell-derived LOs identified a total of distinct 407 proteins ^31^, far fewer than the 3382 distinct proteins we identified in shearosomes from breast cancer cells. We postulate that the greater biomolecule diversity in shearosomes is a direct result of the biomechanical nature of their biogenesis, which differs from previously identified EVs such as exosomes and microvesicles, into which, cells actively package biomolecules ^30,32^.The larger number of proteins present in shearosomes suggests that it is possible to identify numerous shearosome-specific proteins to separate them from LOs by cross-referencing the proteomes. The identification of shearosome-specific markers may allow us to explore both their clinical relevance and compare their influence against other EVs on PMN formation.

A defining characteristic of extracellular vesicles is their uptake by cells ^15,16^. Our data suggest that endothelial cells utilized macropinocytosis for shearosome uptake, whereas monocytes and macrophages relied both on phagocytosis and macropinocytosis, mechanisms typically associated with the ingestion of particles larger than 1 μm ^17^. While we observed that the size of generated shearosomes ranged from 1-11 μm, the mean size of ingested shearosomes by endothelial cells was 2.8 ± 1.2 μm (range: 0.93 – 6.8 μm), suggesting a propensity for uptake of smaller shearosomes. This could be due to inherent size limitations of macropinosome formation, restricting uptake to typically smaller LEVs. Furthermore, shearosome uptake by endothelial cells or M1 macrophages was not completely abrogated using Cytochalasin D or combo of Cytochalasin D/EIPA inhibitors, respectively. One explanation is that macropinocytosis can occur via different mechanisms i.e. lamellipodia-like protrusions, circular ruffles and bleb formation ^17^. It is plausible that endothelial cells utilize more than one of these macropinocytosis mechanisms which the EIPA inhibitor is ineffective against. Moreover, there are various inhibitors used for uptake studies that inhibit at different and sometimes overlapping levels the processes of phagocytosis and macropinocytosis ^33^. To address this, screening compounds against the two processes will be necessary to elucidate the complete repertoire of cellular uptake.

The diversity of proteins we identified in shearosomes may have contributed to their profound extracellular communication capabilities and overall reactivity to stromal cells upon uptake. Shearosomes caused: a) the polarization of monocytes and macrophages towards M2 “tumor-promoting” phenotypes, b) disruption of the endothelial barrier integrity in both 2D and 3D microfluidic models c) activation of endothelial cells, and d) co-interactions between these immune and endothelial cells. All these effects are often reported as components in the formation of pre-metastatic niches ^5,6,19,34^, which are crucial for successful metastases by regulating CTCs extravasation and eventual colonization ^6,7,35–37^. Shearosomes likely interact with many more components of the complex stromal microenvironment than we explored here. While we found that shearosomess promoted immune cell polarization towards M2 macrophage lineages, Headley et al found that dendritic cells activated by shearosomes provoked an immune response that appeared to inhibit metastatic burden ^7^. This raises the question about the overall contribution of shearosomes to malignancy, especially in the context of different immune/stromal landscapes amongst patients and cancer subtypes.

To further explore the contribution of secreted factors from shearosomes, we compared the effects of conditioned media (CM) isolated from shearosomes or direct co-cultures of shearosomes with endothelial cells or monocytes. CM from shearosomes affected VE-cad expression on endothelial cells and CD206 protein expression on monocytes, albeit at lower levels compared to the effect imposed by co-cultures with shearosomes. These findings suggest that chemical factors secreted by shearosomess are not solely driving these phenotypic changes and that shearosomes can exert effects on cells also via uptake/binding. These findings are further supported by our proteomic analysis suggesting that shearosomes contain proteins involved in protein secretion and our experiments using shearosomes generated from TGFβ silenced parental cells which had reduced ability to induce both immune cell polarization and endothelial barrier damage.

The extent to which shearosomes contribute to PMN formation remains an open question. Experiments in animal models suggest that a single gram of tumor mass can release about 4 million cells per day ^38^. Over the course of multi-year disease, primary tumors may therefore release over half a billion cells within four months. Coupled with the frequency of shearosome generation within capillary-sized bifurcations, it is plausible that endothelial and immune cells interact with appreciable numbers of shed shearosomes. Furthermore, since shearosomes are generated at the site of CTC entrapment, shearosomes may be especially capable of priming the premetastatic niche near microvasculature already enriched for CTCs. While we have evidence that shearosomes appear to influence key aspects of cells relevant to the PMN, further experiments are needed as validation. Experiments in this area however need to be carefully designed primarily because the biomechanical nature of shearosome biogenesis makes it difficult to devise negative controls where CTCs do not exhibit shedding of shearosome. For example, while our data suggests that cell treatment with Colchicine inhibits shearosome biogenesis, F-actin is a key component of the cell cytoskeleton that has crucial roles in cell and tissue homeostasis^39^ and therefore more suitable negative controls need to be identified Moreover, it is unclear how the shedding of shearosomes from CTCs and the production of EVs from primary tumors can compete or synergize towards PMN and eventual metastases formation. More sophisticated in vitro models and carefully designed in vivo studies are therefore needed to investigate the contributions of CTC-shed shearosomes on components of the pre-metastatic niche and the progression of metastasis. Altogether, the shedding of shearosomes by CTCs in the bloodstream appears to be another key component of PMN formation in cancer, and additional studies on shearosomes contribution to metastasis interactions will help us elucidate the extent of their influence in various cancer subtypes.

## Materials & Methods

### CTC isolation and enumeration

All reagents were purchased from Sigma Aldrich (UK), unless stated otherwise. Circulating tumor cells (CTCs) were isolated as previously described ^40^. In brief, 10 ml of EDTA-treated peripheral blood was collected per patient, from two small cell lung carcinoma (SCLC) patients in December 2023 following informed consent and according to ethically approved protocols. Sample collection was undertaken via the CHEMORES protocol (molecular mechanisms underlying chemotherapy resistance, therapeutic escape, efficacy and toxicity—improving knowledge of treatment resistance in patients with lung cancer), NHS Northwest 9 Research Ethical Committee reference 07/H1014/96) and The National Research Ethics Service, NHS Central Manchester research ethics committee, reference 07/H1008/229.CTCs were enriched via RosetteSep^TM^ (#15167, Stem Cell Technologies Vancouver, Canada).

CTCs were enumerated as previously described ^41^. In brief, 7.5 ml blood collected in a Cellsave preservative tube was mixed with 6.5 mL dilution buffer and then centrifuged at 800 x g for 10 mins. The sample was loaded onto the AutoPrep where iron beads coated with anti-Epithelial Cell Adhesion Molecule (EpCAM) antibody (“ferrofluid”) were used to capture candidate circulating rare cells (CRCs) from the sample. Cells that bind the magnetic capture reagent were enriched using a strong magnet which causes them to be drawn to the sides of the tube, the remaining unbound sample was aspirated and disposed. The enriched cells were stained in situ for cytokeratin (CK), CD45 (for the exclusion of lymphocytes) and DNA (DAPI). Enriched cells were deposited in a cartridge which was scanned on the Analyzer, images were presented in a gallery, scored by trained analysts and the CTC count was reported. Cells were considered to be CTCs if they were DAPI+, CK+ and CD45-.

Alternatively, during centrifugation to pellet isolated CTCs from SCLC patients, the CTCs pellet was resuspended in 250 μl of media and was introduced directly into microfluidic devices for live imaging of CTCs transit. Additionally, the previously collected supernatant was analyzed to examine the presence of cell fragments. The suspension was initially centrifuged at 150 x g for 7 min (brake applied) to remove any contaminating red blood cells. Then, the supernatant was further centrifuged at 10000 x g for 30 min to pellet potential cell-derived fragments. The pellet was resuspended in a staining solution containing CMFDA green cell tracker and Hoechst 33342, at final concentrations of 10 μM and 16.2 μM, respectively, and then imaged in wells of 96-well plate using a fluorescent inverted Leica microscope (Leica, UK). Fluorescent fragments that were CMFDA-positive and Hoechst-negative, were manually enumerated.

### CTC derived explant generation, disaggregation and culture

CDX models were generated, disaggregated and cultured as previously described according to UK Home Office Regulations and the UK Coordinating Committee on Cancer Research guidelines using approved protocols (70/8252). In brief, 10 mL of EDTA-treated peripheral blood was collected from patients with SCLC enrolled in the ChemoRES study (07/H1014/96). CTCs were enriched by means of RosetteSep and subcutaneously implanted into the flank of 8 to 16-week-old non-obese, diabetic, severe combined immunodeficient, interleukin-2 receptor γ–deficient (NSG) mice (Charles River Laboratories International, Inc., Wilmington, MA). CDX models were generated from the patients’ CTCs enriched from blood samples at pre-chemotherapy baseline or at post-treatment disease progression time points ^40,42^.

CDX tumors were grown to approximately 800 mm^3^ and the mice were killed by schedule 1 method. The tumors were removed and dissociated into single cells using the Miltenyi Biotec tumor dissociation kit (#130-095-929 [Miltenyi Biotec, Germany]) following the manufacturer’s instructions on a gentleMACS octo dissociator (#130-096-427 [Miltenyi Biotec]), as previously described. Single cells were incubated with anti-mouse anti-MHC1 antibody (eBioscience clone, 34-1-2s [ThermoFisher Scientific, Waltham, MA), anti-mouse anti–immunoglobulin G (IgG) 2a+b microbeads and dead cell removal microbead set (Miltenyi Biotec #130-090-101) and applied to an LS column in a MidiMACS Separator (Miltenyi Biotec) for immunomagnetic depletion of mouse cells and dead cells. CDX *ex vivo* cultures were maintained in Roswell Park Memorial Institute (RPMI) 1640 medium supplemented with the following components: 10 nM hydrocortisone, 0.005 mg/mL Insulin, 0.01 mg/mL transferrin, 10 nM β-estradiol, and 30 nM sodium selenite; 5 μM Rho kinase inhibitor added fresh (Selleckchem, Y27632 [Houston, TX]), and 2.5% fetal bovine serum added after 1 week at 37°C and 5% carbon dioxide ^40,42^.

### Cell culture

Cell lines were obtained from ATCC, unless noted otherwise. MDA-MB-231, OVSAHO, LNCaP, Me290, PDAC and B16F10 cell lines were maintained in StableCell™-Dulbecco’s Modified Eagles Medium (DMEM) media, supplemented with 10% (v/v) FBS and 1% (v/v) penicillin/streptomycin and incubated at 37°C/5% CO_2_. Human umbilical vein endothelial cells (HUVECs) were incubated at 37°C/5%CO_2_ and supplemented in 10% FBS and EGM™-2 Endothelial Cell Growth Medium-2 BulletKit™ (Lonza, Switzerland). HUVECs and all other cancer cell lines were sub-cultured as adherent cells every 2-3 days.

THP-1 monocytes were maintained in RPMI-1640, further supplemented with 10% (v/v) heat-inactivated fetal bovine serum (FBS) and 25 mM 4-(2-hydroxyethyl)-1-piperazineethanesulfonic acid (HEPEs) buffer (Gibco, US), 2.5g/l Glucose (Merck, UK), 1mM sodium pyruvate (Gibco, US), 1% (v/v) Penicillin/Streptomycin and 50pM β-mercaptoethanol (Gibco, US) final concentrations, in suspension. After thawing, THP-1 monocytes were passaged every 3 days at a 1:4 split ratio. M1 macrophages were obtained as previously described ^20^ by first treating monocytes with 150 nM phorbol-12-myristate-13-acetate (PMA) for 24hr to obtain M0 macrophages and then further treatment with Interferon gamma (20ng/ml IFN-γ/Biotechne, UK) for 24 hr to obtain M1 macrophages ^20^.

Primary cancer associated fibroblasts (CAFs) cells were provided by Breast Cancer Now Tissue Bank and were maintained in DMEM:F12, (supplemented with 1% (v/v) penicillin/streptomycin, amphotericin-B (2.5µg/ml) and 10% (v/v) heat-inactivated FBS and incubated at 37°C/5% CO_2_. Primary CAFs were sub-cultured every 3 days.

### Preparation of single cell and cluster suspensions

For single cell experiments, cancer cell lines and primary CAFs (mentioned above) were harvested from 80-90% confluent 48-well plates (corning, UK) or 25cm^2^ tissue culture flasks (Corning, UK), by washing with PBS, followed by incubation with 0.25% trypsin (5 min at 37°C/5% CO_2_), inactivation with 10% serum-containing media, centrifugation at 180 x g for 5 min and resuspension of the pellet with fresh media in new culture flasks. Alternatively, THP-1 monocytes (suspension cell line) were collected, centrifuged as above and enumerated. THP-1 derived M0 macrophages were harvested as adherent cells and after trypsinization were scraped using cell scrapers. Where required, the cell pellet was resuspended with staining solution for 15 min at 37°C/5% CO_2_. Staining solutions were prepared to a final volume of 1 ml by diluting to final concentrations of Calcein-AM (5 μM) and/or Hoechst 33342 (16.2 μM) in PBS as required.

For cell cluster experiments, MDA-MB 231 cells were initially harvested, as described above, except the pellet was resuspended in FBS-containing media and cells were added to round bottom 96-well plates (Corning, UK) for 1 day at 37°C/5% CO_2_. Round 96-well plates were pre-treated with 1% Pluronic F-127 (Thermo Fischer Scientific, UK) at room temperature for 1 hr to facilitate formation of spheroids. To stain, 200 μl of the solution with both Calcein-AM and Hoechst (described above) was added directly in the wells. Stained spheroids were gently resuspended to form clusters and suspensions were collected.

### Chemical treatment of cells

Single cell and/or cluster suspensions of MDA-MB 231 cells were pre-treated with Cytochalasin D (Cyto-D) (Merck Life Science, UK), or Colchicine (Colch) (Merck Life Science, UK), for 3 hr at 37°C/5% CO_2_, at final working concentrations of 10 μM, 50 μM respectively, as required.

For internalization inhibition studies, HUVECs, THP-1 monocytes or THP-1 derived M1 macrophages were pre-treated with Cyto-D or 5-(N-Ethyl-N-isopropyl)-Amiloride (EIPA) for 3hr at 37°C/5% CO_2_, at final working concentrations of 10 μM and 50 μM, respectively.

To generate positive controls of disrupted HUVEC endothelium, HUVECs were pre-treated with tumor necrosis factor alpha (ΤNF-α) at a final working concentration of 0.1 μg/ml for 30 hr.

To evaluate intracellular production of nitric oxide by HUVECs, untreated or LEV co-cultured HUVECs (elaborated later) were stained with DAF-FM dye (ThermoFischer Scientific) for 1 hr at a final concentration of 5μM at 37°C/5% CO_2_.

### Microfluidic capillary devices design, fabrication & preparation

Capillary bifurcations were designed according to Murray’s law (**Equation. 1**):

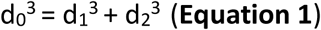

where, d_0_, d_1_ and d_2_ are the effective diameters of the parent and daughter channel vessels in a capillary bifurcation.

To calculate the effective diameter of microchannels narrower than 10 μm, we used the Eqn. 2.

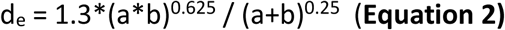

where, a is the width of the channel and b the height.

We then used equations 1 and 2 to generate 13 distinct capillary bifurcations and two control geometries, as outlined in Supplementary Table 1. Distinct geometries of capillary bifurcations were generated by altering channel dimensions, the symmetry or angle between the two daughter channels, and whether or not the daughter channels had equal or unequal dimensions. These were designed to recapitulate the diversity of bifurcations present in in vivo capillaries ^1^.

All devices were fabricated using photolithography and soft lithography. Briefly, silicon wafers (MSE Supplies, Germany) were sterilized with 99% IPA and 100% acetone and oxygen plasma treated (Harrick plasma cleaner) at 0.5 Tor O_2_ at 30W for 1 min before spin coating with a GM1050 SU-8 negative photoresist (Gersteltec, Swtzerland). Wafers were spun first at 1050 rpm for 10 s and then at 950 rpm for 45 s and then heated at 65 and 95°C for 2 and 10 min, respectively. SU-8 coated wafers were exposed to UV photolithography with a UV-KUB 3 mask aligner (KLOE, France) to pattern the capillary designs for all masters, using chromium glass masks (Microlitho, UK). After repeating heating steps from earlier, wafers were developed in SU-8 developer (Kayaku Advanced Materials, USA) for 5 min. The height of the devices was set to 7 μm for constricted capillaries and 20 μm for non-constricted geometries and were verified, using an optical surface profilometer (FILMETRICS, USA), within ±5% range of intended height. The master wafer was taped in a petri dish and Sylgard^TM^ 184 Polydimethylsiloxane prepolymer with its crosslinker (Dow Corning) were mixed at 10:1 w/w ratio, poured onto silicon master molds, degassed for 30 min and transferred in an oven, left to polymerize overnight at 65°C.

Polymerized PDMS replicas were cut out, punched in their inlets and outlets with 4- and 2-mm biopsy punch (IHC WORLD, USA), respectively, and then plasma treated with oxygen (0.5 mm Tor, 30W, 1 min/Blackholelab, France), bonded at glass slides and heated at hotplate for 10min at 95°C. The formed microfluidic devices were flushed 1x with 70% v/v ethanol to sterilize and wet the devices, washed 1x with PBS and 3% Bovine Serum Albumin to prevent cell attachment within the devices. To perform cell experiments, 80-cm long sections of 2 mm outer diameter tubing (Qosina, USA) were connected to 10-ml polypropylene syringes, coupled with 22-gauge needles (Needlez, UK). The tubing/syringe set-up was prefilled with 2 ml PBS, the plungers were removed and the free ends of tubing were connected to the outlets of the devices.

### Computational Fluids Dynamics

Microfluidic channel geometries were imported from AutoCAD into ANSYS 2025 R1 Discovery. Meshing was conducted using ANSYS Meshing using quad-dominant sheet body structured meshes of element sizes of 1 µm. No slip boundary conditions were used at walls and gauge pressure of the inlets were set at 1961 Pa for all except for non-constricted bifurcations which were set at 490 Pa. ANSYS Fluent pressure based solver was used with water (density = 998 kg/m³, viscosity = 0.001 Pa·s). Coupled solver scheme, second order pressure and second order upwind momentum spatial discretization. Convergence criteria set to 1×10⁻^5^ for continuity and momentum equations, and 200 iterations. Fluid shear stress was calculated as the product of strain rate and molecular viscosity.

### Capillary transit experiments

30-to 50 μl of single cell suspensions of patient CTCs or 0.2×10^6^ cells/ml of CDX cells, cancer cell lines, primary CAFs, THP-1 monocytes and THP-1 derived M0 macrophages (as mentioned above) were added in the inlets of microfluidic devices, whose outlets were connected to the tubing/syringe set-up. The syringe was lowered 10-30 cm below the level of the device for constricted bifurcations and 5-10 cm below for non-constricted bifurcations to generate physiological pressures ^43^ and the cells were collected from the outlets of the devices and by emptying the liquid content from the tubing into a 15 ml falcon tube. Similarly, cell transit was tested both in 10 cm and 20 cm hydrostatic pressure to investigate the impact of various shear stress values on cancer cell shedding. Where required, cell transit through the microfluidic devices (n=3 per geometry) was imaged using Nikon Eclipse Ti2 multi-fluorescent inverted microscope (Nikon, UK) with Okolab incubated stage (Nikon, UK). For no-flow control experiments, cells were added in the inlets of microfluidic devices (n=3) but without the implementation of flow, for the same amount of time that the microfluidic transit experiments lasted before extraction from device inlets. As additional no-flow control experiments, MDA-MB 231 cells were stained as previously and transferred in Eppendorf tubes, subjected to centrifugation forces (8000xg) for 3 minutes, transferred to triplicate wells and the LEV: cell ratio was calculated. Cells without centrifugation were also analyzed as control. When required, cells were pre-treated with Cyto-D or Colch, as described above.

### Image & Video analysis

The analysis of individual cells and clusters transiting through capillary bifurcations and control devices was conducted in NIS-Elements (Nikon, UK). As described above/below, cell shedding was verified by detachment of cytoplasmic-stained fragment, but negative for nuclei (Hoechst). Shedding frequency for single cell experiments was calculated by dividing the number of cells that shed at least one LEV by to the overall number of cells that transited through a capillary bifurcation. For clusters, shedding frequency was calculated by dividing the number of clusters that shed at least once to the overall number of clusters that transited.

The spherical diameters of cytoplasmic or nuclear volumes were calculated as previously described ^26^. Briefly, from video frames of the cells within microchannels, cells were constricted only in the vertical axis. Alternatively, the volume of cells trapped in the capillary bifurcation were calculated by a method similar to what was previously described^26^.

Cell lysis during transit in capillary devices was quantified by examining cells that experienced loss of Calcein-AM localization. The lysis frequency of shedding cells was extrapolated by dividing the number of cells that were lysed post-shedding to the number of all cells that shed.

### Viability assay

100 μl of cell suspensions (10^5^ cells/ml) were added in each well of a 96-well plate and left to attach and grow for a day. Next, cells were treated with Cyto-D or Colch, as described above, for either 3 or 24 hr at 37°C/5% CO_2_. Viability of cells at 0 hr was also examined as a control. The media was removed at the relevant timeframe and the staining dye for the live/dead assay was added for 15 mins at 37°C/5% CO_2_. A staining solution consisting of 1X Propidium Iodide (Thermofischer Scientific, UK), 5 μM Calcein-AM (5 μM) and 16.2 μM Hoechst 33342 in 1 ml media was added to cells. For each condition, 4-5 multifluorescent images were obtained at 20x magnification using the Nikon microscope. Alive cells were defined as Calcein^+^, Hoechst^+^ and Propidium Iodide^-^, whereas dead cells as Calcein^-^, Hoechst^+^ and Propidium Iodide^+^. Viability was assessed by extrapolating the percentage of alive cells compared to overall number of cells (Hoechst-positive). Images were analyzed using the Nikon software.

### Post transit cell analysis

50 μl of suspensions of cells and their shed LEVs were collected into wells of 96 well plates as described above. The diameters of LEVs (Hoechst 33542-negative, Calcein-AM positive) were measured using both NIS-Elements (manually measuring diameters using the Annotations and Measurements command) and automated Cell Profiler scripts. In addition, fluorescent images of LEVs (positive for greencell tracker) and cells (positive for green cell tracker and Hoechst) were obtained and analyzed via CellProfiler scripts to measure the diameter of LEVs. In particular, the workflow consisted of i. uploading of raw fluorescent images, ii. identification of objects (both LEVs and cells), iii. separation of cells from LEVs that are negative for Hoechst (blue signal) and iv. measurement of object diameter. Cell death post-transit was assessed by performing a viability assay as elaborated above 3 hours after transit of either MDA-MB 231 or THP-1 monocytes and calculating the percentage of dead cells per condition (n=3). LEVs and cells per condition were enumerated manually (n=3), by analyzing multiple representative multifluorescent images across each well, establishing thus a LEV: cell number ratio.

### Electron microscopy

Cells and their shed LEVs were collected as described above, and the solution was centrifuged at 180 x g for 5 min to separate cells (pellet) and retain LEVs in the supernatant (described below in more detail).

For cryogenic Transmission Electron microscopy, 3µl of concentrated LEV suspension was applied by pipette to glow-discharged Quantifoil R2/2 carbon support films on 200mesh EM grids. The grids were blotted and vitrified by plunge-freezing in ethane using an FEI Vitrobot MkIV with chamber at 22°C and 100% humidity. Grids were imaged at 200keV on an FEI Glacios TEM under cryogenic conditions. Overview images for LEV characterization and diameter measurement were collected at 50µm underfocus at a range of low magnifications. High magnification imaging to characterize the LEV boundaries was performed using a Falcon4i direct electron detector (Thermo Fisher Scientific) and zero-loss-centred energy filtering at 10eV slit width using a Selectris energy filter (Thermo Fisher Scientific). Exposures were collected of 40e^-^/Å^2^ total dose at 1.18Å pixel size and 3µm underfocus, generating movies that were aligned and summed in Thermo Fisher’s Velox software. Measurements on images were made manually using IMOD.

For Scanning Electron microscopy, LEVs were transferred onto 10 x 10 mm square glass slides that were positioned into wells of a 24 well plate. After 1 day, the media was removed and the glass slides were fixed with 4% formaldehyde (diluted in PBS) for 15 min at room temperature to crosslink proteins. Then, glass slides were chemically treated with 1% (v/v) Osmium tetroxide (Agar Scientific, UK) to crosslink lipids. The fixed samples were dehydrated with a sequence of washes with increasing concentrations of ethanol, starting from 30% and continuing with 40%, 50%, 60%, 70%, 80%, 90% and 100% (v/v) ethanol. After dehydration, the samples were chemically dried, by using Hexamethyldisilazane (Merck, UK) for 3 mins. A sticky carbon tape was attached to each square glass slide and silver-containing glue was brushed onto the samples to enhance conductivity. Finally, a 15 μm thick layer of Nickel was deposited onto the samples using a sample sputter coater (Q150TS, Quorum technologies, UK), to improve image resolution. Prepared samples were loaded into the stage holder of the scanning electron microscope (Zeiss Auriga 40 Cross beam scanning electron microscope, Zeiss, Germany). 5 kV of acceleration voltage (using secondary electron detection) was applied to the sample. Samples were scanned to identify LEVs and images at 1000-8000x magnification.

### Large EV generation, isolation and purification

2 ml of untreated MDA-MB 231 cell suspensions (10^6^ cells/ml) were pre-stained with a CMTPX red or CMFDA green cell tracker (ThermoFischer Scientific, UK) for 1 hr at a final concentration of 5 μM. Then, 100 μl of the suspension was added in the inlet of capillary bifurcation device (ENA_7/7) (Supplementary Table 1) and operated as described above. Before the inlet emptied completely, additional 100 μl of the suspension was added sequentially until the entire 2 ml volume passed through the device. Both cells and shed LEVs were collected as described above. Two distinct methodologies were used to purify LEVs from cells in effluent solutions. In the first method, filtration, cells and LEVs containing suspensions were transferred to 10 ml syringes and passed through Whatman Polydisc filters with a pore diameter of 10 μm which permitted LEVs to pass through. In the second method, centrifugation, the cell/LEVs suspensions were centrifuged for 5 min at 180 x g, separating cells into the pellet from LEVs in the supernatant. Supernatants were collected and further centrifuged for 30 min at 10000 x g to form LEV pellets which were later resuspended in fresh media.

To evaluate LEV purity after filtration, 100 μl of samples before and after filtration were transferred into separate wells of a 96 well plate and multifluorescent images of each well were taken as described above. After centrifugation, LEV purity was evaluated by flow cytometry (described below) of 500 μl samples taken before centrifugation or in the pellet and supernatant after centrifugation in an Amnis CellStream flow cytometer (Luminex, US).

### Post transit cell proliferation

To evaluate the proliferative capabilities of MDA-MB231 cells after transit, 5 × 10^5^ cells were flowed through a microfluidic device, the effluent collected, and cells separated from LEVs as described above. Cells were enumerated using a hemocytometer (as above). 250 μl of cell suspension of 0.5×10^6^ cells/ml density were added to wells of a 48-well plate. Equal numbers of cells that did not transit through devices were added to separate wells as controls. Images were taken 4 hr after seeding and then every day up to 7 days post-seeding. In other experiments, post-transit cells and control cells were left to grow for 3 days in 48 well plates and were then harvested and enumerated using a hemocytometer.

### Co-culture of cells with LEVs or conditioned media

HUVECs, THP1 monocytes or THP1-derived M1 macrophages were independently stained with a CMTPX red cell tracker and were independently co-cultured with MDA-MB-231 derived LEVs pre-stained with CMFDA green cell tracker (as prepared elsewhere) at cell: LEV ratios of 3:1. For LEV internalization studies, co-culture was allowed to proceed for 16 hours. For all other studies, the co-culture was performed for 30 hrs to allow for phenotypic changes and gene expression changes.

Conditioned media (CM) was collected from the cell culture supernatant of MDA-MB 231 cells or HUVECs cells cultured for 1 day (as described above). Cell culture supernatant was centrifuged at 180 x g for 5 min and the supernatant collected as CM. To collect CM from co-cultured HUVEC cultures, HUVECs were first co-cultured with LEVs for 30hr. Media was removed from each well to remove non-internalized LEVs and fresh HUVEC media (elsewhere) was added. After 1 day of culture at 37°C/5% CO_2_, media was collected and processed as above to collect CM from LEV-pre-treated HUVECs. CM from LEVs was collected by purifying and centrifuging LEVs as above, resuspend them in fresh media, culture them for at least 24hr at 37°C/5% CO_2_ and centrifuge again at 10000x g for 30 min to collect supernatant, designated as LEVs-CM.

### Confocal microscopy

HUVECs, monocytes, or M1 macrophages were co-cultured as described above with LEVs. Cells were pre-stained with cell tracker red (CMTPX) (adjusted to magenta) and LEVs with cell tracker green (CMFDA). After 16 hr of co-culture, 96-well plates were loaded in the stage of a Leica SP8 inverted scanning confocal microscope (Leica, UK). Images were obtained using a Z step of 2 μm. Excitation and emission wavelengths for cells and LEVs were 561/611 and 488/528nm, respectively. Internalization of LEVs by cells was validated by applying orthogonal cross sectioning to image a cell with an LEV in all pairs of coordinates i.e. x-y, x-z and y-z. Moreover, the diameter of LEVs that were internalized by HUVECs was measured, after firstly verifying LEVs internalization.

### Immunocytochemistry

A standard protocol was adopted for immunocytochemistry (ICC) staining. Cells or MDA-MB 231 derived LEVs were fixed with 4% (v/v) formaldehyde at room temperature for 15 min. Then, cells or LEVs were washed 3x with PBS for 3 min (per wash), incubated with 0.1% (v/v) Triton X at room temperature for 5 min, washed 3x with PBS for 3 min (per wash) and finally incubated with the primary antibody overnight at 4°C. After overnight staining, cells were washed 3x with PBS for 3 min (per wash) and incubated for 2 hr at room temperature with a secondary antibody. Cells were washed and stained for Hoechst-33342 as required at room temperature for 5 min and then imaged in the Nikon or confocal microscope (described below). Multifluorescent images across each condition were obtained at 20x magnification. Primary and secondary antibodies against proteins per cell type were used, as detailed in Supplementary Table 6. In monocyte experiments, where required, a combination of antibodies was used. Samples were further analyzed via flow cytometry as detailed elsewhere (Supplementary Materials & Methods). Excitation and emission wavelengths used for obtaining relevant dot plots are depicted in Supplementary Table 7.

All antibodies were purchased from Abcam (UK), except for primary rabbit anti human monoclonal antibody CXCL10 (ThermoFischer Scientific, UK).

### Flow cytometry

MDA-MB 231 cells, HUVECs, monocytes or purified LEVs (the latter derived from MDA-MB 231 cells as described above) were stained as described above (see also Supplementary Table 6). 500 μl of each sample (10×10^5^ cells/ml or 20×10^4^ LEVs/ml) were loaded into an Amnis Cell Stream flow cytometer (Luminex, US) and samples were run at a flow rate of 20 μl/min. At least 1000 events per sample and condition were recorded. Excitation and emission wavelengths used for obtaining relevant dot plots are depicted in Supplementary table 7. For all histograms generated, antibody fluorescence intensity was represented in the x axis and frequency in the y axis. As a general approach, unstained samples and single-color controls were used for setting gating thresholds.

For LEVs internalization studies, pre-stained cells with a red cell tracker (HUVECs or monocytes or M1 macrophages) were co-cultured independently with pre-stained LEVs with green cell tracker for 16 hr. Internalization frequency was defined as the number of double positive events, cells that had internalized LEVs compared to overall number of cells. Where required, cells were pre-treated with Cyto-D, EIPA or combination, as described elsewhere and then LEVs were added. In each case, unstained cells, untreated cells (stained) and LEVs alone (stained), were used as controls for gating the limits for double positive dot plots (cells that had internalized LEVs) and samples were analyzed as above. For relevant experiments, fluorescent beads of 5- and 10 μm (microParticles GmbH, Germany) were used.

### RNA extraction and real-time PCR

HUVECs, THP-1 monocytes, or THP-1 derived M1 macrophages were independently co-cultured with LEVs generated by MDA-MB 231 cells (as described elsewhere) at ratios of 3×10^5^ cells:1×10^5^ LEVs. 3×10^5^ of the above cells were cultured alone as controls. Additionally, 2×10^5^ LEVs were cultured alone to investigate their RNA content. Cells or LEVs were centrifuged at 180 x g for 5 min or at 10000 x g for 30 min to pellet cells or LEVs, respectively. Then, total RNA was isolated from each sample of cells or LEVs using RNeasy Plus Mini Kit (Qiagen, USA), based on manufacturer’s instructions. The quality, quantity and purity of the extracted RNA were determined using Nanophotometer (IMPLEN, USA). 1 μg of sample was used for reverse transcription, performed via iScript cDNA synthesis kit (Bio rad laboratories, UK). Then, 50 ng of cDNA per sample was used and amplified by applying real-time PCR, using the equipment of Applied Biosystems^TM^ StepOne^TM^ Real-time PCR System (Fischer Scientific, UK). SYBR Green PCR master mix (ThermoFischer Scientific, UK) was used for the final PCR reaction. Relative mRNA expression was extrapolated using the 2 ^− ΔΔCt^ method, for all genes and their primer sequences, as listed in Supplementary Table 8.

### Protein concentration & gel electrophoresis

Initially, suspensions of each fraction were washed 1x with chilled (4°C) PBS and lysed using 300 μl of radioimmunoprecipitation assay (RIPA) lysis buffer (Thermo Fischer Scientific, UK) for 15 min on ice in a plate shaker. All fractions were centrifuged at 18900 x g for 15 min to remove debris and the supernatant containing the lysate was collected. All other previous centrifugation steps were at 180 x g for 5 min and at 9600 x g for 30 min, for cells and LMPs, respectively. The protein lysates were stored at −80°C or placed on ice for immediate use.

Protein quantity of MDA-MB 231 control cells, cells post-transit (bifurcated capillary devices) and LEVs were measured using Bicinchoninic acid (BCA) assay kit (Life Technologies, UK), per manufacturer’s instructions. The absorbance of each sample at 562 nm was analyzed using a Varioskan Flash plate reader (ThermoFischer Scientific, UK).

For qualitative protein analysis, gel electrophoresis was performed. The separating gel was prepared by mixing 1.6 ml PBS, 2 ml Acrylamide/Bis-acrylamide 30% (v/v) solution, 1.3 ml 1.5 M Tris (Life Technologies, UK), 50 μl 10% (v/v) Sodium Dodecyl Sulfate (Thermo Fischer Scientific), 50 μl 10% (v/v) Ammonium Persulfate (Bio Rad, UK) and 2 μl tetramethylethylenediamine (Bio Rad, UK). The stacking gel was prepared by mixing 1.4 ml PBS, 0.33 ml Acrylamide/Bis-acrylamide 30% (v/v) solution, 0.25 ml 1.5 M Tris, 20 μl 10% (v/v) Sodium Dodecyl Sulfate, 20 μl 10% (v/v) Ammonium Persulfate and 2 μl tetramethylethylenediamine. Both gels were left to polymerize at room temperature for 20 min. Both the separating and stacking gel were enclosed in a glass cassette (Bio Rad, UK) and incorporated in the electrophoresis chamber (Bio Rad, UK). 250 μg/ml of protein samples (15 μl) are mixed with 5μl Laemmli protein buffer (Bio Rad, UK) and heated for 10 min at 95°C. 12 μl of each sample and protein ladder solution (Bio Rad, UK) are added across the wells of the gel. The electrophoresis chamber was filled with 10% (v/v) Sodium Dodecyl Sulfate and the samples were run for 120 min at 125 V and 400 A. The gel was then removed carefully from the enclosed glass cassette and stained with 50 ml Coomassie blue (Bio Rad) in a glass sealed container for 1 hr at room temperature on a plate shaker (Thermo Fischer Scientific, UK). The staining solution was removed and the gel was destained using 50 ml of Coomassie brilliant blue R-250 destaining solution (Bio rad, UK) for 1 hr at room temperature on a plate shaker. Images were obtained with a mobile phone (Redmi, China).

### Western blotting

LEVs purified from MDA-MB-231 cells were collected and transferred into a 15 mL tube, then centrifuged at 1500 rpm for 10 minutes. Following centrifugation, the supernatant was discarded, and the cell pellet was resuspended in 500 µL of 0.1% NaCl solution. The suspension was transferred into a 1.5 mL Eppendorf tube and further centrifuged at 13,000 rpm for 10 minutes. The resulting supernatant was again discarded. Cell lysis buffer was freshly prepared by mixing 5 mL RIPA buffer (Sigma-Aldrich, 0000351333), 100 µL PhosSTOP (Roche, 04906837001), and 10 µL protease inhibitor (Sigma-Aldrich, I3786-1ML). The cell pellet was resuspended in 20 µL of cell lysis buffer, gently agitated, and incubated on ice for 15 minutes. After incubation, the suspension was centrifuged at 13,000 rpm for 10 minutes, and the clarified supernatant (cell lysate) was carefully transferred to a clean Eppendorf tube.

Protein concentration in lysate was determined using a bicinchoninic acid (BCA) assay. The BCA reagent was prepared by combining 49 mL Reagent A (Thermo Scientific, 23228) with 1 mL Reagent B (Thermo Scientific, 1859078). Lysate samples were diluted at a ratio of 1:20 (1.5 µL cell lysate with 28.5 µL cell lysis buffer) and maintained on ice. A protein standard curve was generated using bovine serum albumin (BSA) at concentrations of 1000, 750, 500, 250, 125, 25, and 0 µg/mL, diluted in cell lysis buffer. Standards and diluted lysate samples (25 µL each) were loaded into a 96-well plate, followed by the addition of 200 µL BCA reagent to each well. The plate was shaken gently for 30 seconds, incubated at 37°C for 30 minutes, and absorbance was measured at 570 nm using a Tecan plate reader. Based on BCA results, lysate volumes containing 30 µg of total protein were calculated. Samples were prepared to a final volume of 50 µL comprising 12.5 µL of 4X LDS buffer (Invitrogen, NP0007), 5 µL of 10X reducing agent (Invitrogen, NP0009), the calculated lysate volume, and cell lysis buffer.

Electrophoresis running buffer was prepared by combining 950 mL ultrapure water with 50 mL NuPAGE SDS MOPS running buffer (20X). Samples were loaded onto a 4–12% Bis-Tris gel (NP0321BOX). The gel cassette was initially placed in a shallow bath of running buffer to remove the tape and comb, ensuring no gel residues obstructed the wells. The gel cassette was then positioned in a mini gel tank (ThermoScientific, A25977) filled with running buffer, with wells oriented forward. Samples were vortexed briefly, heated at 90°C for 3 minutes to denature proteins, then cooled to room temperature before loading onto the gel. PageRuler Ladder (5 µL; ThermoScientific, 26619) was loaded into the first lane for molecular weight referencing. Subsequently, 20 µL of the prepared sample was loaded into individual well, and electrophoresis was performed at 140 V for 1 hour.

Post-electrophoresis, the gel cassette was opened in a shallow bath of running buffer, and the gel was gently released using a scraper. The gel was transferred to an iBlot cassette, air bubbles were carefully removed using a roller. The cassette was closed and transferred to an iBlot 3 Western Blot Transfer System, run using the broad molecular weight range setting. Successful protein transfer was confirmed by visible ladder transfer onto the membrane. The membrane was briefly rinsed in tris-buffered saline (TBS) and blocked in a solution consisting of 5 mL Licor Intercept Blocking Buffer (Licor, 927-60001) and 5 mL ultrapure water for 1 hour at room temperature on a rocker.

Primary antibody incubation was performed overnight at 4°C to detect HSP90 and CD81 as previously, using a solution containing 5 mL Licor Intercept Blocking Buffer, 5 mL ultrapure water, and 10 µL Tween-20 (Sigma-Aldrich, P7949). Vinculin-(E1E9V) XP® (Cell Signal Technologies #13901) was used as dye loading control at 1:1000. After incubation, membranes were washed three times in TBST (TBS containing 0.1% Tween-20) for 10 minutes each at room temperature. Secondary antibody incubation followed the same protocol as previously, with subsequent washes in TBST performed three times, each for 5 minutes. Finally, the membranes were scanned using the Li-cor Odyssey XF imaging system, detecting the fluorescent signals at wavelengths corresponding to the secondary antibodies used.

### Sample preparation for mass spectrometry & proteomics

500,000 control cells, cells post-shedding and purified LEVs were prepared as previously. Cells and LEVs were pelleted at 180 x g for 5 min or at 10,000 x g for 30 min, respectively, to remove media and washed 1x with cold PBS for 5 min. Pelleted samples were lysed by adding 200 μl of 8 M freshly prepared urea (SIGMA-ALDRICH, St Louis, MO, USA). This lysis step was performed on ice for 20 min with brief vortexing throughout. Samples were then transferred to −80^ο^C for storage. Protein quantification was performed using Pierce™ Protein Assay Kit (Thermo Scientific, Rockford, IL, USA) per manufacturer instructions.

A total of 0.4 μg/μl protein (150 μl total) was used per sample for sample processing and digestion. Reduction of disulfide bonds was carried out by adding Dithiothreitol (DTT, SIGMA-ALDRICH, St Louis, MO, USA) at a working concentration of 10 mM and incubation for 40 min and 50°C. Samples were then alkylated using 50 mM working concentration of iodacetamide (IAA, SIGMA-ALDRICH, St Louis, MO, USA) and incubation for 30 min at room temperature in the dark. Samples were then diluted with 100 mM ammonium bicarbonate (ABC) to achieve a final urea concentration of 2 M. This was followed by trypsin digestion where trypsin (Pierce™ Trypsin Protease MS-grade, Thermo Scientific, UK) was reconstituted in 100 mM ABC and was used at a concentration of 1 μg per 100 μg of total protein. Sample digestion was carried out for 18 hours at 37°C. Trypsin digestion was then quenched with 0.5% (v/v) Trifluoroacetic acid (TFA, Fisher Scientific, Leicestershire, UK) and samples were desalted using SepPak C18 Plus cartridges (Waters, Milford, MA, USA). Finally, peptides were dried using a speedvac concentrator (Savant™ SC250EXP SpeedVac™, Thermo Scientific) and stored at −80C.

For mass spectrometry analysis, peptides were reconstituted in 0.1% trifluoracetic acid (TFA) at a concentration of 200 ng/μL for liquid chromatography–mass spectrometry (LC–MS) Data Independent Acquisition (DIA) analysis. DIA analysis was performed on an UltiMate 3000 system (Thermo) coupled with the Orbitrap Ascend Mass Spectrometer (Thermo) using a 25 cm capillary column (Waters, nanoE MZ PST BEH130 C18, 1.7 μm, 75 μm × 250 mm). 1 μg of peptide was analysed per sample. DIA was performed over a 100 min gradient, 5%-35% mobile phase B composed of 80% acetonitrile, 0.1% formic acid. MS1 spectra were collected with mass resolution of 60K in the m/z range of 380-985, with Maximum Injection Time 100 ms and automatic gain control (AGC) 4×105. DIA MS2 spectra were collected with higher energy collisional dissociation (HCD) fragmentation collision energy (CE) 32%, orbitrap resolution 15K with isolation window 10 m/z and 1 m/z overlap, Maximum Injection Time 40 ms and AGC 1×105. Raw data were processed in the DIA-NN software version 1.8.1 ^44^ for protein identification and quantification using a fasta file containing reviewed UniProt human proteins ^45^. Carbamidomethylation of C and oxidation of M were defined as fixed and variable modifications respectively. Retention time (RT)-dependent cross-run normalization was selected with match between run (MBR) enabled and proteins were filtered at 1% False discovery rate (FDR). Gene identifier names (Gene IDs) were filtered to only include proteins identified in at least 75% (2/3) of samples. Proteins identified in LEVs were searched against gene sets from the Reactome database (https://reactome.org/). The top 15 gene sets were ranked based on the number of overlapping gene IDs with LEVs. LEVs’ protein annotations relevant to protein secretion molecular functions, as analyzed by pathway analysis, were extracted from the database.

### LEV uptake under flow

For this series of experiments, a microfluidic device (length: 50 mm, width: 1 mm), previously established in the lab was used ^46^. The inlet of a microfluidic device was connected to the outlet of a 12V adjustable peristaltic dosing pump (Amazon, UK, G628-1-2) via 50 cm long tubing of 2 mm diameter (Qosina, US). The outlet of the microfluidic device was connected with a 50 cm long tubing of 2 mm diameter and the free end was inserted in a 50 ml falcon tube that was prefilled with 10 ml of media. A third piece of 50 cm long tubing of 2 mm diameter was inserted in the inlet of the peristaltic pump and its free end was also inserted in the same 50 ml falcon tube. 10×10^4^ purified and pre-stained LEVs (CMFDA green cell tracker) were added in the falcon tube either with 30×10^4^ pre-stained THP-1 monocytes (CMPTX red cell tracker) or with 30×10^4^ pre-stained THP-1 derived M1 macrophages (CMTPX red cell tracker) separately. In each experiment, the peristaltic pump was turned on for continuous circulation of cells and LEVs. After 16 hrs of co-culture under flow, media was collected, centrifuged at 180 x g and the pellet was resuspended in fresh media. 500 μl of each sample was collected and LEV internalization was compared using flow cytometry as described above.

### RNA silencing

10^5^ MDA-MB-231 tumor cells were grown in each well of a 24-well plate for a total of 2×10^6^ cells. TGFb was knocked down by following the siRNA transfection protocol using DharmaFECT^TM^ (Horizon Discovery) transfection according to manufacturer recommended protocols. Cells were cultured for additional 48 hr and media was changed every day. Then, cells were harvested and used to generate and count LEVs as described previously.

### 2D HUVEC cell and dextran permeability assays

15×10^3^ HUVECs were cocultured in wells of a 96 well plate with 5×10^3^ purified LEVs derived from MDA-MB 231 cells or 5×10^3^ purified LEVs derived from siTGFb-transfected MDA-MB 231 cells as described above. Equal numbers of HUVES but cultured without LEVs were used as negative controls. HUVECs were pre-stained with a CMFDA green cell tracker. At 0hr, both fluorescent and brightfield images were obtained at 10x and 20x magnification. After 30hr, similar images were obtained again to characterize the extent of HUVEC monolayer disruption. In other experiments, HUVECs were stained with VE-cad and Hoechst instead of CDFMA cell tracker as described above. Cell-free vs. cell-containing areas were extrapolated using ImageJ.

HUVEC monolayer disruption was further characterized by performing a dextran permeability assay. Millicell hanging cell culture inserts of either 0.4 μm or 8 μm pores (Merck, UK) were transferred in wells of a 24-well plate. The well was filled with 900 μl around the cell culture insert and 200 μl of HUVECs (75×10^3^ cells/ml) were then transferred inside the cell culture inserts and left for a day to attach and form the monolayer. Then, HUVECs were treated with 5×10^3^ purified LEVs derived from MDA-MB 231 cells (as described elsewhere) for 30 hr. Then, 100 μl of fresh media was added to 0.4 μm or 8 μm, containing either 10,000 molecular weight Dextran Alexa Fluor^TM^ 647 (ThermoFischer Scientific, UK) or 10×10^4^ cells/ml pre-stained MDA-MB 231 cells (green cell tracker), respectively, for 24 hr. 100 μl of media was collected from the bottom of the well at 24 hr and fluorescence was measured using a Varioskan Flash plate reader (ThermoFischer Scientific, UK) at 650/668 nm values of peak excitation/emission. The fluorescence values obtained at 24hr were normalized initially based on blank samples (cell-free media) and baseline subtracted using fluorescence at 0 hr.

### 3D microfluidic culture of endothelial networks

Commercial microfluidic devices (IdenTx, AIM Biotech, Singapore), consisting of three parallel microchannel segments separated by pillars were used. 10 μL of Matrigel solution (1 mg/ml, Corning, UK), 5 μL of fibrinogen (6 mg/ml, Sigma, UK), and 5 μL of thrombin (4 units/ml, Sigma, UK) were mixed. Then, 50×10^4^ HUVECs were resuspended in 20 μL of the mixed solution and immediately injected into the central microchannel segment. All the cell seeding procedures were performed with devices placed on ice. Seeded chips were then incubated at 37 °C for 30 min to solidify the gel, before filling the side channels with pre-warmed media, as above. To create pressure-driven flow in the chip, one of the adjacent media channels was filled with 200 μL of the media, and the other channel was filled with 100 μL of the media. The chips were kept in the incubator (37 °C/5% CO_2_) for 7 days to form 3D endothelial networks, and media was changed every 24 hr. LEVs, generated and isolated as previously from MDA-MB 231 cells, were enumerated via flow cytometry. The generated LEVs were incubated with the developed 3D endothelial networks under pressure-driven flow for 24 hr at a cell: LEV ratio of 3:1. Then, 3D endothelial networks were fixed and stained for VE-cad, as described previously. Fluorescent images were acquired using a confocal microscope and cell-free area and fluorescence intensity were calculated with ImageJ (3 biological replicates and 3 technical replicates).

Perfusion studies were conducted by diluting 20 μL of 1 μm fluorescent carboxylate-modified microspheres (Fluospheres, Invitrogen) 1:100 in media. This solution was injected into one side channels of the endothelial network-laden IdenTx devices to initiate flow by hydrostatic pressure. Confocal microscopy was employed as previously to capture images of the microparticles and the networks. Quantitative analysis of microparticle colocalization with the VE-cadherin stained endothelial networks was performed using ImageJ software (3 biological replicates and 3 technical replicates).

### Monocyte differentiation

CM was collected from untreated HUVECs or HUVECs cocultured with LEVs (derived from MDA-MB 231 cells as above), as described above. 15×10^3^ THP-1 monocytes were transferred into wells of a 96-well plate and 100% of 200 μl of CM from above conditions was added for 30 hr. Additional experimental conditions included the addition of either 5×10^3^ LEVs or CM collected from LEVs on monocytes for 30 hr. Untreated monocytes were used as a negative control. After 30hr, media was removed and monocytes were stained via immunocytochemistry for CD206 and TNF-a, as described above. Multifluorescent and brightfield images were obtained at 20x magnification, using the confocal microscope. Alternatively, 10×10^4^ THP-1 monocytes were added in wells of a 24-well plate and all above experimental conditions were repeated. 33×10^3^ purified LEVs either from MDA-MB231 cells or siTGFb-transfected MDA MB-231 cells were co-cultured with monocytes, as above. 500 μl from each sample was collected and analyzed via flow cytometry as described above. The frequency of CD206+ events was calculated to estimate the polarization of monocytes towards M2 macrophages.

### Monocyte adhesion

15×10^3^ monocytes were stained with a CMTPX red Cell Tracker and were then transferred in wells of a 96 well plate. Then, monocytes were treated independently with 5×10^3^ purified LEVs (derived from MDA-MB 231 cells as above), with 100% CM collected from LEVs and 100% CM collected from MDA-MB 231 cells (as described above) for 30 hr. Untreated monocytes were used as a negative control. Fluorescent images of each well were obtained before and after media removal (removing non-adhered monocytes) and the percentage of remaining adhered cells was calculated.

Alternatively, 15×10^3^ HUVECs were added in wells of a 96-well plate and left to grow for 1 day. In the meantime, 30×10^3^ THP-1 monocytes were cultured in wells of a 48-well plate, left to grow for 1 day and then stained with a CMFDA green cell tracker. HUVECs or THP-1 monocytes were treated with 5×10^3^ or 10×10^3^ purified LEVs (derived from MDA-MB 231 cells as above) for 30hr, respectively. Negative control of both untreated HUVECs and untreated THP-1 monocytes were used. After 30hr, media was removed both from HUVECs and monocytes to remove non-internalized LEVs and four experimental conditions were performed: a) untreated THP-1 monocytes were added to untreated HUVECs monolayers, b) untreated THP-1 monocytes were added to LEV-pre-treated HUVECs monolayers, c) LEV-pre-treated THP-1 monocytes were added to untreated HUVECs monolayers, and d) LEV-pre-treated monocytes were added to LEV-pre-treated HUVECs monolayers. Using a Varioskan flash plate reader, the green fluorescence of monocytes at 0 hr was measured at 492/517 nm values of peak excitation/emission. After 4hrs of co-culture, the media was removed from each well and fresh media was added. Fluorescence of monocytes was measured by plate reader as described above. Cell-free media was used for normalization and the percentage of remaining fluorescence after media removal (removing non-adhered monocytes) was estimated.

### Monocyte proliferation assay

12×10^3^ monocytes were added in each well of a 96-well plate and left to grow for 1 day. Then, they were treated independently with 4×10^3^ purified LEVs (derived from MDA-MB 231 cells as above) or 100% CM collected from LEVs (as described above) for altogether 48 hrs. Untreated monocytes were used as a negative control. To evaluate monocyte proliferation the MTT cell proliferation assay kit (Abcam, UK) was used, per manufacturer’s instructions. The absorbance at 590 nm of each well was read using a Varioskan flash plate reader and normalized as described elsewhere.

### Graphs & Statistical Analyses

All data were presented as mean ± standard deviation (SD). Unless stated otherwise, all experiments were performed independently at least three times (n=3) for each experimental condition. Data were plotted graphically and analyzed using GraphPad Prism software (version 9.0). Significance analyses were done by applying either two-tailed student’s t-test for comparisons between two different experimental groups or one-way analysis of variance (Turkey’s multiple comparisons or Bonferonni) between three or more experimental groups (95% CI). P values ≥ 0.05 were considered as not significant. P values below this value were considered as significant (0.01 to 0.05, single asterisk*), very significant (0.001 to 0.01, double asterisk **), extremely significant (0.0001 to 0.001, triple asterisk *** or < 0.0001, quadruple asterisk ****). Illustration was prepared using Biorender.

## Acknowledgements

We would like to thank all the patients for their blood donations. We would like to acknowledge the help of the CRUK MI Visualization, Irradiation & Analysis Facility’ and Henry Banks for his assistance. Patient recruitment was supported by the National Institute for Healthcare Research (NIHR) Manchester Biomedical Research Centre, the NIHR Manchester Clinical Research Facility at The Christie Hospital and the CRUK Lung Cancer Centre of Excellence. Sample collection was undertaken through the molecular mechanisms underlying chemotherapy resistance, therapeutic escape, efficacy, and toxicity improving knowledge of treatment resistance in patients with lung cancer or CHEMORES protocol. We acknowledge the ICR proteomics core facility for the performance of the proteomic experiment. We would like to thank the CRUK Convergence Science Centre for funding the CRUK Microfabrication and Prototyping Facility and Tweety Tang, Florent Seichepine and Shahrzad Forouzanfar for their help.

## Author contributions

A.V. and S.H.A. conceptualized the project. A.V. designed all experiments, fabricated microfluidic devices and ran all microfluidic experiments, developed methodologies on LEVs generation and purification, characterized LEVs, performed LEVs co-culture experiments, analyzed data and conducted statistical analyses. B.C. conducted western blotting and G.S. assisted with Western blotting and conducted centrifugal force experiments. S.B.S. performed the 3D vascular network formation experiments and analyzed data. S.A. conducted proteomics experiments and S.A. & P.H. conducted proteomics analysis. K.S. and M.C. processed patient and clinical samples. F.B. supervised clinical sample collection. K.S., S.B.S., K.L.S. and C.D. coordinated CDX experiments. K.S., K.L.S. and C.D. coordinated primary CTC experiments. A.V. performed primary CTC experiments on microfluidics and identified fragments in patient samples. B.C. performed western blot. GS performed the cell centrifugation experiment on LEV biogenesis. T.M.P. performed cryoEM experiments and analyzed data. M.A., A.Z. and D.V assisted with macrophage, monocyte and endothelial experiments, respectively. A.C. assisted with cluster experiments. S.H.A. conducted computational fluid dynamics and supervised the overall project direction. A.V and S.H.A. wrote the manuscript with edits from all authors.

## Data and materials availability

The mass spectrometry proteomics data have been deposited to the ProteomeXchange Consortium via the PRIDE ^47^ partner repository with the dataset identifier PXD050444. All other data are available in the main text or in the supplementary materials.

## Supplementary Information

Supplementary Figures S1 to S24

Supplementary Tables S1 to S8

Supplementary Movies S1 to S8

