## Supplementary Information for "Circulating tumor cells shed shearosome extracellular vesicles in capillary bifurcations that activate endothelial and immune cells"

Angelos Vrynas et al.

* Corresponding author: s.au<att>imperial.ac.uk

**This file includes:**

Figs. S1 to S24

Tables S1 to S8

**Other Supplementary Materials for this manuscript include the following:**

Movies S1 to S8

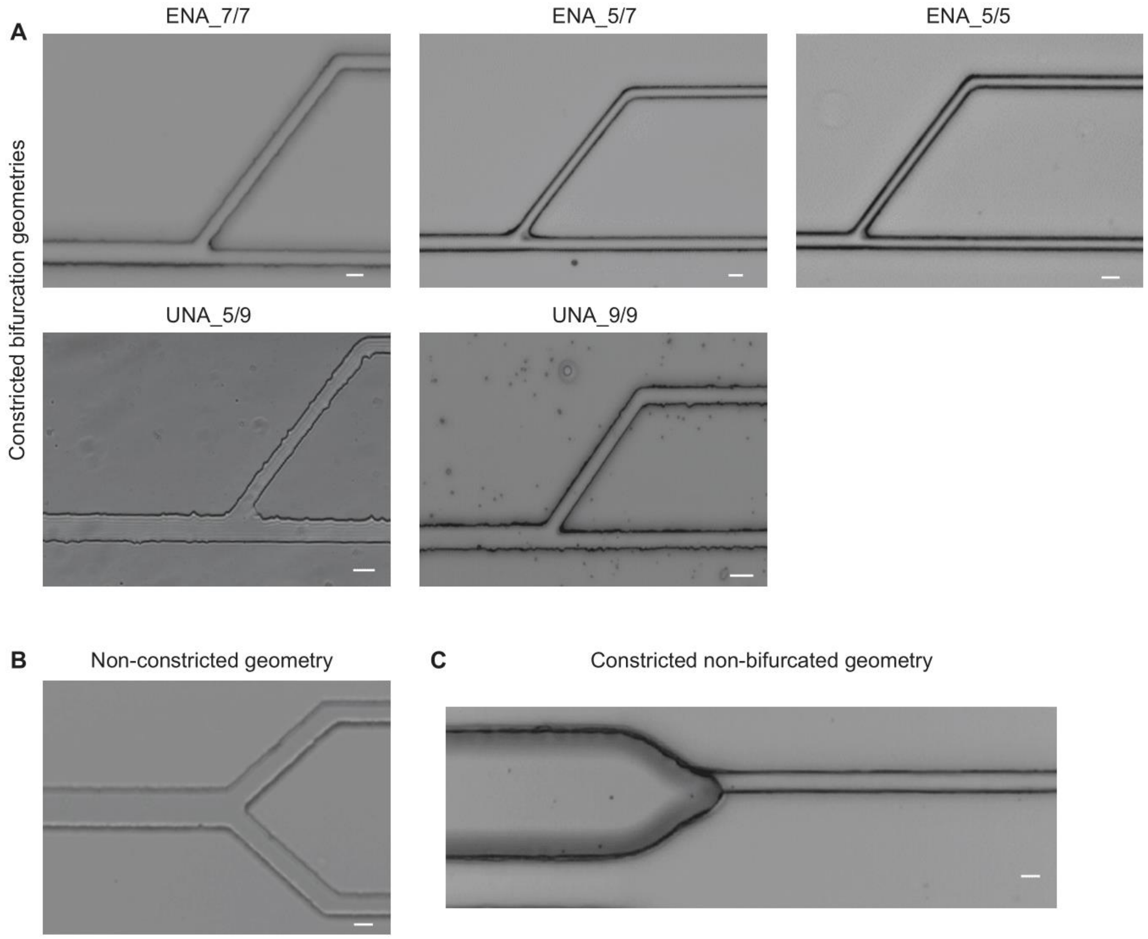


**Fig. S1. Microfluidic devices**. (**A**) Brightfield images of various constricted bifurcation variants. Scale bar: 10 µm (excluding for UNA_9/9: Scale bar: 15 µm). (**B**) Brightfield image of a non-constricted geometry. Scale bar: 20 µm. (**C**) Brightfield image of a constricted non-bifurcated geometry. Scale bar: 7 µm.

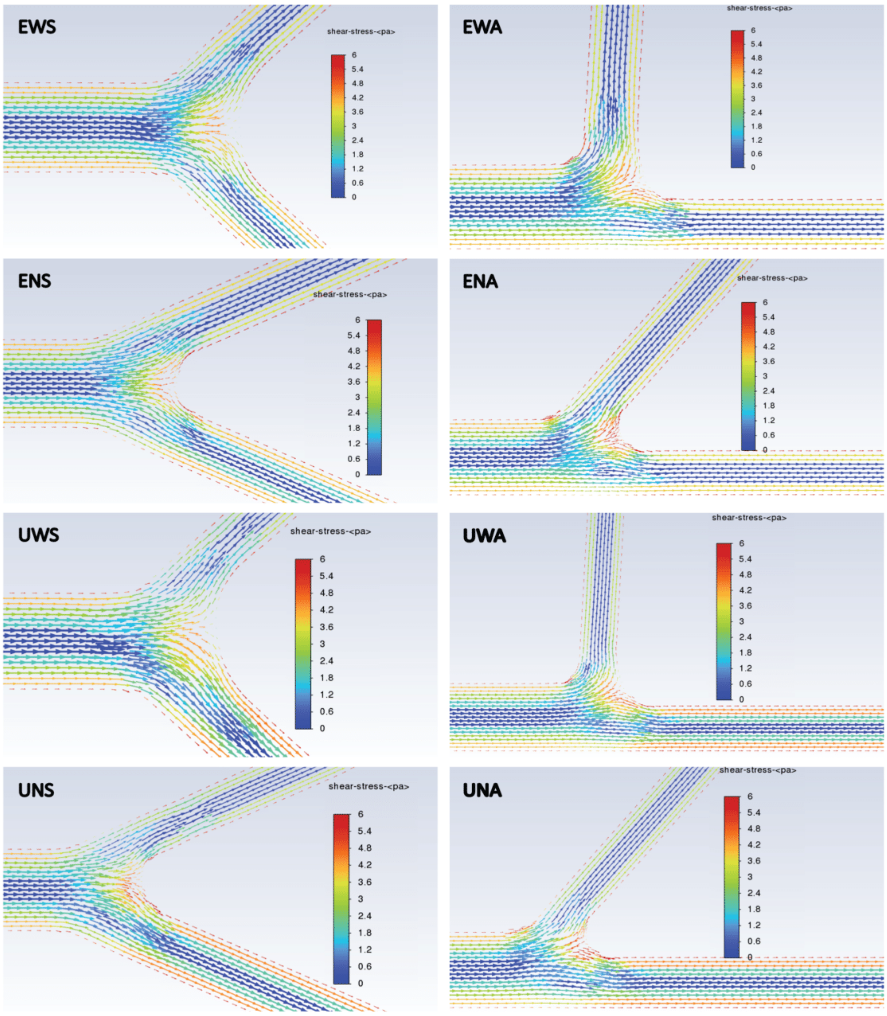


**Fig. S2. Computational fluid dynamics simulations**. CFD simulation of 8 bifurcation variants (EWS, EWA, ENS, ENA, UWS, UWA, UNS and UNA) conducted in ANSYS Fluent showing colorized heat map of fluid shear stress (Pa) velocity vectors.

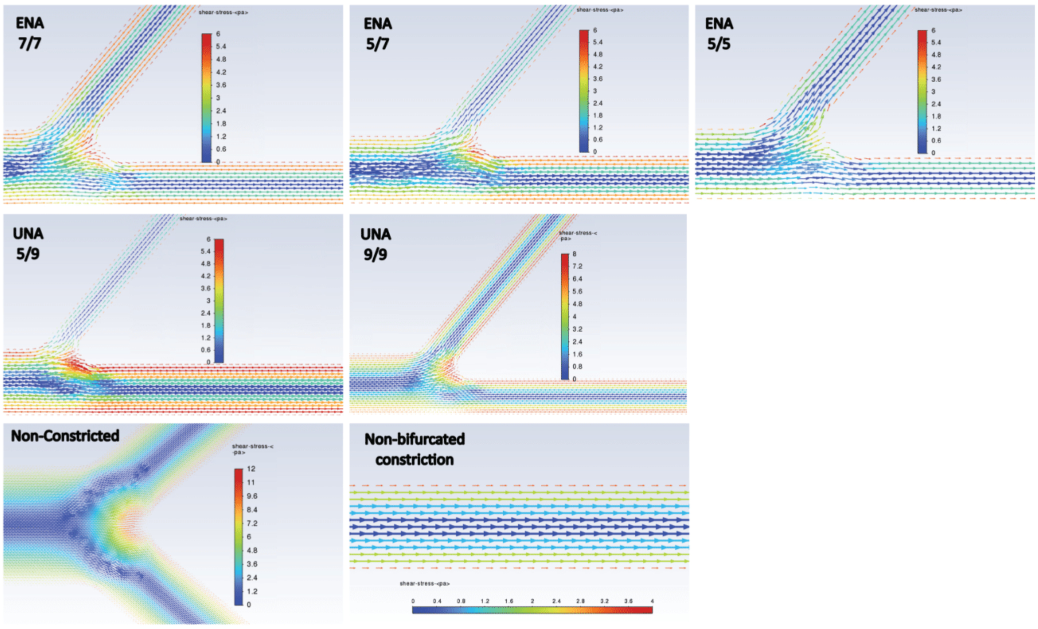


**Fig. S3. Computational fluid dynamics simulations**. CFD simulation of 5 supplementary bifurcation variants (ENA7/7, ENA5/7, ENA5/9, UNA5/9, UNA9/9), non-constricted bifurcation, and constricted non-bifurcated geometries conducted in ANSYS Fluent showing colorized heat map of fluid shear stress (Pa) velocity vectors.

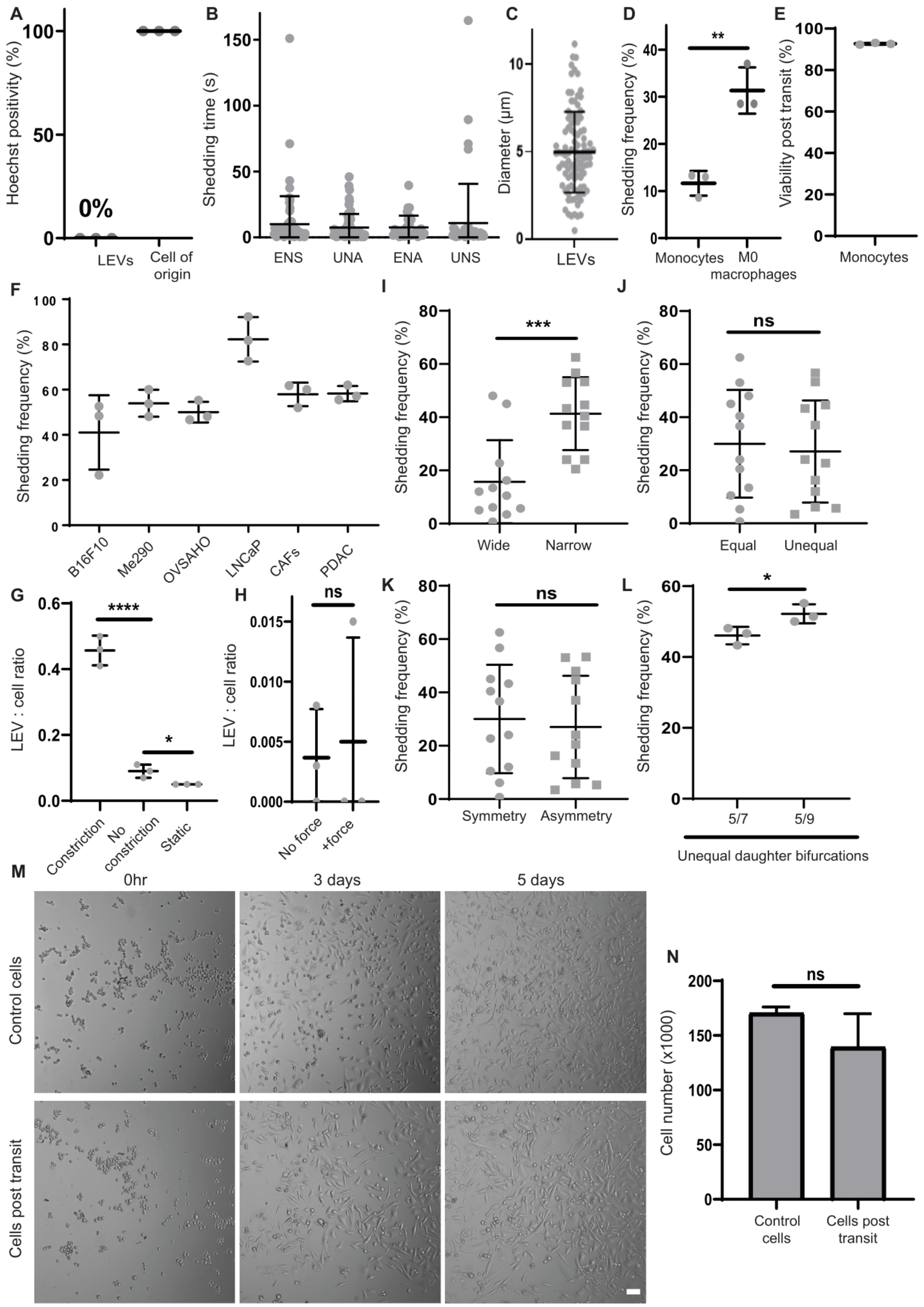

A
Hoechst positivity (%)
0%
LEVs
Cell of origin
B
Shedding time (s)
ENS
UNA
ENA
UNS
C
Diameter (µm)
LEVs
D
Shedding frequency (%)
**
Monocytes
M0 macrophages
E
Viability post transit (%)
Monocytes
F
Shedding frequency (%)
B16F10
Me290
OVSAHO
LNCaP
CAFs
PDAC
I
Shedding frequency (%)
***
Wide
Narrow
J
Shedding frequency (%)
ns
Equal
Unequal
G
LEV : cell ratio
****
*
Constriction
No constriction
Static
H
LEV : cell ratio
ns
No force
+force
K
Shedding frequency (%)
ns
Symmetry
Asymmetry
L
Shedding frequency (%)
*
5/7
5/9
Unequal daughter bifurcations
M
0hr
3 days
5 days
Control cells
Cells post transit
N
ns
Cell number (x1000)
Control cells
Cells post transit

**Fig. S4. Biomechanical investigation of LEV biogenesis.** (**A**) Percentage of Hoechst-positive MDA-MB 231 cells or their derived LEVs after transit through bifurcation variant ENA using live imaging (n=3). (**B**) MDA-MB 231 cells' shedding time in 4 different capillary bifurcation variants (n=66 for ENS, n=61 for UNA, n=31 for ENA, n=46 for UNS). Each data point represents a single cell. (**C**) Diameter of LEVs counted using CellProfiler (n=101). (**D**) Shedding frequency (%) of THP-1 monocytes and THP-1 derived macrophages (n=3). (**E**) Viability (%) of THP-1 monocytes post transit capillary bifurcations (n=3). (**F**) Percentage of shedding frequency for melanoma (B16F10, Me290), ovarian (OVSAHO), prostate (LNCaP), pancreatic (PDAC) cancer cell lines and primary cancer-associated fibroblasts during live imaging (n=3). (**G**) LEV : cell ratio, enumerated post MDA-MB 231 cells transit in bifurcation variant UNA_5/9 or non-constricted geometry or static conditions (n=3). (**H**) LEV : cell ratio, enumerated for MDA-MB231 cells that were centrifuged at 8000 RCF for 3 mins. Non-centrifuged cells were used as control (n=3). (**I-L**) Percentage of MDA-MB 231 cells shedding frequency in different bifurcation variants (**I**) wide vs narrow (n=12) (**J**) equal vs unequal (n=12) (**K**) symmetry vs asymmetry (n=12) and (**L**) unequal capillary bifurcation geometries UNA_5/7 vs UNA_5/9 during live imaging (n=3). (**M**) Brightfield images of control cells or cells post transit (MDA-MB 231 cells) at 0hr, 3 days and 5 days. Scale bar: 100 µm. (**N**) Quantification of MDA-MB 231 cell number 72 hrs post transit through bifurcation variant ENA versus control non-transiting cells.

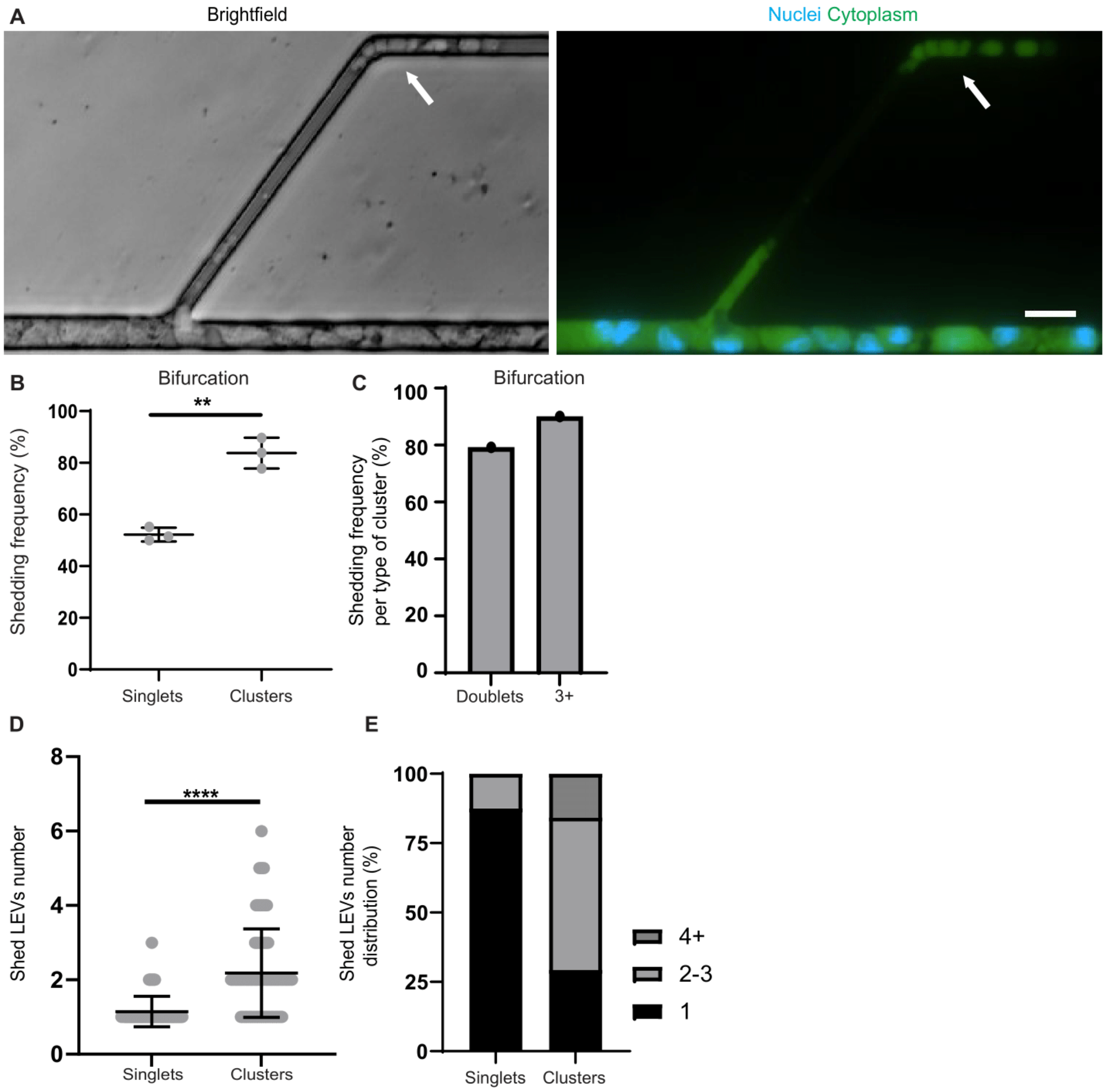


**Fig. S5. Investigation of LEV biogenesis from tumor cell clusters.** (**A**) Brightfield & multifluorescent image of trapped large cluster of MDA-MB231 cells shedding multiple LEVs (arrow) in bifurcation variant UNA_5/9. Cytoplasm was stained with CMFDA cell tracker (green) and nuclei with Hoechst-33342 (blue). Arrow depicts the multiple shed LEVs. Scale bar: 20 µm. (**B**) Percentage of either single MDA-MB 231 cells or clusters that shed at least once during live imaging in bifurcation variant UNA_5/9. (n=3). (**C**) Percentage of MDA-MB 231 cell clusters that shed per type of cluster, 2-cell cluster (doublet) (n=101) or 3-cell cluster and higher (n=40). Data were pooled together from cell transit in bifurcation variants UNA_5/9 and UNA_7/7. (**D**) Number of LEVs shed per single MDAMB 231 cell (n=48 singlets) or cluster (n=66 clusters) in bifurcation variant UNA_5/9. (**E**) Distribution of (**D**) per number of shed LEVs.

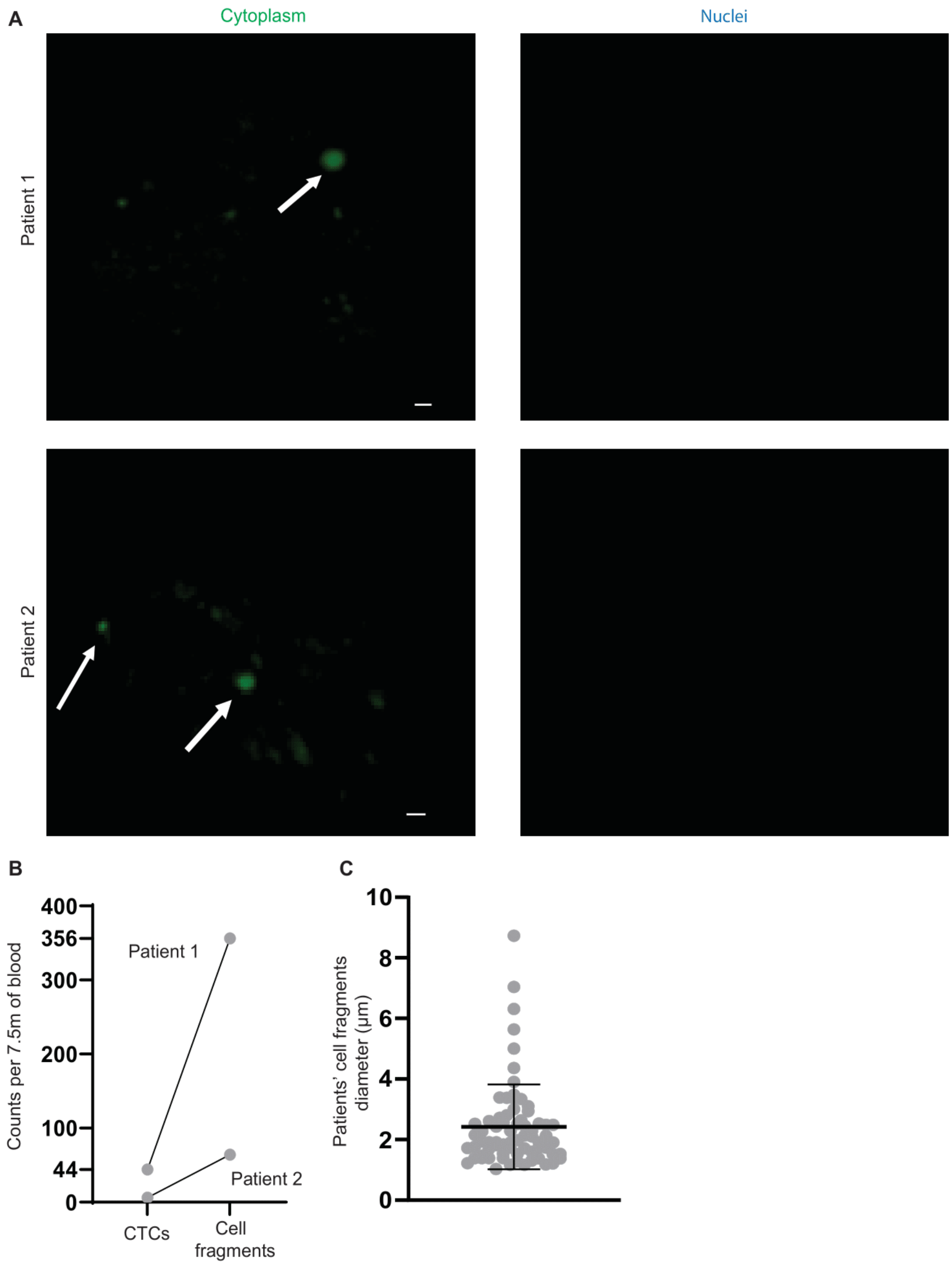


**Fig. S6. Isolation of patient cellular fragments.** (**A**) Multifluorescent images of isolated cell fragments from patients with small cell lung carcinoma (patient 1: top & patient 2: bottom), stained with CMFDA for cytoplasm (green) and Hoechst-33342 for nuclei (blue) (n=2 patients). Arrows indicate cellular fragments. Scale bar: 5 µm (Patient 1) & Scale bar: 4 µm (Patient 2). (**B**) CTCs and cell fragments numbers from patient blood (n=2 patients). (**C**) Diameter (µm) of patient-isolated cell fragments (n=70 fragments).

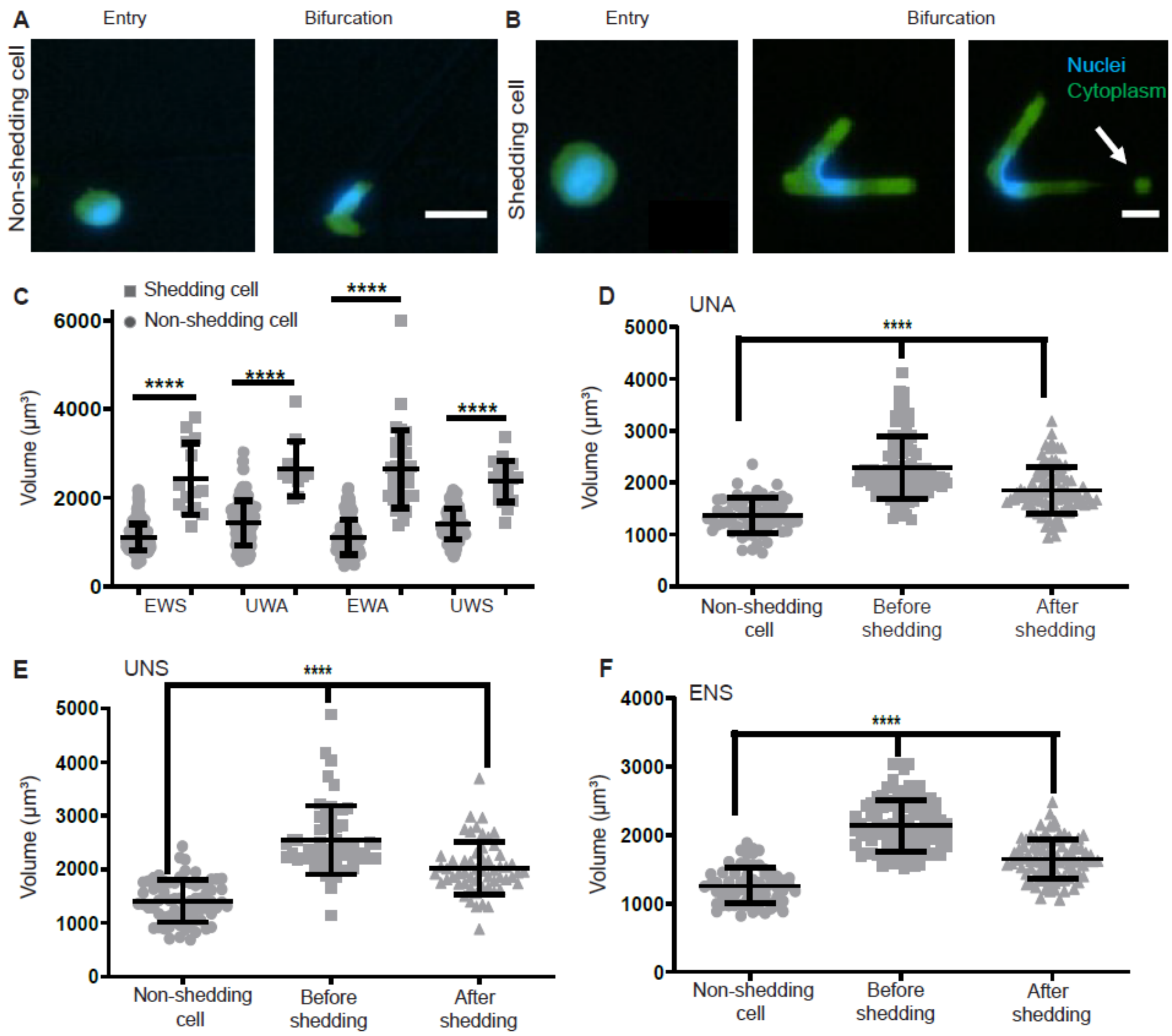


**Fig. S7. Tumor cell size promotes shedding.** (**A-B**) 20x multifluorescent images of non-shedding (A) or shedding (**B**) (arrow) MDA-MB 231 cells. Cytoplasm was stained with Calcein-AM (green) and nuclei with Hoechst-33342 (blue). Scale bar: 20µm. (**C**) Cytoplasmic volume ($\mu m^3$) of non-shedding MDA-MB 231 cells and MDA-MB 231 cells that eventually shed, during transit in 4 bifurcated capillary geometries (EWS, UWA, EWA, UWS). (**D**) Cytoplasmic volume ($\mu m^3$) of non-shedding MDA-MB 231 cells (n=62) and MDA-MB 231 cells that eventually shed (before and after shedding) (n=97), during transit in bifurcation variant UNA. (**E**) Cytoplasmic volume ($\mu m^3$) of non-shedding MDA-MB 231 cells (n=71) and MDA-MB 231 cells that eventually shed (before and after shedding) (n=57), during transit in bifurcation variant UNS. (**F**) Cytoplasmic volume ($\mu m^3$) of non-shedding MDA-MB 231 cells (n=68) and MDA-MB 231 cells that eventually shed (before and after shedding) (n=92), during transit in bifurcation variant ENS. (**C-F**) Each data point represents a single cell.

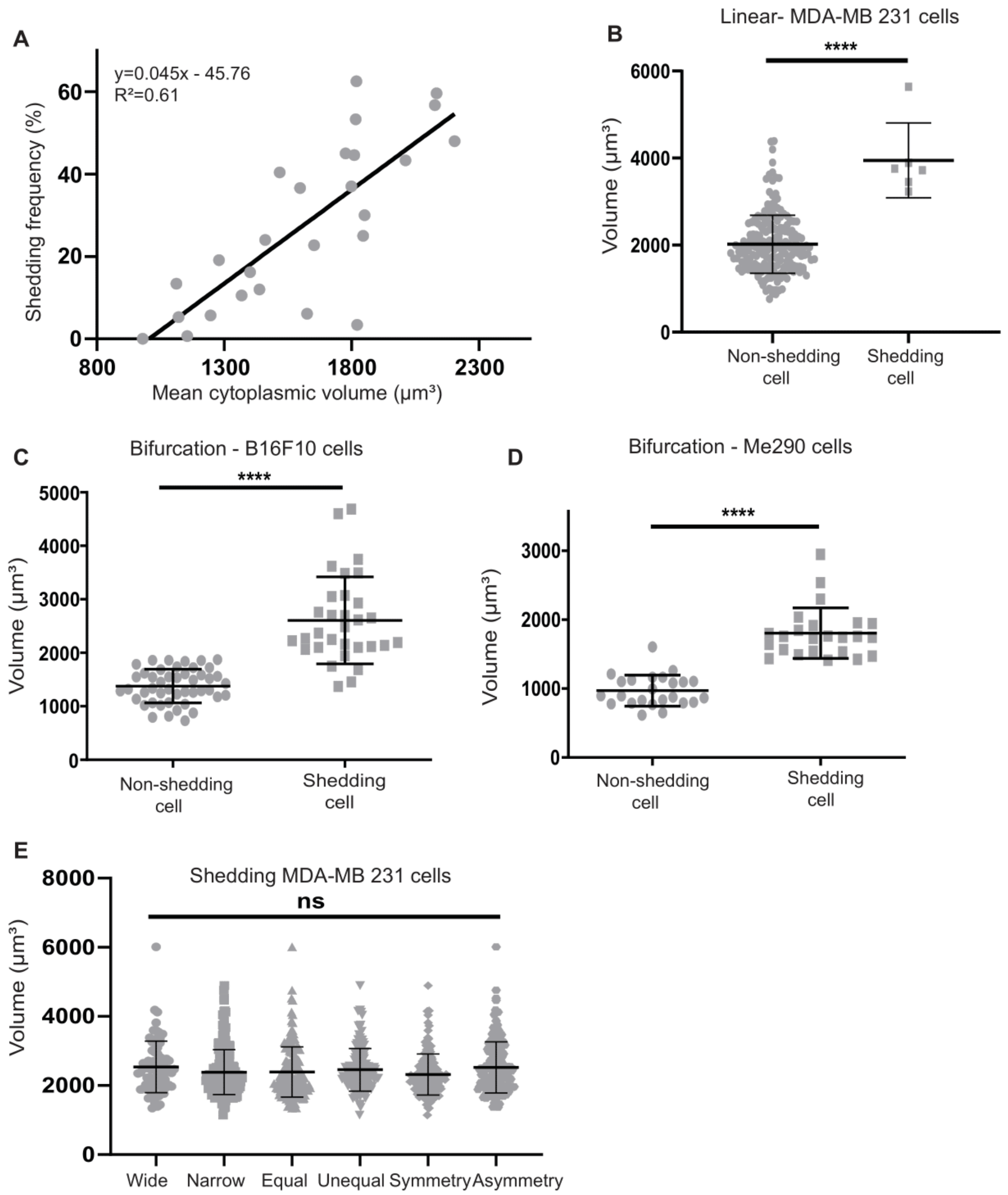


**Fig. S8. Influence of cell size on shedding during transit in various microfluidic variants.** (**A**) Graph plot of shedding frequency (y axis) and mean cytoplasmic volume (µm$^3$) (x axis) (n=26). Data were pooled together from various bifurcation devices. (**B-D**) Each data point represents a single cell. (**B**) Cytoplasmic volume (µm$^3$) of non-shedding MDA-MB 231 cells (n=198) and MDA-MB 231 cells that eventually shed (n=6), during transit in constricted non-bifurcated (linear) geometry. (**C-D**) Cytoplasmic volume (µm$^3$) of non-shedding cells (n=46 for B16F10 (**C**), n= 24 for Me290 (**D**)) or shedding cells (n=31 for B16F10 (**C**), n= 24 for Me290 (**D**)), during transit in bifurcation variant UWA (**C**) and ENA (**D**). (**E**) Cytoplasmic volume (µm$^3$) of shedding MDA-MB 231 cells, per bifurcation design parameter (n=79 for wide, n=244 for narrow, n=170 for equal, n=153 for unequal, n=159 for symmetry, n=164 for asymmetry.

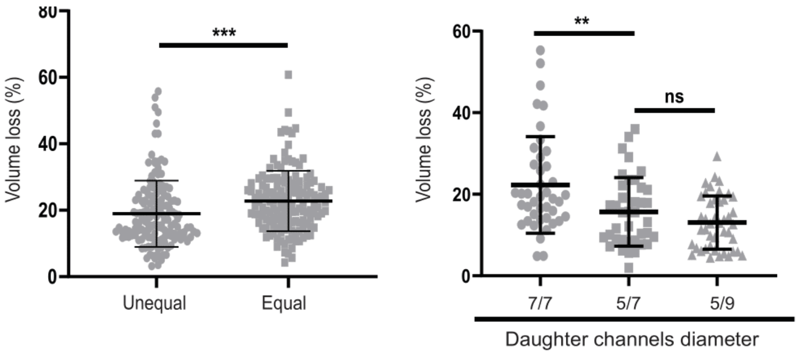


**Fig. S9. Tumor cell volume losses post shedding.** (**A-B**) Each data point represents a single cell. (**A**) Percentage of volume loss of post shedding MDA-MB 231 cells during transit in unequal bifurcation variants (UNA, UNS) (n=154) or equal bifurcation variants (ENA, ENS) (n=148 cells). (**B**) Percentage of volume loss of post shedding MDA-MB 231 cells during transit in 3 independent bifurcation variants (n=42 cells for ENA_7/7, n=39 cells for UNA_5/7, n= 44 cells for UNA_5/9).

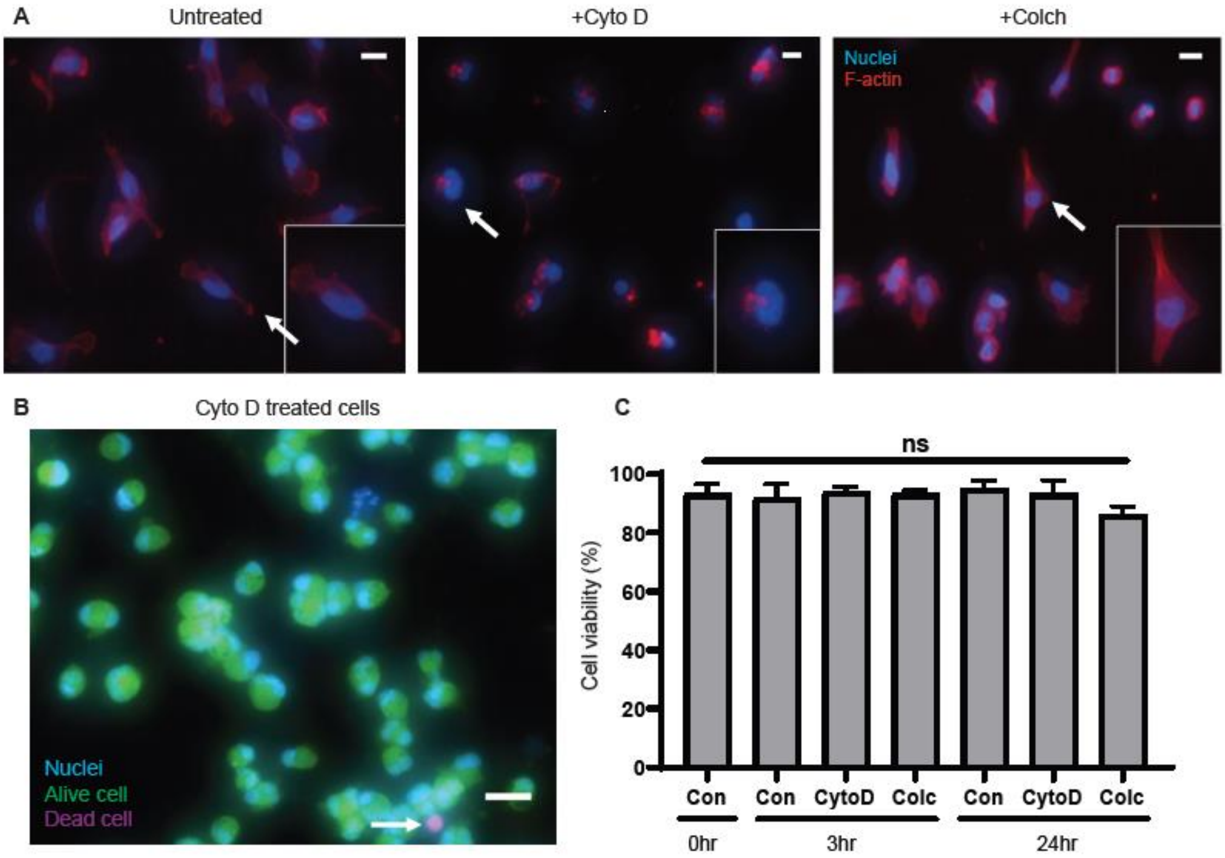


**Fig. S10. Validation of F-actin polymerization inhibitors or promoters.** (**A**) Multifluorescent images of untreated (left), Cytochalasin D (Cyto-D) treated (middle), or Colchicine (Colch) treated MDA-MB 231 cells. F-actin was stained with Phalloidin (red) and nuclei with Hoechst-33342 (blue). Arrows indicate the zoomed cell (inset). Scale bar: 20 µm. (**B**) Multifluorescent image obtained from live/dead assay. Live cells were stained with Calcein-AM (green), dead cells with Propidium Iodide (red) and nuclei with Hoechst-33342 (blue). Arrow indicates a dead cell. Scale bar: 50µm. (**C**) Quantification of live/dead assay for untreated cells (Con), Cyto-D treated or Colch-treated cells at 3 or 24hrs (n=3).

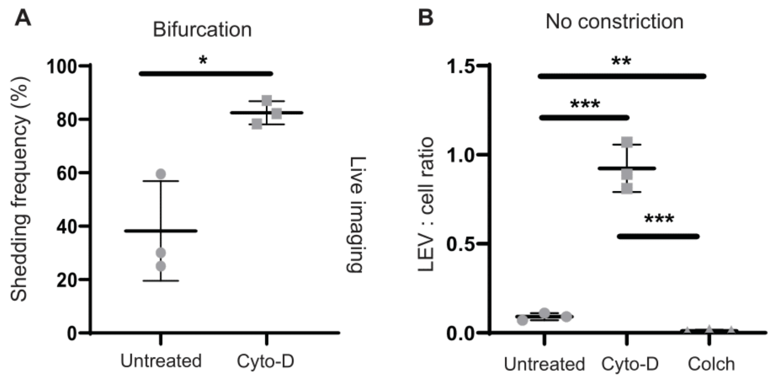


**Fig. S11. Investigation of F-actin polymerization inhibitors or promoters on single cell shedding.** (**A**) Shedding frequency (%) of untreated or Cyto-D treated MDA-MB 231 cells during transit in bifurcation variant ENA (live imaging) (n=3). (**B**) LEV : cell ratio of untreated, Cyto-D or Colch-pre-treated MDA-MB 231 cells enumerated post transit in non-constricted geometry (n=3).

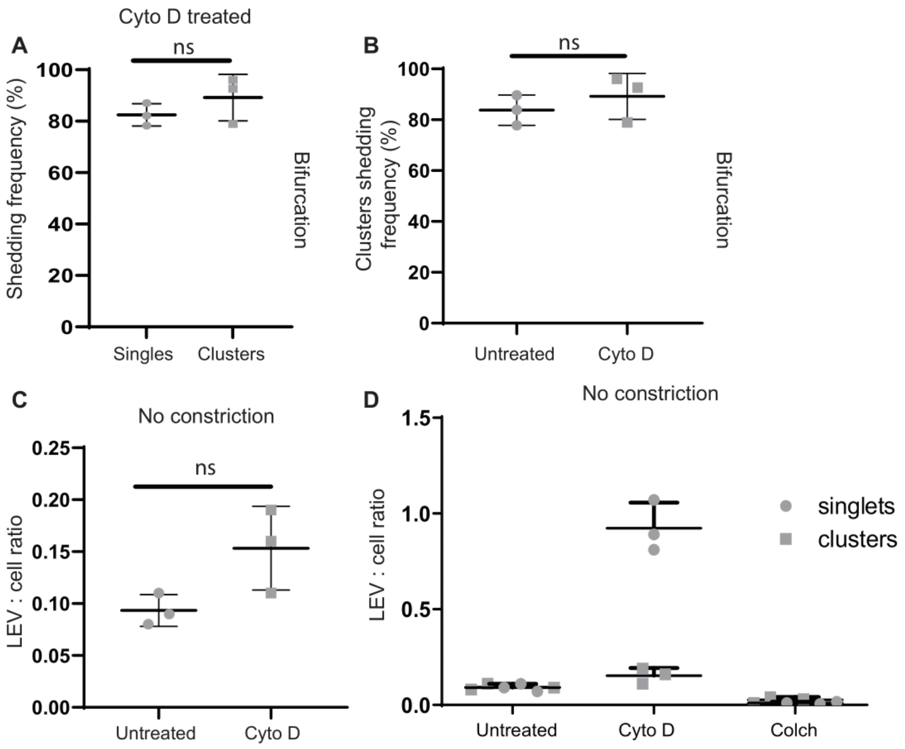


**Fig. S12. Investigation of F-actin polymerization inhibitors or promoters on clustered cells shedding.** (**A**) Percentage of Cyto-D-treated clusters or Cyto-D treated single MDA-MB 231 cells that shed at least once during transit in bifurcation variant UNA_5/9 (live imaging) (n=3). (**B**) Percentage of Cyto-D treated clusters or untreated clusters (MDA-MB 231 cells) that shed at least once during transit in bifurcation variant UNA_5/9 (n=3). (**C**) LEV : cell ratio of Cyto-D treated or untreated clusters enumerated post transit in non-constricted geometry (n=3). (**D**) LEV : cell ratio of Cyto-D treated, Colch-treated and untreated single cells or clusters, enumerated post transit in non-constricted geometry (n=3).

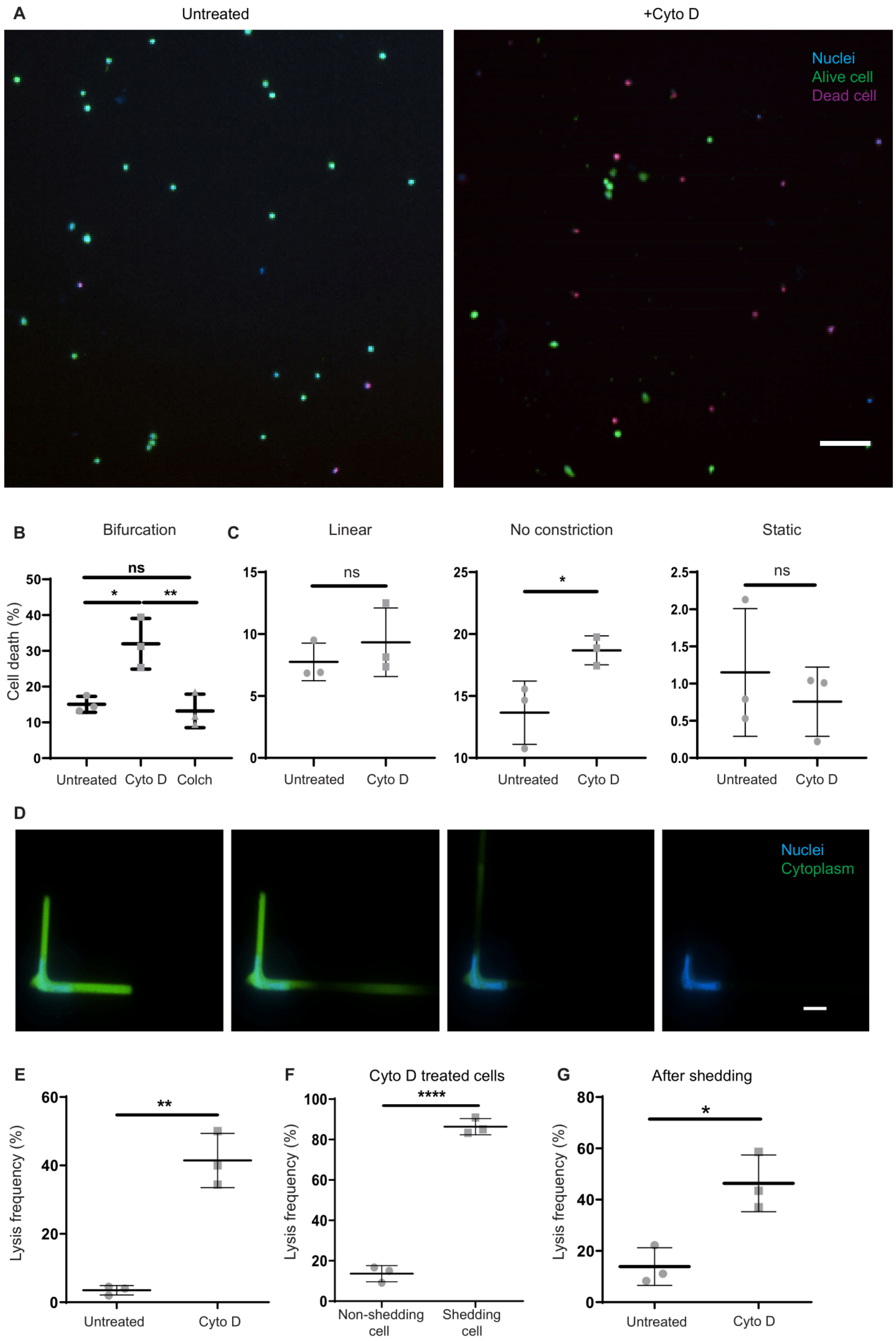
A
Untreated
+Cyto D
Nuclei
Alive cell
Dead cell
B
Bifurcation
Cell death (%)
ns
*
**
Untreated
Cyto D
Colch
C
Linear
ns
Untreated
Cyto D
No constriction
*
Untreated
Cyto D
Static
ns
Untreated
Cyto D
D
Nuclei
Cytoplasm
E
Lysis frequency (%)
**
Untreated
Cyto D
F
Cyto D treated cells
Lysis frequency (%)
****
Non-shedding cell
Shedding cell
G
After shedding
Lysis frequency (%)
*
Untreated
Cyto D

**Fig. S13. Investigation of tumor cell viability in microfluidic variants**. (**A**) Multifluorescent images of a live/dead assay for untreated (left) or Cyto-D treated (right) MDA-MB 231 cells collected post transit from capillary bifurcations. Live cells were stained with Calcein-AM (green), dead cells with Propidium Iodide (red) and nuclei with Hoechst-33342 (blue). Scale bar: 100 µm.**(B)** Percentage of untreated, Cyto-D treated and Colch-treated MDA-MB231 cells that were stained dead post transit form bifurcation variant UNA_5/9 (n=3). (**C**) Percentage of untreated or Cyto-D treated cells that were stained dead post transit in constricted non-bifurcated (linear) geometry (left), non-constricted geometry (middle) or static conditions (right) (n=3). (**D**) Timelapse of a MDA-MB 231 cell trapped in bifurcation variant EWA, that shed and eventually was lysed. Cytoplasm was stained with CMFDA cell tracker (green) and nuclei with Hoechst-33342 (blue). Scale bar: 10 µm. (**E**) Percentage of untreated or Cyto-D treated MDA-MB 231 cells that were lysed during transit in bifurcation variant ENA (n=3). (**F**) Percentage of non-shedding or shedding Cyto-D treated MDA-MB 231 cells that were lysed during transit in bifurcation variant ENA (n=3). (**G**) Percentage of untreated or Cyto-D treated MDA-MB 231 cells that were lysed post shedding, during transit in bifurcation variant ENA (n=3).

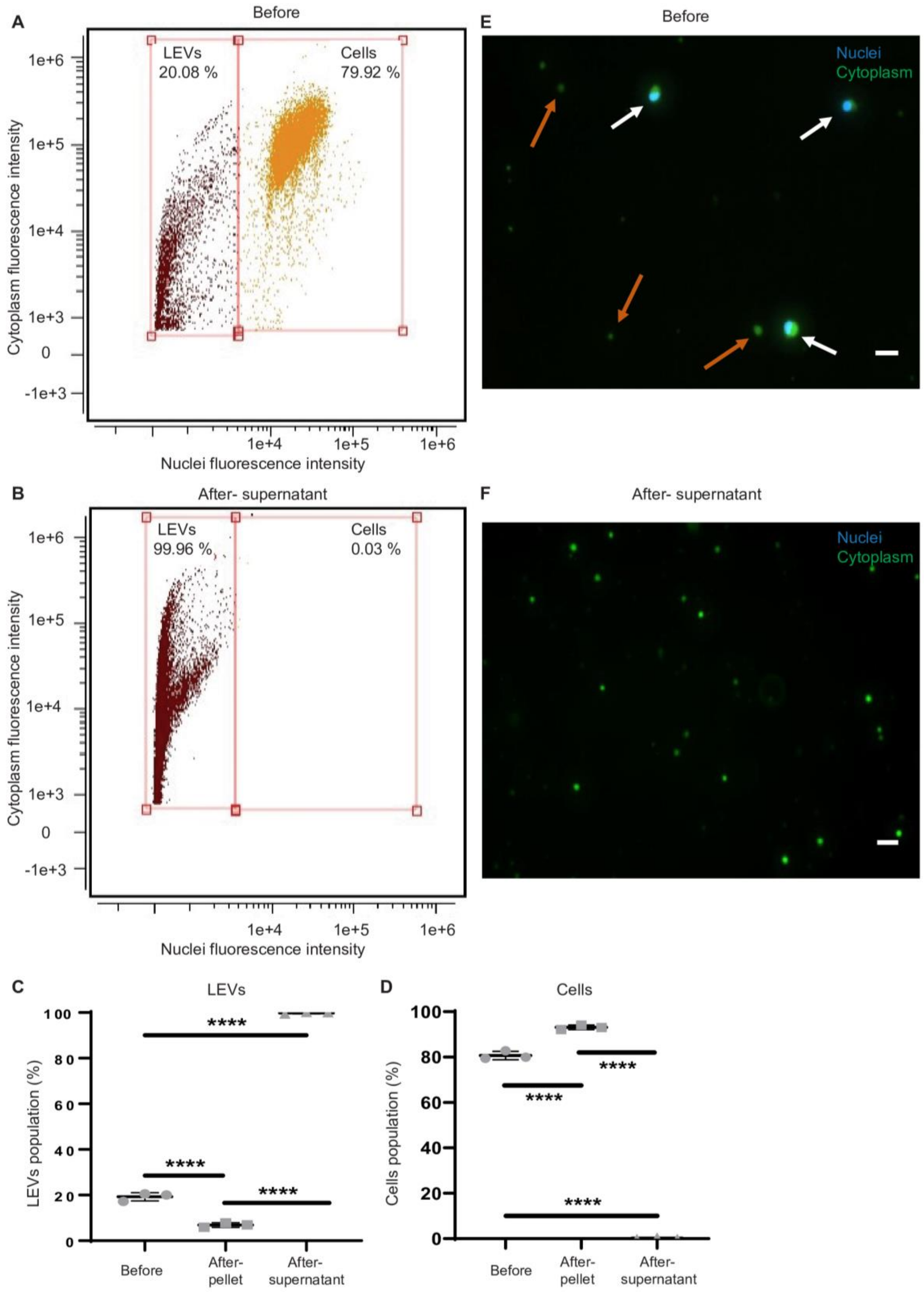
A
Before
LEVs
20.08 %
Cells
79.92 %
1e+6
1e+5
1e+4
1e+3
0
-1e+3
Cytoplasm fluorescence intensity
1e+4
1e+5
1e+6
Nuclei fluorescence intensity
B
After- supernatant
LEVs
99.96 %
Cells
0.03 %
1e+6
1e+5
1e+4
1e+3
0
-1e+3
Cytoplasm fluorescence intensity
1e+4
1e+5
1e+6
Nuclei fluorescence intensity
C
LEVs
****
****
****
100
80
60
40
20
0
LEVs population (%)
Before
After-
pellet
After-
supernatant
D
Cells
****
****
****
100
80
60
40
20
0
Cells population (%)
Before
After-
pellet
After-
supernatant
E
Before
Nuclei
Cytoplasm
F
After- supernatant
Nuclei
Cytoplasm

**Fig. S14. Separation of cells and their shed LEVs.** (**A-B**) Dot plot of gated Cytoplasm-positive MDA-MB 231 cells and their derived cytoplasm-positive LEVs, separated based on Hoechst (nuclei) intensity, before (**A**) and after centrifugation (supernatant) (**B**). (**C-D**) Percentage of LEVs (**C**) or cells (**D**) that were present in the sample before centrifugation, after centrifugation (pellet) and after centrifugation (supernatant) (n=3). (**E-F**) 20x multifluorescent images before (**E**) and after centrifugation (supernatant) (**F**). Cytoplasm was stained with CMFDA cell tracker (green) and nuclei with Hoechst-33342 (blue). Scale bar: 20 µm. White arrows indicate cells and orange arrows indicate LEVs (**E**).

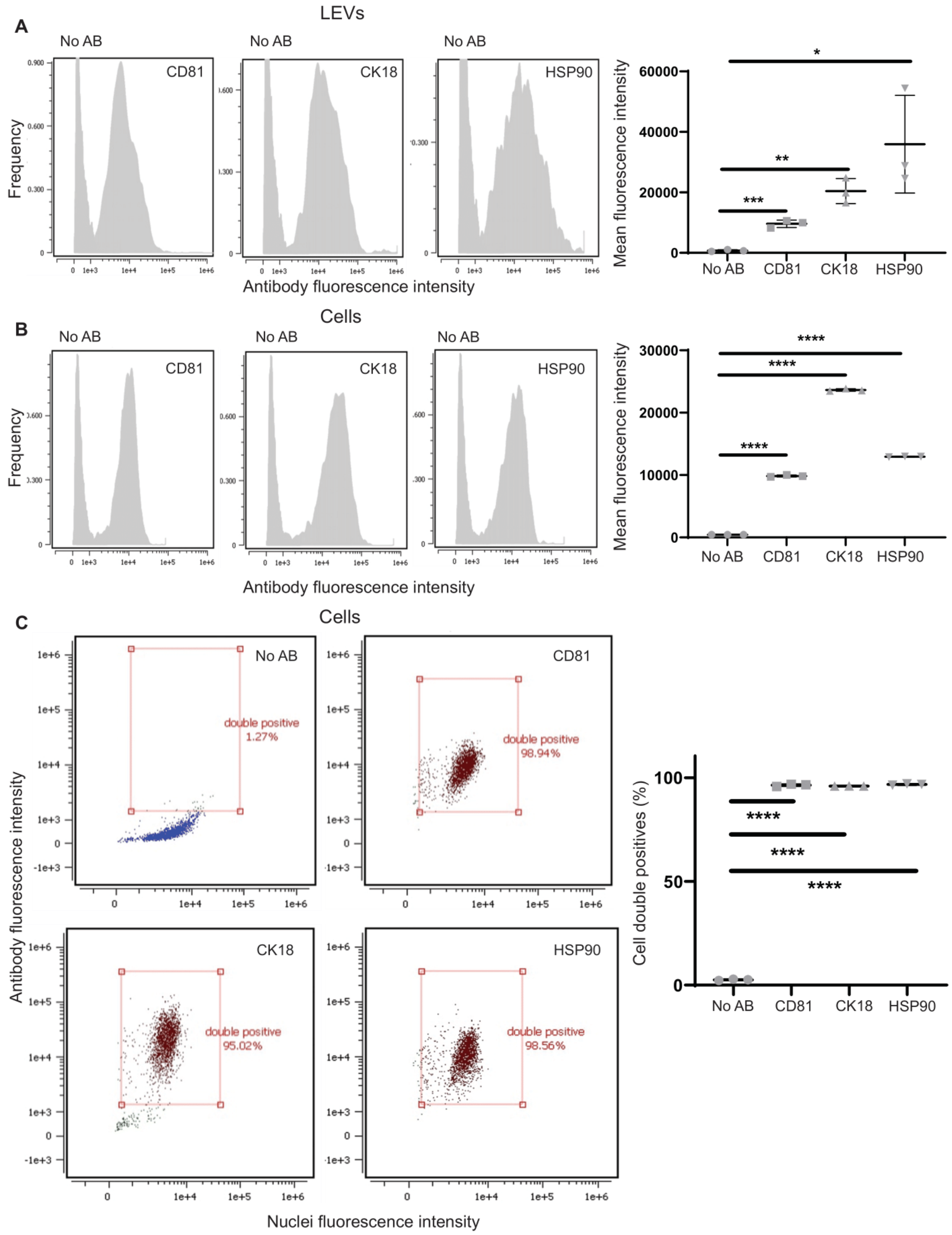

A
LEVs
No AB
CD81
CK18
HSP90
Frequency
Antibody fluorescence intensity
Mean fluorescence intensity
No AB
CD81
CK18
HSP90
B
Cells
No AB
CD81
CK18
HSP90
Frequency
Antibody fluorescence intensity
Mean fluorescence intensity
C
Cells
No AB
double positive 1.27%
CD81
double positive 98.94%
CK18
double positive 95.02%
HSP90
double positive 98.56%
Antibody fluorescence intensity
Nuclei fluorescence intensity
Cell double positives (%)

**Fig. S15. Protein inclusion on tumor cells and their shed LEVs.** (**A-C**) Data obtained from flow cytometry analysis. (**A**) Histograms (left) & quantification (far right) of the mean fluorescence intensity of antibody-stained (Cluster of differentiation 81-CD81, Cytokeratin 18-CK18 and Heat shock protein 90-HSP90) or non-antibody stained (No AB) LEVs, derived from MDA-MB 231 cells (n=3). (**B**) Histograms (left) & quantification (far right) of the mean fluorescence intensity of antibody-stained (CD81, CK18 and HSP90) or non-antibody stained (No AB) MDA-MB 231 cells (n=3). (**C**) Dot plots (left) & quantification (far right) of the percentage of double positive events for Hoechst-positive MDA-MB 231 cells that express CD81 (top right) or CK18 (bottom left) or HSP90 (bottom right). Non-antibody stained (No AB) cells were used as control (top left) (n=3). Antibody fluorescence and Hoechst fluorescence intensity are recorded in each dot plot in the y and x axes, respectively.

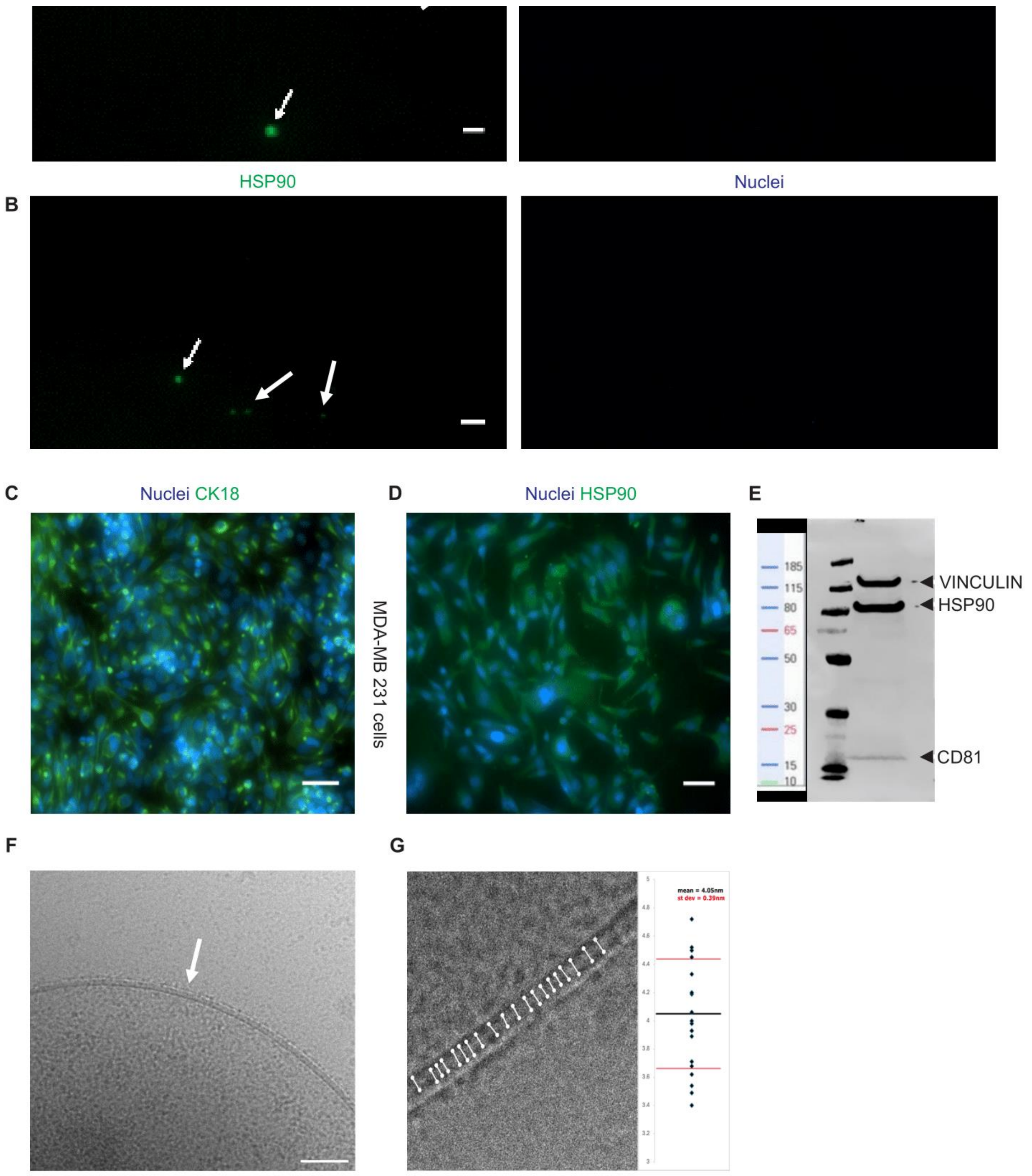
HSP90
Nuclei
B
C
Nuclei CK18
D
Nuclei HSP90
E
MDA-MB 231 cells
185
115
80
65
50
30
25
15
10
VINCULIN
HSP90
CD81
F
G
mean = 4.05nm
st dev = 0.39nm

**Fig. S16. Imaging of protein expression on tumor cells and their shed LEVs.** (**A**) Multifluorescent image of purified LEVs stained for CK18 (green) (arrows). Nuclei was stained with Hoechst-33342 (blue). Arrows indicate LEVs. Scale bar: 10 µm. (**B**) Multifluorescent image of purified LEVs stained for HSP90 (green) (arrows). Nuclei was stained with Hoechst-33342. Arrows indicate LEVs. Scale bar: 10 µm (**C**) Multifluorescent image of MDA-MB 231 cells stained for CK18 (green). Nuclei was stained with Hoechst-33342 (blue). Scale bar: 50 µm. (**D**) Multifluorescent image of MDA-MB 231 cells stained for HSP90 (green). Nuclei was stained with Hoechst-33342 (blue). Scale bar: 50 µm. (**E**) Western blot on LEVs for HSP90 and CD81 proteins. Vinculin was used as a loading control. (**F**) Transmission electron microscopy image of the lipid bilayer of an LEV produced from MDA-MB-231 cancer cell. White arrow indicates the lipid bilayer. Scale bar: 30 nm. (**G**) Quantification of the lipid bilayer width across an individual LEV (n=20).

A

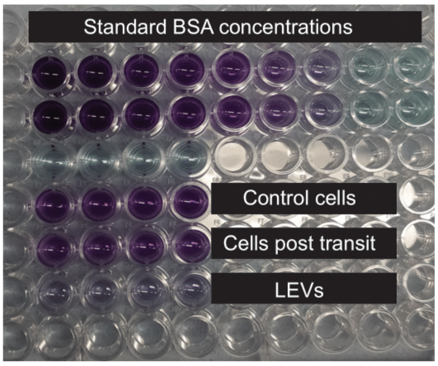


B

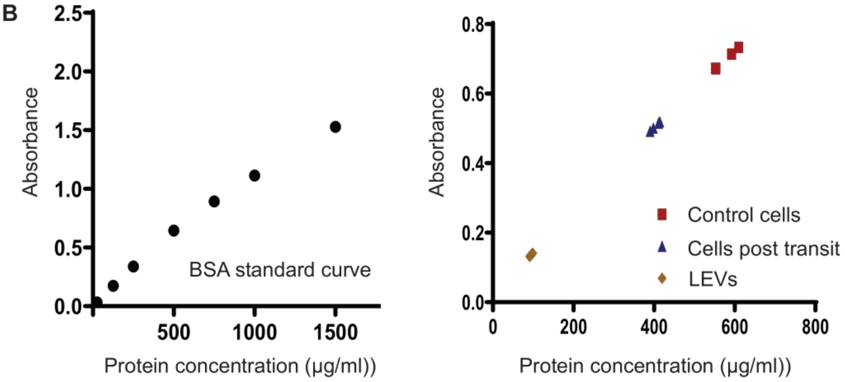


C

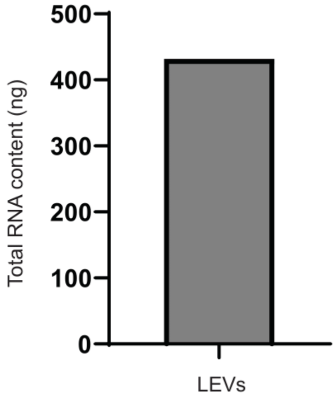


D

| Gene | CT (Sample 1) | CT (Sample 2) |
|---|---|---|
| IL-6 | 11.96 | 10.3 |
| CXCL10 | 10.94 | 8.83 |
| FN | 11.24 | 6.06 |
| ALDH1 | 10.86 | 8.99 |
| BCL2 | 8.78 | 6.38 |
| Vimentin | 9.42 | 10.01 |
| CD44 | 8.85 | 5.61 |
| RPLO | 11.27 | 10.16 |

**Fig. S17. LEVs contain proteins and RNA.** (**A**) BCA assay for estimation of protein concentration for MDA-MB 231 control cells, MDA-MB 231 cells post transit and their derived LEVs (n=4). (**B**) BSA standard curve (left) and respective data points for control cells, cells post transit and LEVs (right) (n=4). (**C**) Total RNA content (ng) measured from 2x$10^5$ LEVs (n=1). (**D**) List of mRNA molecules that were present in purified LEVs, validated via reverse transcription polymerase chain reaction with their CT values against a housekeeping gene (RPLO) (n=2).

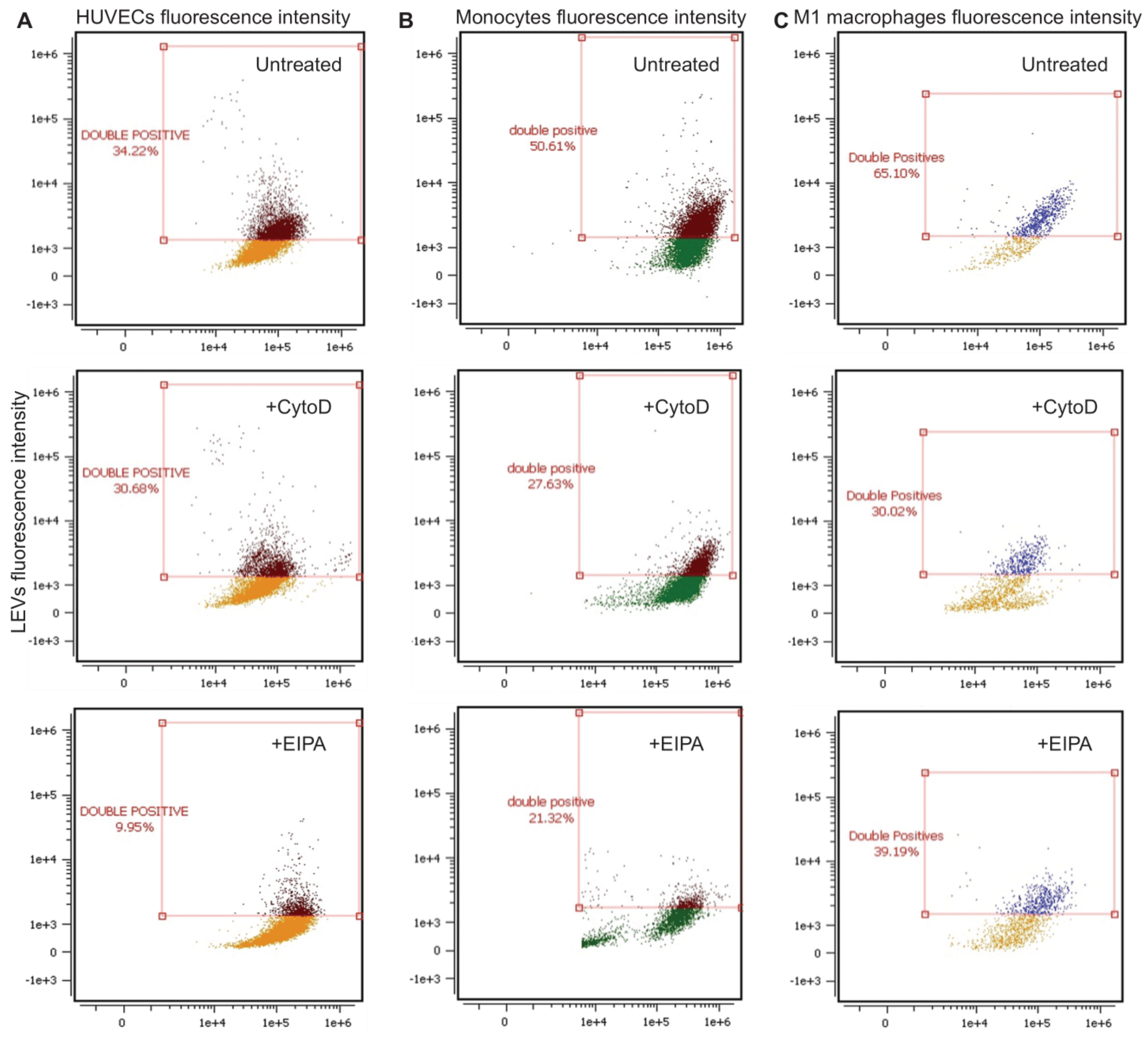


**Fig. S18. Internalization of tumor-cell derived LEVs by HUVECs, monocytes and M1 macrophages.** (**A-C**) All data were obtained by flow cytometry. Dot plots of double positive events of cells (x axis) that have internalized LEVs (y axis) after 16 hr co-culture (top) and/or pre-treatment with Cyto-D (middle) or EIPA (bottom), for HUVECs (**A**), monocytes (**B**) and M1 macrophages (**C**). Cells were stained with CMTPX cell tracker (x axis fluorescence intensity) and LEVs were stained with CMFDA cell tracker (y axis fluorescence intensity).

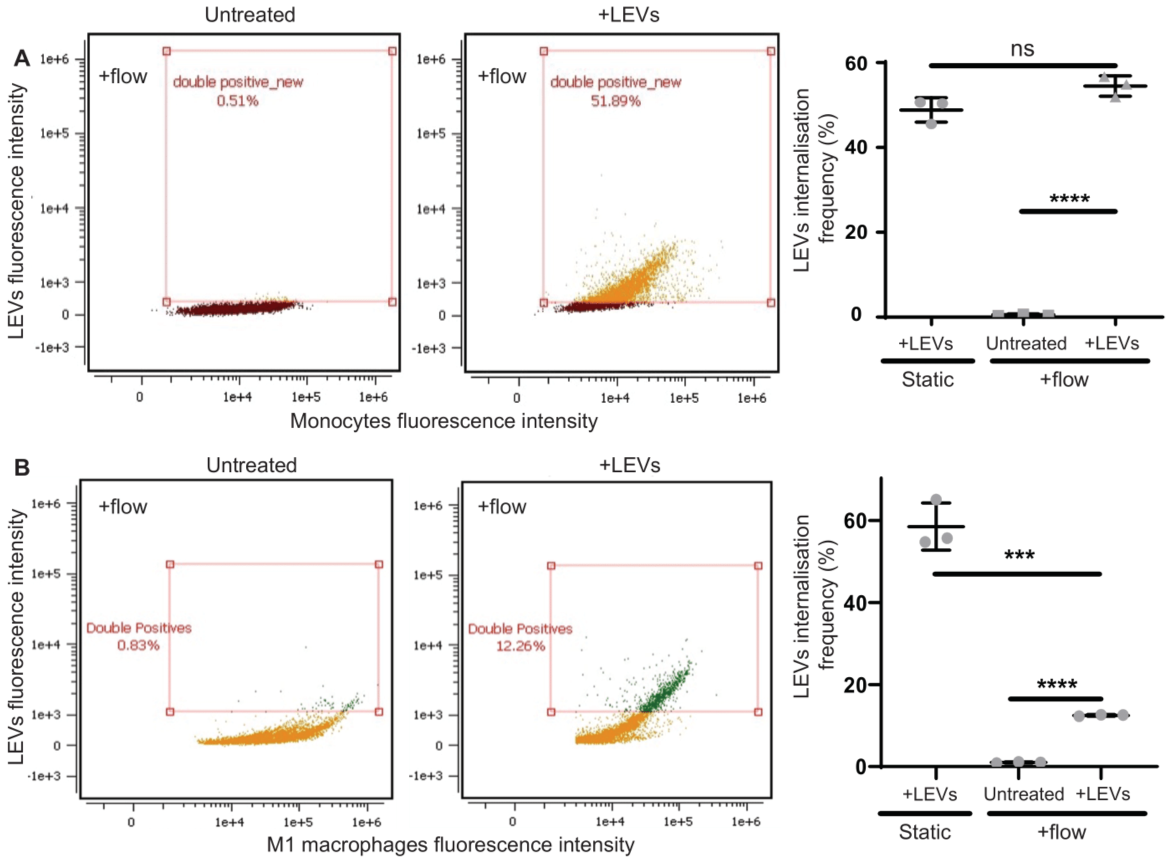


**Fig. S19. Internalization of tumor-cell derived LEVs by monocytes and M1 macrophages under flow conditions.** (**A**) Dot plots of untreated (left) or LEV-treated monocytes (middle) & quantification (right) of internalized LEVs by monocytes under flow conditions (n=3). (**B**) Dot plots of untreated (left) or LEV-treated M1 macrophages (middle) & quantification (right) of internalized LEVs by M1 macrophages under flow conditions (n=3). Cells were stained with CMTPX cell tracker (x axis fluorescence intensity) and LEVs were stained with CMFDA cell tracker (y axis fluorescence intensity).

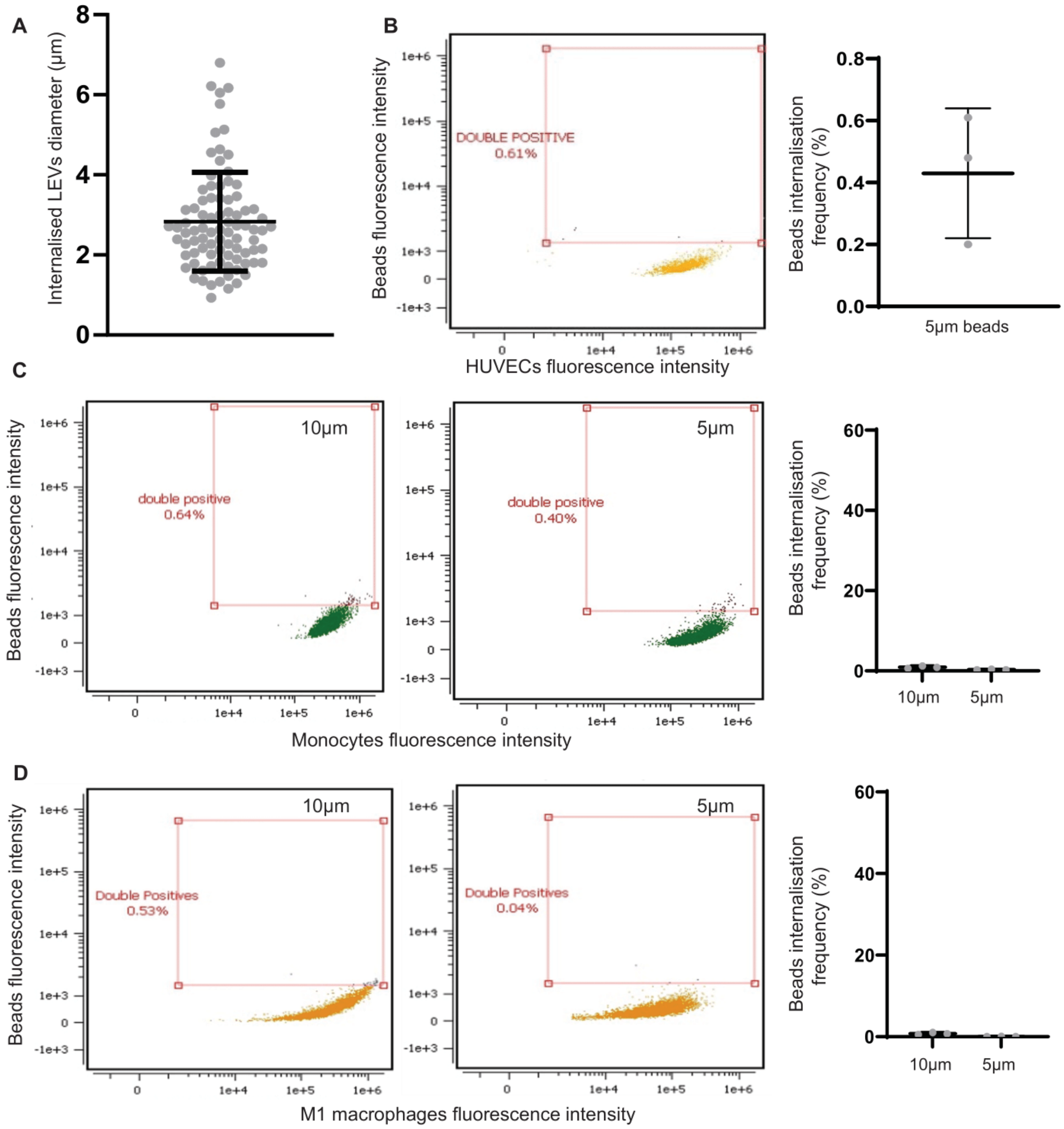


**Fig. S20. Investigation of fluorescent beads internalization by HUVECs, monocytes and M1 macrophages.** (**A**) Diameter (µm) of internalized LEVs by HUVECs (n= 87 LEVs), verified by confocal microscopy. (**B**) Dot plot (left) & quantification (right) of 5 µm beads internalized by HUVECs (n=3). (**C**) Dot plots of 10 µm beads (left) and 5 µm beads (middle) & quantification (right) of their internalization by monocytes (n=3). (**D**) Dot plots of 10 µm beads (left) and 5 µm beads (middle) & quantification (right) of their internalization by M1 macrophages (n=3). All data excluding (A) were obtained from flow cytometry analysis. Cells were stained with CMTPX cell tracker (x axis fluorescence intensity) and fluorescent beads were purchased and used (y axis fluorescence intensity).

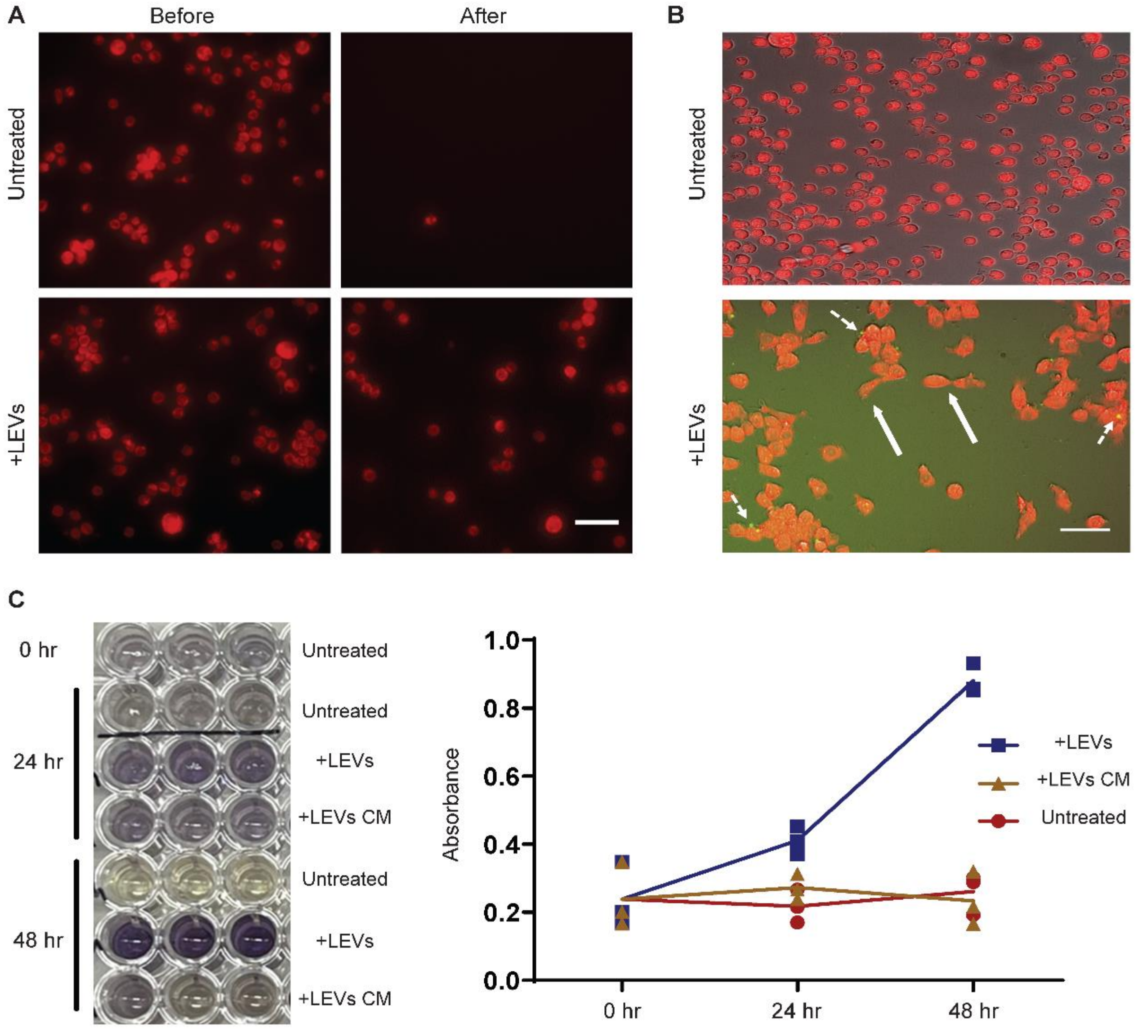


**Fig. S21. Impact of tumor cell-derived LEVs on monocytes.** (**A**) Images of fluorescently tagged monocytes with CMTPX cell tracker (red) (untreated-top or LEV-treated-bottom) that remained adhered before (left) and after (right) cell media removal. Scale bar: 50 µm. (**B**) Images of fluorescently tagged untreated (top) or LEV treated monocytes (bottom), stained with CMTPX cell tracker (red) after 30hr co-culture. LEVs-were pre-stained with CMFDA cell tracker (green). Dashed arrows indicate locations that LEVs are in proximity to monocytes. Full arrows indicate monocytes that have assumed stretched morphologies. Scale bar: 50 µm. (**C**) Absorbance assay (left) & quantification (right) of proliferation rates at 0, 24 and 48 hr for untreated monocytes, monocytes that were treated with LEVs or monocytes treated with CM from LEVs (n=3).

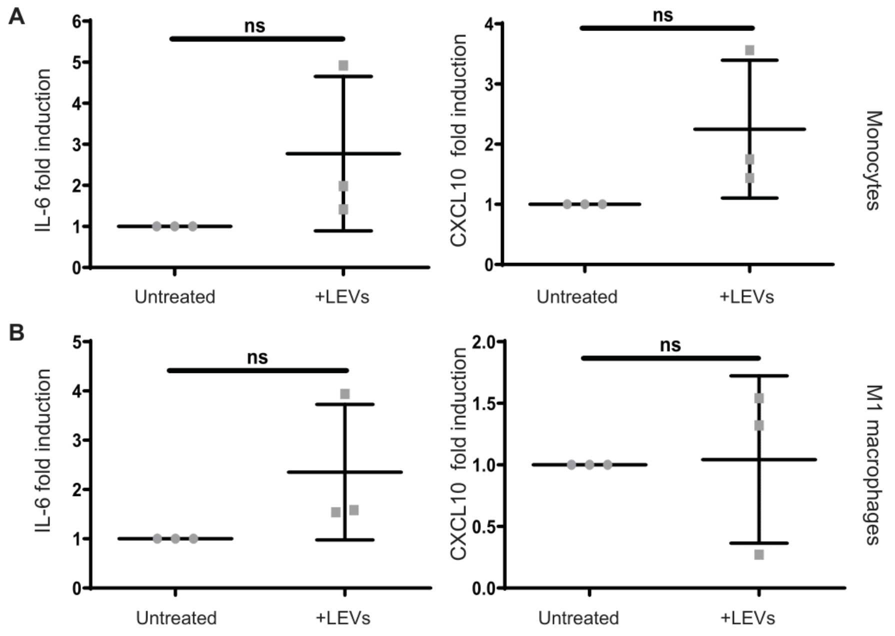


**Fig. S22. Investigation of tumor-cell derived LEVs on monocytes gene expression.** (**A**) Reverse transcription polymerase chain reaction of untreated or LEV-treated monocytes (30hr) for IL-6 (left) and CXCL10 (right) genes (n=3). (**B**) Reverse transcription polymerase chain reaction of untreated or LEV-treated M1 macrophages (30hr) for IL-6 (left) and CXCL10 (right) genes (n=3).

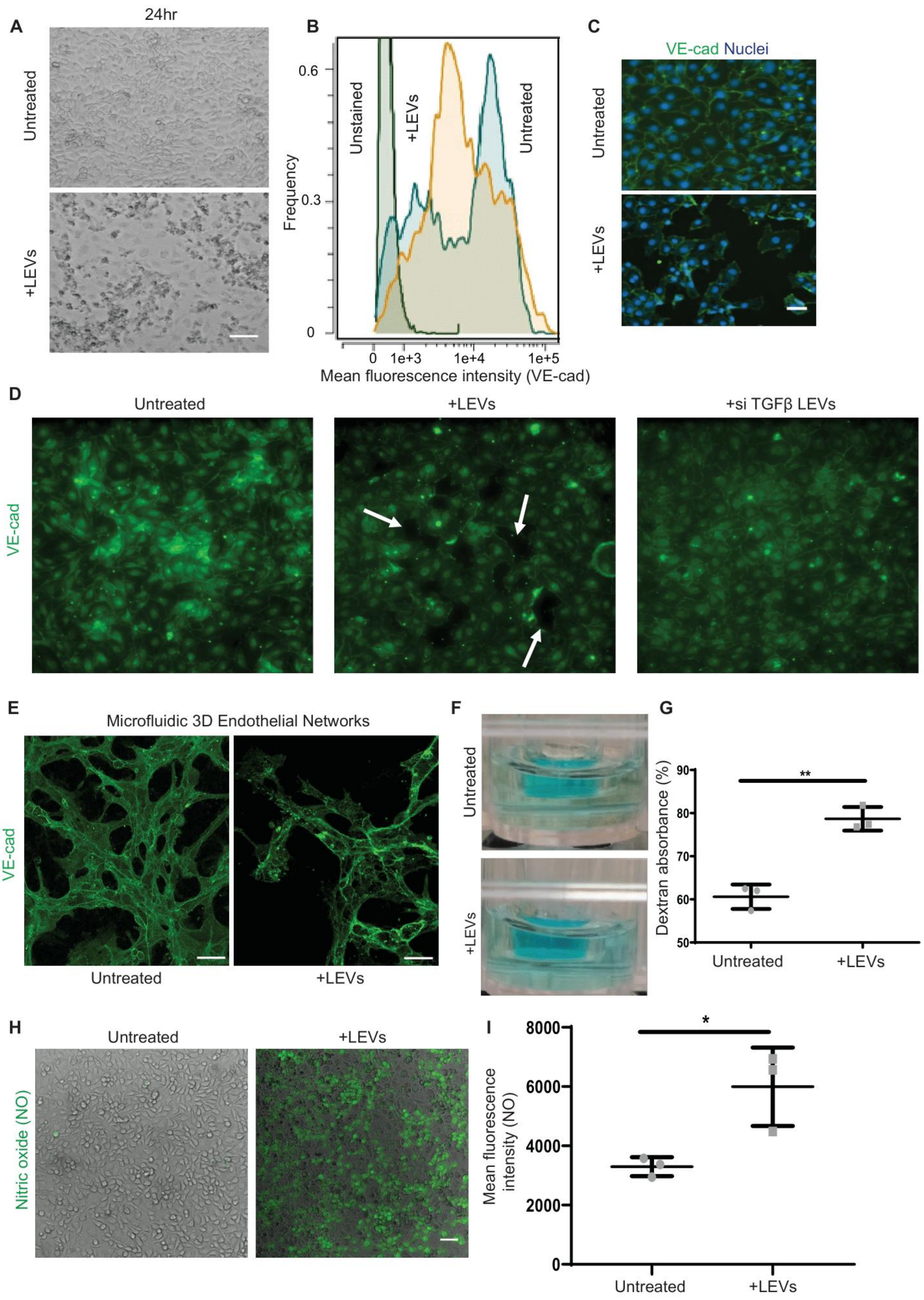

A
24hr
Untreated
+LEVs
B
0.6
0.3
0
Frequency
Unstained
+LEVs
Untreated
0 1e+3 1e+4 1e+5
Mean fluorescence intensity (VE-cad)
C
VE-cad Nuclei
Untreated
+LEVs
D
Untreated
+LEVs
+si TGFβ LEVs
VE-cad
E
Microfluidic 3D Endothelial Networks
VE-cad
Untreated
+LEVs
F
Untreated
+LEVs
G
**
Dextran absorbance (%)
90
80
70
60
50
Untreated
+LEVs
H
Untreated
+LEVs
Nitric oxide (NO)
I
*
8000
6000
4000
2000
0
Mean fluorescence intensity (NO)
Untreated
+LEVs

**Fig. S23. Tumor cell-derived LEVs reduce VE-cad expression on HUVECs and increase dextran permeability on HUVEC monolayers.** (**A**) Untreated or LEV-treated 2D HUVEC monolayers at 24hrs. Scale bar 50 µm. (**B**) Histogram of VE-cad mean fluorescence intensity of 2D HUVECs, analyzed via flow cytometry (n=3) (**C**) Multifluorescent images of untreated or LEV-treated 2D HUVEC monolayers (30hr). HUVECs were stained for VE-cad (green) and nuclei with Hoechst-33342 (blue). Scale bar 50 µm. (**D**) VE-cad staining of untreated, LEV-treated or siTGFβ LEV-treated 2D HUVEC monolayers. White arrows indicate gaps in the monolayer. (**E**) Fluorescent staining of VE-cadherin in untreated or LEV-treated microfluidic 3D endothelial networks. Scale bar: 100 µm. (**F-G**) Images (left) of untreated (top) or LEV-pre-treated (bottom) 2D HUVEC monolayers grown in transwell inserts (**F**) & quantification (**G**) of absorbance (right) of permeated dextran, collected from the media (n=3). (**H-I**) Intracellular nitric oxide (NO) (green) live staining (**H**) & quantification (**I**) of untreated and LEV-treated 2D HUVEC monolayers (n=3). Scale bar 50 µm.

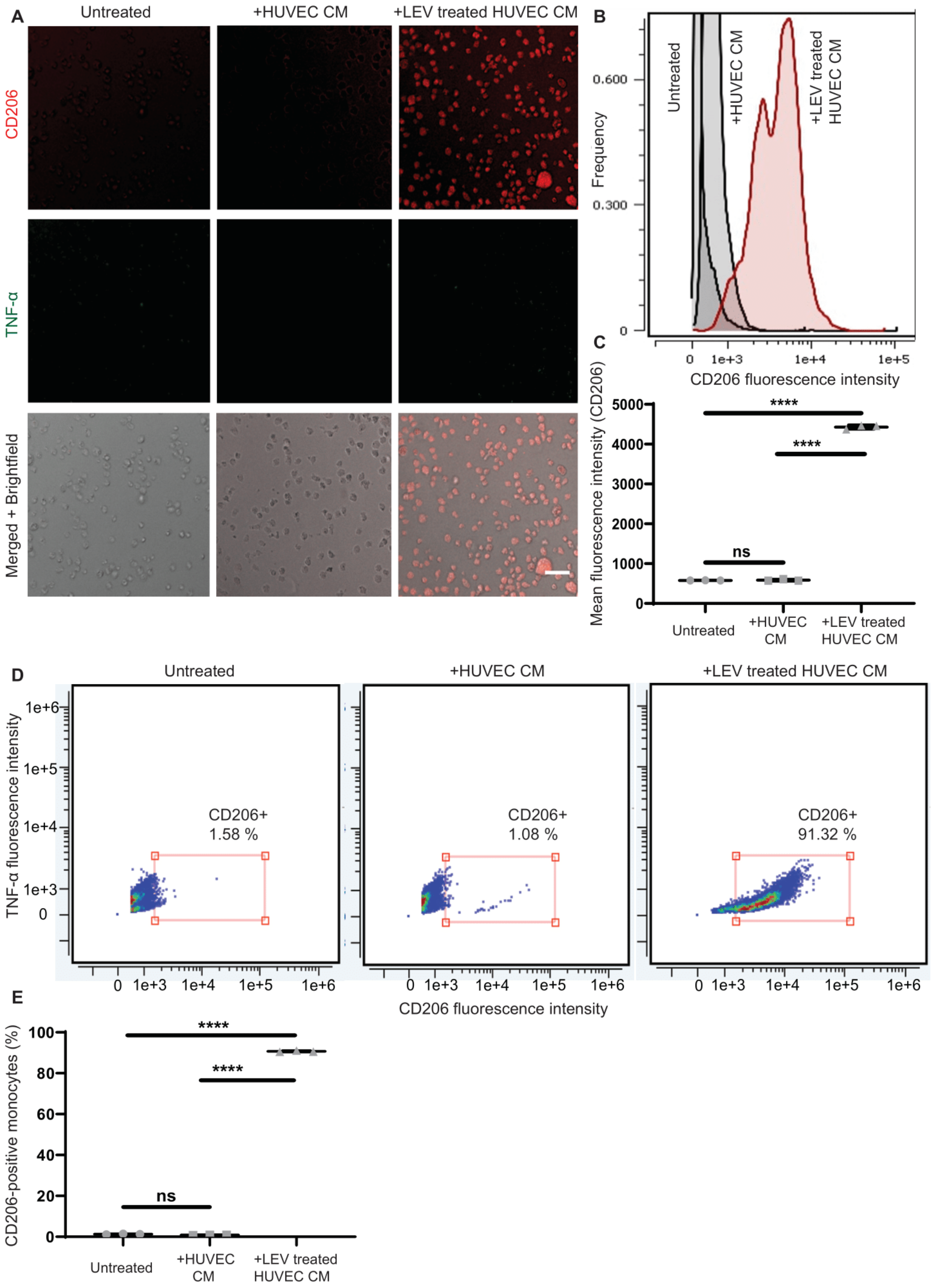
A
Untreated
+HUVEC CM
+LEV treated HUVEC CM
CD206
TNF-α
Merged + Brightfield
B
Frequency
0.600
0.300
0
Untreated
+HUVEC CM
+LEV treated HUVEC CM
0
1e+3
1e+4
1e+5
CD206 fluorescence intensity
C
Mean fluorescence intensity (CD206)
5000
4000
3000
2000
1000
0
****
****
ns
Untreated
+HUVEC CM
+LEV treated HUVEC CM
D
Untreated
+HUVEC CM
+LEV treated HUVEC CM
TNF-α fluorescence intensity
1e+6
1e+5
1e+4
1e+3
0
CD206+
1.58 %
CD206+
1.08 %
CD206+
91.32 %
0 1e+3 1e+4 1e+5 1e+6
CD206 fluorescence intensity
E
CD206-positive monocytes (%)
100
80
60
40
20
0
****
****
ns
Untreated
+HUVEC CM
+LEV treated HUVEC CM

**Fig. S24. CM from LEV-pre-treated HUVECs polarizes monocytes to M2 macrophages.** (**A**) Multifluorescent images of untreated monocytes (left) or monocytes that were treated either with CM from untreated HUVECs (middle) or CM from LEV-pre-treated HUVECs (right). Monocytes were stained for CD206 (red) and TNF-α (green). Scale bar: 50 µm. (**B-C**) Histogram (**B**) & quantification (**C**) of CD206 mean fluorescence intensity for untreated monocytes, monocytes treated with CM from untreated HUVECs and monocytes treated with CM from LEV-pre-treated HUVECs, analyzed via flow cytometry (n=3). (**D-E**) Dot plots (**D**) & quantification (**E**) of CD206 positive events for untreated monocytes (left), monocytes treated with CM from untreated HUVECs (middle) and monocytes treated with CM from LEV-pre-treated HUVECs (right), analyzed via flow cytometry (n=3). CD206 and TNF-α fluorescence intensity are represented in x and y axes, respectively.

**Table S1.**

Microfluidic capillary designs. Design parameters: a) $d_0$, $d_1$ and $d_2$ represent effective diameter of parent and two daughter bifurcation channels, b) $\theta^o$ represents the angle between daughter bifurcation channels, c) Symmetry between two daughter channels represents equal angle deviation from the parent channel. Non-bifurcated devices represent a straight channel without branching. Non-constricted devices represent large channels that do not constrict cells.

| Name | $d_0$ (µm) | $d_1$ (µm) | $d_2$ (µm) | Angle ($\theta^o$) | Symmetry |
|---|---|---|---|---|---|
| EWS | 10 | 6.3 | 6.3 | 87$^o$ | YES |
| EWA | 10 | 6.3 | 6.3 | 87$^o$ | NO |
| UWS | 10 | 5.5 | 7 | 87$^o$ | YES |
| UWA | 10 | 5.5 | 7 | 87$^o$ | NO |
| ENS | 10 | 6.3 | 6.3 | 43.5$^o$ | YES |
| ENA | 10 | 6.3 | 6.3 | 43.5$^o$ | NO |
| UNS | 10 | 5.5 | 7 | 43.5$^o$ | YES |
| UNA | 10 | 5.5 | 7 | 43.5$^o$ | NO |
| ENA_5/5 | 7.9 | 5 | 5 | 43.5$^o$ | NO |
| ENA_7/7 | 11.3 | 7 | 7 | 43.5$^o$ | NO |
| ENA_9/9 | 14.7 | 9 | 9 | 43.5$^o$ | NO |
| UNA_5/7 | 9.6 | 5 | 7 | 43.5$^o$ | NO |
| UNA_5/9 | 11.4 | 5 | 9 | 43.5$^o$ | NO |
| Non-bifurcated | 7 | N/A | N/A | N/A | N/A |
| Non-constricted | 50 | 20 | 20 | 87$^o$ | YES |

**Table S2.**

Proteins identified in MDA-MB 231 cell-derived LEVs, via mass spectrometry based proteomic profiling, using LC-MS DIA analysis.

See attached excel file *Table S2*.

**Table S3.**

MISEV Category 1 transmembrane proteins identified in MDA-MB 231 cell-derived LEVs, via mass spectrometry based proteomic profiling.

| Transmembrane protein | MISEV Category 1 |
|---|---|
| HLA-A | Non-tissue specific (1A) |
| ADAM10 | Non-tissue specific (1A) |
| BSG | Non-tissue specific (1A) |
| CD47 | Non-tissue specific (1A) |
| CD55 | Non-tissue specific (1A) |
| CD59 | Non-tissue specific (1A) |
| CD81 | Non-tissue specific (1A) |
| CD82 | Non-tissue specific (1A) |
| GNA11 | Non-tissue specific (1A) |
| GNA13 | Non-tissue specific (1A) |
| GNAI1 | Non-tissue specific (1A) |
| GNAI2 | Non-tissue specific (1A) |
| GNAI3 | Non-tissue specific (1A) |
| GNAO1 | Non-tissue specific (1A) |
| GNAQ | Non-tissue specific (1A) |
| ITGA1 | Non-tissue specific (1A) |
| ITGA2 | Non-tissue specific (1A) |
| ITGA3 | Non-tissue specific (1A) |
| ITGA5 | Non-tissue specific (1A) |
| ITGA6 | Non-tissue specific (1A) |
| ITGB1 | Non-tissue specific (1A) |
| ITGB3 | Non-tissue specific (1A) |
| ITGB4 | Non-tissue specific (1A) |
| ITGB5 | Non-tissue specific (1A) |
| LAMP1 | Non-tissue specific (1A) |
| LAMP2 | Non-tissue specific (1A) |
| NT5E | Non-tissue specific (1A) |
| ABCC1 | Cell/tissue specific (1B) |
| APP | Cell/tissue specific (1B) |
| CD9 | Cell/tissue specific (1B) |
| EPCAM | Cell/tissue specific (1B) |
| ERBB2 | Cell/tissue specific (1B) |
| HLA-DRA | Cell/tissue specific (1B) |

**Table S4.**

MISEV Category 2 cytosolic proteins identified in MDA-MB 231 cell-derived LEVs, via mass spectrometry based proteomic profiling.

| Transmembrane protein | MISEV Category 2 |
|---|---|
| ARF6 | Proteins with lipid or protein binding ability (2A) |
| FLOT1 | Proteins with lipid or protein binding ability (2A) |
| FLOT2 | Proteins with lipid or protein binding ability (2A) |
| PDCD6IP | Proteins with lipid or protein binding ability (2A) |
| RHOA | Proteins with lipid or protein binding ability (2A) |
| SDCBP | Proteins with lipid or protein binding ability (2A) |
| TSG101 | Proteins with lipid or protein binding ability (2A) |
| VPS4A | Proteins with lipid or protein binding ability (2A) |
| VPS4B | Proteins with lipid or protein binding ability (2A) |
| HSP90AA1 | Proteins with lipid or protein binding ability (2A) |
| HSP90AB1 | Proteins with lipid or protein binding ability (2A) |
| HSP90B1 | Proteins with lipid or protein binding ability (2A) |
| HSPA13 | Proteins with lipid or protein binding ability (2A) |
| HSPA14 | Proteins with lipid or protein binding ability (2A) |
| HSPA4 | Proteins with lipid or protein binding ability (2A) |
| HSPA4L | Proteins with lipid or protein binding ability (2A) |
| HSPA5 | Proteins with lipid or protein binding ability (2A) |
| HSPA8 | Proteins with lipid or protein binding ability (2A) |
| HSPA9 | Proteins with lipid or protein binding ability (2A) |
| HSPB1 | Proteins with lipid or protein binding ability (2A) |
| HSPB11 | Proteins with lipid or protein binding ability (2A) |
| HSPBP1 | Proteins with lipid or protein binding ability (2A) |
| HSPD1 | Proteins with lipid or protein binding ability (2A) |
| HSPE1 | Proteins with lipid or protein binding ability (2A) |
| HSPG2 | Proteins with lipid or protein binding ability (2A) |
| HSPH1 | Proteins with lipid or protein binding ability (2A) |
| GAPDH | Promiscuous incorporation in EVs (2B) |
| ACTBL2 | Promiscuous incorporation in EVs (2B) |
| ACTL6A | Promiscuous incorporation in EVs (2B) |
| ACTN1 | Promiscuous incorporation in EVs (2B) |
| ACTN4 | Promiscuous incorporation in EVs (2B) |
| ACTR10 | Promiscuous incorporation in EVs (2B) |
| ACTR1A | Promiscuous incorporation in EVs (2B) |
| ACTR1B | Promiscuous incorporation in EVs (2B) |
| ACTR2 | Promiscuous incorporation in EVs (2B) |
| ACTR3 | Promiscuous incorporation in EVs (2B) |
| TUBA1C | Promiscuous incorporation in EVs (2B) |
| TUBA4A | Promiscuous incorporation in EVs (2B) |
| TUBB | Promiscuous incorporation in EVs (2B) |
| TUBB1 | Promiscuous incorporation in EVs (2B) |
| TUBB2A | Promiscuous incorporation in EVs (2B) |
| TUBB2B | Promiscuous incorporation in EVs (2B) |
| TUBB3 | Promiscuous incorporation in EVs (2B) |

| | |
|---|---|
| TUBB4A | Promiscuous incorporation in EVs (2B) |
| TUBB4B | Promiscuous incorporation in EVs (2B) |
| TUBB6 | Promiscuous incorporation in EVs (2B) |
| TUBG1 | Promiscuous incorporation in EVs (2B) |
| TUBGCP2 | Promiscuous incorporation in EVs (2B) |
| TUBGCP3 | Promiscuous incorporation in EVs (2B) |
| TUBGCP5 | Promiscuous incorporation in EVs (2B) |
| TUBGCP6 | Promiscuous incorporation in EVs (2B) |

**Table S5**

Proteins annotated for protein secretion identified in MDA-MB 231 cell-derived LEVs, via mass spectrometry based proteomic profiling.

| Protein list |
|---|
| ARF1 |
| ADAM10 |
| ANP32E |
| AP1G1 |
| AP2B1 |
| AP2M1 |
| AP2S1 |
| AP3B1 |
| AP3S1 |
| ARCN1 |
| ARFGAP3 |
| ARFGEF2 |
| ARFIP1 |
| ATP1A1 |
| ATP6V1H |
| CLTA |
| CLTC |
| COPB1 |
| COPB2 |
| COPE |
| CTSC |
| DNM1L |
| DST |
| EGFR |
| ERGIC3 |
| GBF1 |
| IGF2R |
| KIF1B |
| KRT18 |
| LAMP2 |
| LMAN1 |
| M6PR |
| MAPK1 |
| NAPA |
| NAPG |
| OCRL |
| PPT1 |
| RAB14 |
| RAB2A |
| RAB5A |
| RAB9A |
| RPS6KA3 |
| SCAMP1 |
| SCAMP3 |

| |
|---|
| SCRN1 |
| SEC22B |
| SEC24D |
| SEC31A |
| SNAP23 |
| SNX2 |
| SOD1 |
| STAM |
| STX12 |
| STX7 |
| TMED10 |
| TMED2 |
| TMX1 |
| TPD52 |
| TSG101 |
| USO1 |
| VAMP3 |
| VPS45 |
| VPS4B |
| YIPF6 |
| YKT6 |
| ZW10 |

**Table S6.**

List of analyzed proteins via immunocytochemistry (including acronyms), on cells or LEVs, and primary/secondary antibodies used (with concentrations).

| Protein | Cell | Primary antibody (final concentration) | Secondary antibody (final concentration) |
|---|---|---|---|
| Cytokeratin 18 (CK18) | MDA-MB 231 cells or LEVs from MDA-MB 231 | Primary rabbit anti human monoclonal antibody CK18 (1 µg/ml) | Goat anti rabbit AlexaFluor 488 (2 µg/ml) |
| Cluster of differentiation (CD81) | MDA-MB 231 cells or LEVs from MDA-MB 231 | Primary rabbit anti human monoclonal antibody CD81 (0.66 µg/ml) | Goat anti rabbit AlexaFluor 488 (2 µg/ml) |
| Heat shock protein 90 (HSP90) | MDA-MB 231 cells or LEVs from MDA-MB 231 | Primary rabbit anti human monoclonal antibody HSP90 (1.74 µg/ml) | Goat anti rabbit AlexaFluor 488 (2 µg/ml) |
| Filamentous actin (F-actin) | MDA-MB 231 cells | Phalloidin 647 (1000 time dilution from stock vial-Abcam ab176759) | N/A |
| Vascular endothelial cadherin (VE-cad) | HUVECs | Primary rabbit anti human monoclonal antibody VE-cad (1 µg/ml) | Goat anti rabbit AlexaFluor 488 (2 µg/ml) |
| Vascular cell adhesion molecule (VCAM) | HUVECs | Primary rabbit anti human monoclonal antibody VCAM (2 µg/ml) | Goat anti rabbit AlexaFluor 488 (2 µg/ml) |
| Tumor necrosis factor α (TNF-α) | Monocytes | Primary rabbit anti human monoclonal antibody TNF-α (0.09 µg/ml) | Goat anti rabbit AlexaFluor 488 (2 µg/ml) |

| | | | |
|---|---|---|---|
| Cluster of differentiation (CD206) | Monocytes | Primary rabbit anti human monoclonal antibody PE/Cy7® Anti-Mannose Receptor (100 times dilution) | Goat anti rabbit AlexaFluor 647 (2 µg/ml) |
| C-X-C motif chemokine (CXCL10) | Monocytes | Primary rabbit anti human monoclonal antibody CXCL10 (2.5 µg/ml) | Goat anti rabbit AlexaFluor 488 (2 µg/ml) |

**Table S7.**

List of excitation and emission values used for dot plots generation via flow cytometry.

| Type | x axis | y axis |
|---|---|---|
| MDA-MB 231 cells | Hoechst fluorescence intensity (Ex/Em: 405/456) | CK18 or CD81 or HSP90 fluorescence intensity (Ex/Em: 488/528) |
| LEVs | Red cell tracker fluorescence intensity (Ex/Em: 561/611) | CK18 or CD81 or HSP90 fluorescence intensity (Ex/Em: 488/528) |
| HUVECs | N/A | VE-cad fluorescence intensity (Ex/Em: 488/528) |
| Monocytes | CD206 fluorescence intensity (Ex/Em: 561/611) | TNF-a fluorescence intensity (Ex/Em: 488/528) |

**Table S8.**

List of mRNA molecules (with acronyms), including forward and reverse sequence (primers).

| Gene name | Forward primer | Reverse primer |
|---|---|---|
| Interleukin 6 (IL-6) | GGTCCAGTTGCCTTCTCCCTG | TGCCCATGCTACATTTGCCG |
| C-X-C motif chemokine 10 (CXCL10) | TGAGCCTACAGCAGAGGAACC | GCCTCTGTGTGGTCCATCCTT |
| Fibronectin (FN) | GATGCACCATCCAACCTGCG | GATTGAGTCCCGGACCGTGT |
| Aldehyde dehydrogenase 1 (ALDH1) | CCCATCACAGGAGAGAAC | CCCTTAAATGCCATCCTG |
| B-cell lymphoma 2 (BCL2) | CAAGAAACTTCTACGACAGC | AAGCCATTTTCCTCTTCTTG |
| Cluster of differentiation (CD44) | TTATCAGGAGACCAAGACAC | ATCAGCCATTCTGGAATTTG |
| Vimentin (Vim) | GGAAACTAATCTGGATTCACTC | CATCTCTAGTTTCAACCGTC |
| Ribosomal protein large P (RPLO) | CGGTTTCTGATTGGCTAC | ACGATGTCACTTCCACG |
| Cluster of differentiation (CD206) | CGTTCGGTTCACCCACTGGA | ACATCCCATAAGCCCCCTGC |
| Arginase 1 (ARG-1) | TGGCAAGGTGGCAGAAGTCA | TGGCATGGCCAGAGATGCTT |
| Vascular endothelial cadherin (VE-cad) | GCCCACAGGCACGATCTGTT | CATCCGGTTCTGGGGCTCAT |

**Movie S1. MDA-MB 231 cancer cell sheds LEVs in capillary bifurcation.**

Brightfield video of MDA-MB 231 cells shedding LEVs in capillary bifurcation variant UNA. White arrows indicate regions of cells shedding.

**Movie S2. Fluorescently-tagged MDA-MB 231 cancer cell sheds LEVs in capillary bifurcation.**

A MDA-MB 231 cell shedding LEVs in capillary bifurcation variant UNA. Cytoplasm was stained with CMFDA cell tracker (green) and nuclei with Hoechst-33342 (blue).

**Movie S3. CDX19 explant cell sheds a LEV in capillary bifurcation.**

A CTC-derived CDX19 cell shedding in capillary bifurcation variant UNA_5/9 a single LEV. Cytoplasm was stained with CMFDA cell tracker (green).

**Movie S4. Patient CTC sheds a LEV in capillary bifurcation.**

Patient CTC (isolated from small cell lung carcinoma patient) shedding in capillary bifurcation variant UNA_5/9 a single LEV. Cytoplasm was stained with CMFDA cell tracker (green) and nuclei with Hoechst-33342 (blue).

**Movie S5. Large cluster sheds multiple LEVs in capillary bifurcation.**

A multicellular MDA-MB 231 cell cluster shedding in capillary bifurcation variant UNA_5/9 multiple LEVs. Cytoplasm was stained with CMFDA cell tracker (green) and nuclei with Hoechst-33342 (blue).

**Movie S6. Non-shedding cancer cell transits in a capillary bifurcation.**

A non-shedding MDA-MB 231 cell transiting in capillary bifurcation variant ENA. Cytoplasm was stained with CMFDA cell tracker (green).

**Movie S7. Cancer cell undergoes lysis post-shedding in a capillary bifurcation.**

A MDA-MB 231 cell shedding in capillary bifurcation variant EWA and undergoing lysis. Cytoplasm was stained with Calcein-AM (green) and nuclei with Hoechst-33342 (blue).

**Movie S8. Co-culture of endothelial cells with LEVs.**

A human umbilical vein endothelial (HUVEC) cell near LEVs. HUVECs were stained with CMFDA cell tracker (green) and LEVs with CMTPX cell tracker (red).
